# Sex-specific Effects of Outdoor Air Pollution on Subcortical Microstructure and Weight Gain: Findings from the ABCD Study®

**DOI:** 10.1101/2025.06.04.657916

**Authors:** Miguel Ángel Rivas-Fernández, Jonatan Ottino-González, Sevan Esaian, Victoria E. Goldman, Megan. M. Herting, Tanya L. Alderete, Shana Adise

## Abstract

Obesity is associated with structural alterations of brain regions that support eating behavior. Exposure to air pollutants might exacerbate this association through neurotoxic effects on the brain. This study evaluated whether air pollution exposure 9-10 years old children, coupled with brain microstructure development in appetite-regulating regions, is associated with body mass index (BMI) changes over two years, and whether these associations differ by sex.

Data were gathered from the Adolescent Brain Cognitive Development Study® (n_baseline_=4,802, ages=9-10, males=49.9%, n_follow-up_=2,439, ages=11-12, males=51.1%). Annual average estimates of ambient fine particulate matter (PM2.5), nitrogen dioxide (NO2), ground-level ozone (O3), and redox-weighted oxidative capacity (Oxwt, a joint measure of NO2 and O3) were gathered from youth’s residential addresses. Brain microstructure in 16 subcortical regions was assessed using diffusion-weighted MRI, focusing on proxies of cellular and neurite density: restricted normalized isotropic (RNI) and directional (RND) diffusion, respectively. Linear mixed-effects models examined whether air pollution and brain microstructure are related to BMI changes over two years, and whether these associations differed by sex.

Exposure to PM2.5 coupled with high RND estimates in right caudate nucleus, bilateral putamen, and pallidum were associated with higher BMI over time, with pronounced effects in males (all *p*<0.05).

PM2.5 coupled with greater neurite density in regions involved in reward-processing and decision-making were associated with higher BMI over a 2-year follow-up, especially in males. This research highlights air pollution as a modifiable risk factor for how differences in basal ganglia neurite density map onto obesity risk, with important implications for public health policy.

**HIGHLIGHTS:**

- High PM2.5 exposure and subcortical neurite density is associated with weight gain
- PM2.5 and subcortical development associations with BMI are pronounced in males
- Significant associations with BMI were found in regions involved in food intake

## 1. INTRODUCTION

Childhood obesity represents a major public health challenge, and prevalence rates are increasing (Ng et al., 2024). Converging evidence suggests that this complex disease is associated with structural and functional variations in brain regions that regulate appetite (Gómez-Apo et al., 2021; Raji et al., 2010; Ronan et al., 2020). Thus, altered structure and function within these regions may contribute to altered eating behaviors and subsequent weight gain. Recent work has showed that alterations within these appetite-regulating structures may predate weight gain (Adise, Ottino-Gonzalez, et al., 2024), implying altered development. However, the factors contributing to altered brain development are not well understood, especially within the context of obesity. Environmental air pollutants have emerged as a potential contributor to altered brain development (Morrel et al., 2025) via negative effects of neuroinflammation (Costa et al., 2019), and they have also been associated with increased obesity rates in children (Zheng et al., 2024). While there appears to be a relationship between altered brain development, air pollution, and obesity, no studies have examined these in tandem. Understanding these relationships is important given that childhood obesity rates are continuing to rise despite efforts to curtail disease progression (Ng et al., 2024). This study can provide insights into the contributing role of air pollution on childhood obesity. This knowledge is important for informing public health policies aimed at reducing the harmful effects of air pollution and/or weight gain on brain health.

Outdoor air pollution is emitted from different sources like manufacturing, transportation, and energy production. While there are many air pollutants, three criteria pollutants have been identified as being particularly harmful to health (Wang et al., 2025), including fine inhalable particulate matter with a diameter of 2.5 micrometers or smaller (PM2.5), nitrogen dioxide (NO2), and ground-level ozone (O3). These pollutants have been previously identified as leading contributors to pediatric health complications, including respiratory disease (Garcia et al., 2021), obesity (Zheng et al., 2024), altered brain volume and cortical thickness (Morrel et al., 2025), and differences in executive functions (Lopuszanska & Samardakiewicz, 2020; Thompson et al., 2023). Importantly, air pollution exposure during childhood and adolescence may be more detrimental to health than at later lifespan stages (Harr et al., 2022; Shi et al., 2022). This is because the body and brain are still developing during the transition from childhood to adolescence (Vijayakumar et al., 2018), which is associated with substantial shifts in emotional and cognitive regulation (Luna, 2009; Yurgelun-Todd, 2007). Therefore, understanding whether exposure to certain environmental air pollutants may contribute to different health outcomes during critical developmental periods is needed.

While air pollution may impact overall health (Manisalidis et al., 2020), its potential influence on association between the brain and obesity risk is particularly complex. That is because evidence suggests that inhalation of toxic chemicals like air pollution may trigger neuroinflammation (Lucchini et al., 2012), which occurs when the brain’s immune response activates astrocytes and microglia to remove harmful pathogens (DiSabato et al., 2016). While acute neuroinflammation is not necessarily harmful, animal models have suggested that chronic neuroinflammation can erode the myelin sheath integrity of a neuron and ultimately cause macro- and microstructural changes that could lead to cell death (Costa et al., 2019). As such, air pollution may indirectly cause structural and functional changes to the brain via neuroinflammation (Morrel et al., 2025). However, no previous research has evaluated the role of air pollution as a moderating factor in brain-obesity phenotypes.

Recently, a novel neuroimaging technique, known as restricted spectrum imaging (RSI), has emerged as a tool to provide *in-vivo* correlates of gray matter tissue microstructure, which may better capture brain microstructural differences and provide insight into overall brain health. RSI is thought to provide insight examining intra- and extra-cellular changes at the microstructural level, which might represent an indirect proxy for tissue cytoarchitecture (White et al., 2013, 2014). In gray matter, RSI can be utilized to generate restricted (i.e. intracellular) normalized isotropic diffusion (RNI), which is suggestive of changes in cellular density (i.e., glia and cell bodies) and restricted (i.e. intracellular) normalized directional diffusion (RND), suggestive of neurite organization and density (i.e., axons and dendrites) (White et al., 2013, 2014), although more research is needed to confirm this. However, among adolescents RNI and RND differences have been linked with higher body mass index (BMI) (Li et al., 2023). Interestingly, cross-sectional and longitudinal changes in RNI and RND within both gray and white matter have also been associated with both increased exposure to outdoor PM2.5 (Bottenhorn et al., 2024; Burnor et al., 2021; Cotter et al., 2024; Sukumaran et al., 2023). Yet, no studies have examined if outdoor air pollution may moderate the association between RNI and RND differences in subcortical brain regions involved in food intake control, and changes in weight gain over time during the transition to adolescence.

Given this neuroimaging advancement, the current longitudinal study utilized RSI to evaluate whether air pollution exposure and brain microstructure development are associated with BMI changes from 9-12 years old among a large sample of youth enrolled in the Adolescent Brain Cognitive Development (ABCD) Study®. In the present study, we examined whether one year of annual average exposure to daily ambient PM2.5, NO2, O3 (8-hour maximum), and redox-weighted oxidative capacity (Oxwt—a combined measure of NO2 and O3) at ages 9–10 was associated with changes in BMI over a two-year follow-up period. We then examined whether exposure to air pollution coupled with brain microstructure development is associated with BMI changes over a two-year period (from age 9 to 11 or 10 to 12). Our questions of interest focused on understanding these associations, particularly among the subcortex, as many of these regions are involved in food intake (Kung et al., 2022). We hypothesized that air pollution exposure coupled with gray matter microstructural alterations (i.e., greater cellular density [RNI] and neurite density [RND]) would be associated with greater weight gain over time. Because studies have shown sex differences in the associations between brain structure and air pollution (Cotter et al., 2024; Peterson et al., 2022) and weight gain (Adise, Ottino-Gonzalez, et al., 2024), we also stratified the analyses by sex. In line with this previous research (Adise, Ottino-Gonzalez, et al., 2024; Cotter et al., 2024; Peterson et al., 2022), we expected to find sex differences in subcortical alterations associated with weight gain trajectories over time as well as to how air pollution may modify this association. This longitudinal study will advance our current understanding of how air pollution may couple with weight-based brain phenotypes to contribute to increased risk of overweight and obesity in children. These insights could have important implications for shaping public health strategies to mitigate the adverse effects of air pollution exposure and to prevent childhood obesity.

## 2. METHODS

### 2.1 Study design

The ABCD Study^®^ is a 21-site, 10-year cohort research study that began between 2016 and 2018, involving youth aged 9 to 10 years at the start. This large-scale research project involves annual examinations with biannual follow-up assessments. The ABCD Study^®^ recruitment strategy was designed to reflect the demographic characteristics from the American Community Survey in the United States. Detailed information about the study’s design, assessment and objectives can be found in numerous publications (www.ABCDStudy.org). The ABCD Study^®^ is responsible for obtaining approval from caregivers and assent from the youth involved in this project and all the protocols were approved by centralized Institutional Review Board at the University of California, San Diego. This manuscript focuses on baseline and year 2 follow-up assessments included in the 5.1 data release (https://dx.doi.org/10.15154/z563-zd24).

The ABCD Study had minimal exclusion criteria but to obtain a sample optimal for the current manuscript’s analyses, several exclusion criteria were applied (e.g., medical or neurological conditions, drug exposure during pregnancy, head injuries, extreme prematurity, or very low birth weight). Additional quality control assessment was also performed on anthropometric and MRI data. A detailed description of the exclusion criteria employed in the present study can be found in the **Supplementary Materials**.

### 2.2. Demographics

Demographic information about the youth was reported by their caregivers, including sex at birth and date of birth (used to calculate age at the time of the visit). Caregivers provided the youth’s race based on a 19-item response that was collapsed into six categories (American Indian/Native American and Native Hawaiian/Pacific Islander, Asian, Black, Mixed, Other, White), and ethnicity based on a binary response (Hispanic/Latinx, non-Hispanic/Latinx). Caregivers also reported the level of education attained in the household based on a 21-item response that was coded as follows: less than high school (<13 years), high-school graduate (13-14 years), some college (15-17 years), bachelor’s degree (18 years), and postgraduate degree (19-21 years). In the present study, the household education level was determined to the highest level attained by either caregiver.

### 2.3. Physical assessments

Height and weight were assessed by a trained researcher and converted to BMI (kg/m^2^). See the **Supplementary Materials** for more details. The CDC’s BMI percentiles (Kuczmarski et al., 2002), which consider sex, age, height, and weight, were used to classify youth into weight categories (e.g., underweight, healthy weight, overweight, obese) for descriptive purposes only. Statistical analyses were conducted using raw BMI values (see Adise, Rhee, et al., 2024 for justification). Additionally, the youth’s pubertal status was assessed using the Pubertal Development Scale (Petersen et al., 1988) (see further details in the **Supplemental Materials**). Handedness of participants was assessed using the Youth Edinburgh Handedness Short Form (Veale, 2014).

### 2.4. Magnetic Resonance Imaging acquisition and preprocessing

Magnetic Resonance Imaging (MRI) data were acquired with 3T MRI scanners across 21 sites (n _scanners_ = 31). Brain structural imaging data were collected through T1-weighted and multi-shell Diffusion Weighted Imaging (DWI) sequences. A detailed description of the preprocessing pipeline employed can be found in **Supplemental Materials**.

Complete information about imaging acquisition and processing protocols of the ABCD data are described in previous publications (Casey et al., 2018; Hagler et al., 2019). The present study focused on the following water diffusion metrics derived from the RSI model: RND and RNI diffusion. RND is a measure of directional intracellular, restricted diffusion, which occurs when the movement of water molecules is restricted by impermeable barriers. Regions with unidirectionally organized neurites have high RND. The RNI diffusion is a measure of isotropic intracellular restricted diffusion, which occurs when diffusion takes place in any direction. Increased estimates of this biomarker can be suggestive of changes in cellularity density (e.g., glial and cell bodies) (White et al., 2013). Each of these measures was normalized and defined as the Euclidean norm (square root of the sum of squares) for the corresponding model coefficients divided by the norm of all model coefficients. As such, these measures were unitless and ranged from 0 to 1, and the square of each measure was equivalent to the signal fraction for their respective model components (Hagler et al., 2019). This study evaluated the RND and RNI metrics in eight bilateral subcortical regions of interest (ROIs) with an important role in food intake behaviour (Kung et al., 2022): accumbens area, amygdala, caudate, hippocampus, thalamus, pallidum, putamen and ventral diencephalon. The automatic subcortical segmentation of these subcortical regions was based on an atlas containing probabilistic information on the location of these brain structures (Fischl et al., 2002).

### 2.5 Outdoor Air pollution estimates

Caregivers provided their residential addresses, which were geocoded to obtain estimates of outdoor air pollution. Annual average air pollution estimates were collected in 2016, corresponding to the start of the ABCD Study enrolment (i.e., when children were 9-10 years old). However, it should be noted that baseline data of the ABCD Study was collected between 2016 and 2018. Air pollution data was collected at a 1-km^2^ resolution using hybrid spatiotemporal models that combine satellite-based aerosol optical depth models, land-use regression, and chemical transport model outputs (Di et al., 2019, 2020; Requia, Coull, et al., 2020). Further information about the methods and protocols employed to estimate air pollution data in the ABCD Study^®^ are described elsewhere (Badilla et al., 2024; Fan et al., 2021).

The average annual air pollution concentrations were linked to the youth’s geocoded residential addresses. Daily exposure estimates were calculated across the U.S. for daily fine particulate matter (PM2.5) in micrograms per cubic meter (µg/m³), daily nitrogen dioxide (NO2) in parts per billion (ppb), and 8-hour maximum ozone (O3) levels in ppb, all at a 1 km² resolution (Di et al., 2019, 2020; Requia, Di, et al., 2020). Due to their potential to induce oxidative stress when combined, the redox-weighted oxidative capacity (Oxwt) was estimated as the ratio of the weighted redox potentials of NO2 and O3 using the following formula: [(1.07 * NO2) + (2.075 * O3)] / 3.145 (Williams et al., 2014).

Urbanicity of participant’s addresses was assessed with a categorical variable indicating if participants’ addresses were in census tracts considered to be urban (50,000 or more people), urban clusters (at least 2,500 and less than 50,000 people), or rural (less than 2,500 people per tract, and/or not included in urban areas or clusters). These categories were established according to the Census Tract Urban Classification and based on the 2010 census estimates. Given the high percentage of missing data for pollutants associated with the second and third addresses in the current sample (see further details in **Supplemental Materials**), this study focused exclusively on the primary residential address.

### 2.6 Statistical analyses

Statistical analyses were performed using R Studio (Team, 2016) (version 2024.4.2.764). Skewness and kurtosis were assessed for each numerical variable prior to any statistical analysis using the *e1071* R package (v. 1.7.16). Similarly, multicollinearity was evaluated by means of the variance inflation factor, using the *car* R package (Fox & Weisberg, 2019). Scripts employed to prepare data, perform the analysis, and plot results of the current manuscript are available elsewhere (https://github.com/Adise-lab/abcd-air-pollution-bmi).

#### 2.6.1 Associations between air pollutants and BMI

Given that positive associations between air pollution and body weight have not been reported in some previous research (Fioravanti et al., 2018), we first examined the relationship between BMI and each air pollutant (PM2.5, NO2, O3, and Oxwt) both across sexes and separately in males and females. We used linear mixed-effects models (LMEs), adjusting for age, sex and variables that have been shown to be potential confounders with air pollution and BMI, such as household education (Inoue et al., 2023), race, ethnicity (Geldsetzer et al., 2024), puberty (Song et al., 2023), and urbanicity (Sajjad Abdollahpour et al., 2024). To account also for co-exposure, models for PM2.5 additionally included NO2 and O3 estimates as covariates; models for NO2 included PM2.5 and O3; models for O3 included PM2.5 and NO2; and models for Oxwt included PM2.5. Site and participant identifiers were included as crossed random effects in the model.

#### 2.6.2 Air pollution, brain subcortical microstructure, and weight gain

Following this analysis, only pollutants showing associations with BMI were evaluated to examine whether air pollution moderated the relationship between brain microstructure and BMI. LME models included BMI as the dependent variable and RNI and RND (across 16 subcortical ROIs, separate models) as independent variables while controlling for age, sex and variables that have been shown to be potential confounders with brain structure, BMI and/or air pollution, including household education, a marker of socioeconomic status (Inoue et al., 2023), urbanicity (Polemiti et al., 2024; Sajjad Abdollahpour et al., 2024), race, ethnicity (Geldsetzer et al., 2024), puberty (Holm et al., 2023), and in-scanner head motion measured as the mean framewise displacement during the DTI acquisition. Consistent with the previous statistical analyses, co-exposure effects were also accounted for by including air pollutants as covariates in LMEs models.

All continuous variables (e.g., ROIs, air pollutants, puberty, and head motion) were mean-centered. In contrast, categorical variables (e.g., education, sex, race, ethnicity, and urbanicity) were effects coded. Models also included a two-way interaction between ROI and each air pollutant and the three-way interaction between ROI, air pollutants and age (as a proxy of time), which were considered of interest in the present study. Furthermore, all LMEs models included the MRI scanner and participant identifier (ID) as crossed random effects.

The main analysis included both males and females to evaluate the moderating effects of air pollution on the relationship between subcortical microstructure and weight gain. However, given findings showing that air pollution (Cotter et al., 2024; Peterson et al., 2022) has differential effects on brain structure in males and females, we also conducted sex-stratified LME models. Using the Benjamini-Hochberg method (Benjamini & Hochberg, 1995), the *p*-values of statistical models were adjusted separately for RNI (16 regions) and RND (16 regions). Standardized beta coefficients (β) and confidence intervals (CI) are also reported. The model formula for air pollutants that previously showed an association with BMI across sexes is detailed below:

### PM2.5

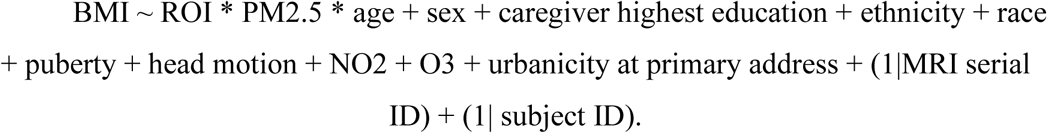

**Note:** Sex-stratified models were the same but without the sex covariate.

## 3. RESULTS

### 3.1 Demographics

Demographic data of baseline and follow-up visits are summarized in **Table 1**. After applying all exclusion criteria, the baseline sample comprised 4,802 participants, with a mean age of 9.93 ± 0.62 years. Among them, there were 2,397 males and 2,405 females. The follow-up sample included 2,439 individuals, with a mean age of 11.91 ± 0.64 years, consisting of 1,246 males and 1,193 females.

**Table 1.** Sample characteristics at baseline and year 2 visits.

|  | **Baseline**<br>**N = 4,802** | **Year 2**<br>**N = 2,439** |
| --- | --- | --- |
| **Age, years (mean, SD)** | 9.93 (0.62) | 11.91 (0.64) |
| **Puberty, (mean, SD)** | 1.93 (0.82) | 2.67 (0.99) |
| **BMI, (mean, SD)** | 18.73 (3.75) | 20.44 (4.36) |
| **Pollutant levels (mean, SD)** |  |  |
| *PM2.5 $\mu\text{g}/\text{m}^3$* | 7.59 (1.56) | 7.55 (1.53) |
| *NO2, ppb* | 18.71 (5.75) | 18.71 (5.8) |
| *O3, ppb* | 41.74 (4.47) | 41.56 (4.43) |
| *Oxwt, ppb* | 33.9 (3.49) | 33.79 (3.57) |
| **Sex, n (%)** |  |  |
| *Female* | 2405 (50.1%) | 1193 (48.9%) |
| *Male* | 2397 (49.9%) | 1246 (51.1%) |
| **Weight status, n (%)** |  |  |
| *Healthy weight* | 3335 (69.5%) | 1663 (68.2%) |
| *Overweight* | 756 (15.7%) | 395 (16.2%) |
| *Obese* | 711 (14.8%) | 381 (15.6%) |
| **Ethnicity, n (%)** |  |  |
| *Latinx* | 1014 (21.1%) | 499 (20.5%) |
| *Non-Latinx* | 3788 (78.9%) | 1940 (79.5%) |
| **Race** |  |  |
| *Asian* | 115 (2.4%) | 52 (2.1%) |
| *Black* | 617 (12.8%) | 250 (10.3%) |
| *Mixed* | 547 (11.4%) | 249 (10.2%) |
| *NHPI/AINA* | 24 (0.5%) | 8 (0.3%) |
| *Other* | 230 (4.8%) | 109 (4.5%) |
| *White* | 3269 (68.1%) | 1771 (72.6%) |
| **Highest household education, n (%)** |  |  |
| *< HS Diploma* | 194 (4%) | 85 (3.5%) |
| *HS Diploma/GED* | 348 (7.2%) | 150 (6.2%) |
| *Some College* | 1097 (22.8%) | 542 (22.2%) |
| *Bachelor's degree* | 1274 (26.5%) | 688 (28.2%) |
| *Post Graduate Degree* | 1889 (39.3%) | 974 (39.9%) |
| **Urbanicity** |  |  |
| *Rural* | 393 (8.2%) | 199 (8.2%) |
| *Urban clusters* | 160 (3.3%) | 90 (3.7%) |
| *Urbanized* | 4249 (88.5%) | 2150 (88.2%) |
| **Percentage of time at primary address, mean (SD)** | 97.01 (10.98) | 96.87 (11.11) |
| **Number of years living at primary address, mean (SD)** | 5.66 (3.77) | 5.75 (3.75) |

**Note:** BMI = Body Mass Index. NHPI / AINA = American Indian, Alaska Native/Native Hawaiian, Pacific Islander; HS = High school; GED = Generalized Education Degree. Race and ethnicity were self-reported by the caregiver and collapsed into six and two categories, respectively. PM2.5 was measured in micrograms per cubic meter ( $\mu\text{g}/\text{m}^3$ ), while NO2 and O3 were measured in parts per billion (ppb), all at a 1 km<sup>2</sup> resolution. Urbanicity of participant's addresses was assessed with a categorical variable indicating if participants' addresses were in census tracts considered to be urban (50,000 or more people), urban clusters (at least 2,500 and less than 50,000 people), or rural (less than 2,500 people per tract, and/or not included in urban areas or clusters). These categories are
established according to the Census Tract Urban Classification and based on the 2010 census estimates.

**Table 2.** Main effects and interactions of linear mixed effect models evaluating the relationship between each air pollutant and BMI across sexes and stratified by sex.

|  | Pollutant | Pollutant |  | Age |  | Pollutant X Age |  |
| --- | --- | --- | --- | --- | --- | --- | --- |
| | | $\beta$ (95%CI) | p-value | $\beta$ (95%CI) | p-value | $\beta$ (95%CI) | p-value |
| Across sexes | PM2.5 | 0.145 (-0.002, 0.291) | 0.056 | 0.848 (0.791, 0.894) | **<0.001** | 0.053 (0.013, 0.093) | **0.010** |
|  | NO2 | 0.088 (-0.065, 0.241) | 0.260 | 0.841 (0.789, 0.893) | **<0.001** | 0.014 (-0.025, 0.054) | 0.484 |
|  | O3 | 0.022 (-0.101, 0.145) | 0.727 | 0.840 (0.788, 0.892) | **<0.001** | -0.018 (-0.059, 0.022) | 0.374 |
|  | Oxwt | 0.057 (-0.072, 0.185) | 0.389 | 0.841 (0.789, 0.893) | **<0.001** | -0.007 (-0.047, 0.033) | 0.735 |
| Males | PM2.5 | 0.200 (0.003, 0.039) | **0.048** | 0.873 (0.810, 0.936) | **< 0.001** | 0.060 (0.004, 0.116) | **0.033** |
|  | NO2 | 0.053 (-0.153, 0.258) | 0.616 | 0.868 (0.805, 0.931) | **<0.001** | -0.019 (-0.072, 0.034) | 0.478 |
|  | O3 | 0.027 (-0.142, 0.196) | 0.754 | 0.866 (0.803, 0.929) | **<0.001** | -0.046 (-0.101, 0.009) | 0.101 |
|  | Oxwt | 0.048 (-0.129, 0.225) | 0.597 | 0.866 (0.803, 0.929) | **<0.001** | -0.049 (-0.104, 0.005) | 0.077 |
| Females | PM2.5 | 0.117 (-0.050, 0.284) | 0.176 | 0.765 (0.675, 0.855) | **<0.001** | 0.052 (-0.008, 0.111) | 0.088 |
|  | NO2 | -0.001 (-0.175, 0.173) | 0.995 | 0.771 (0.681, 0.861) | **<0.001** | 0.054 (-0.005, 0.114) | 0.075 |
|  | O3 | 0.077 (-0.078, 0.232) | 0.331 | 0.769 (0.679, 0.859) | **<0.001** | 0.013 (-0.046, 0.072) | 0.671 |
|  | Oxwt | 0.071 (-0.092, 0.233) | 0.396 | 0.770 (0.680, 0.860) | **<0.001** | 0.039 (-0.019, 0.098) | 0.187 |

**Note:** Estimated regression coefficients ( $\beta$ ), 95% confidence intervals (CI) and p-values from the linear mixed-effects models evaluating the relationship between each air pollutant and Body Mass Index. Models were controlled for age, sex, education, race, ethnicity, puberty, co-exposure to other air pollutants and urbanicity. Random intercepts were modelled from site and subject ID. Sex-stratified models were the same but without the sex covariate.

Most participants identified as white (68.1%), 21.1% identified as Latinx, 39.3% had at least one caregiver with a postgraduate degree and the majority lived in urbanized areas with 50,000 or more people (88.5%). Regarding BMI, 69.5% of participants had a healthy weight at baseline, while 15.7% had overweight and 14.8% had obesity. Two years later, these proportions remained similar (68.2% healthy weight, 16.2% overweight and 15.6% obesity). The mean concentration of each air pollutant was as follows: PM2.5 had a mean of 7.59± 1.56 µg/m³; NO2 had a mean of 18.71 ± 5.75 ppb; O3 had a mean of 41.74 ± 4.47 ppb, and Oxwt had a mean of 33.9 ± 3.49.

### 3.2 Associations between air pollutants and BMI

Across both sexes, LMEs models revealed a significant interaction between PM2.5 and time, where higher exposure to PM2.5 was associated with an increase in BMI over time (β = 0.053, CI95% = [0.013, 0.093], *p*-value = 0.010). There were no associations between any other pollutant (i.e., NO2, O3, and Oxwt) and BMI (see Table 2, and Figure 1).

**Figure 1.**
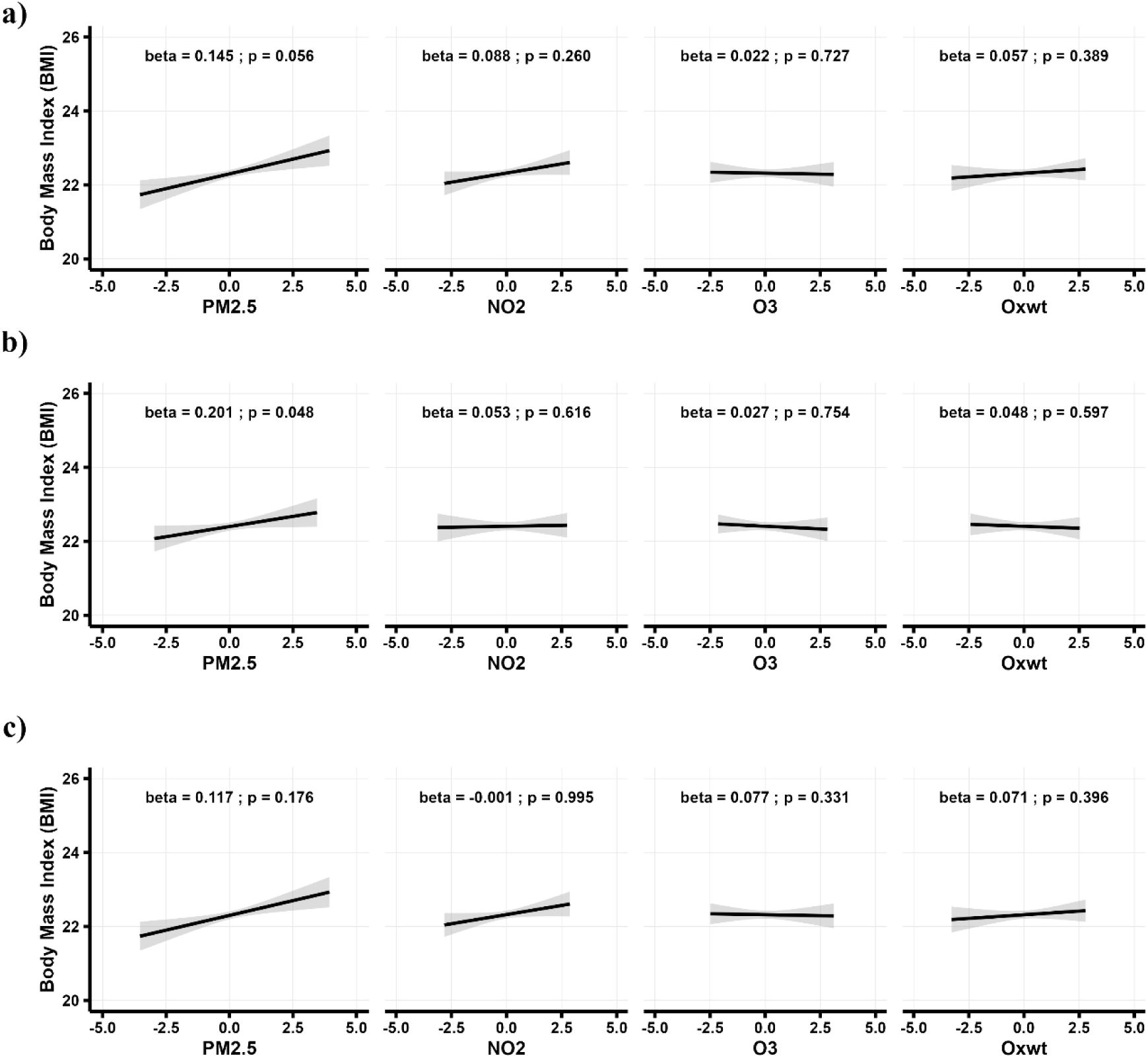
Relationships between each air pollutant and Body Mass Index (BMI), across sexes (a), in males (b) and in females (c). Linear mixed-effects models were adjusted for age, sex, household education, ethnicity, race, puberty, in-scanner head motion, co-exposure to other air pollutants, and urbanicity at primary address. Random intercepts were modelled from MRI scanner device and subject ID.

**Table 2.**
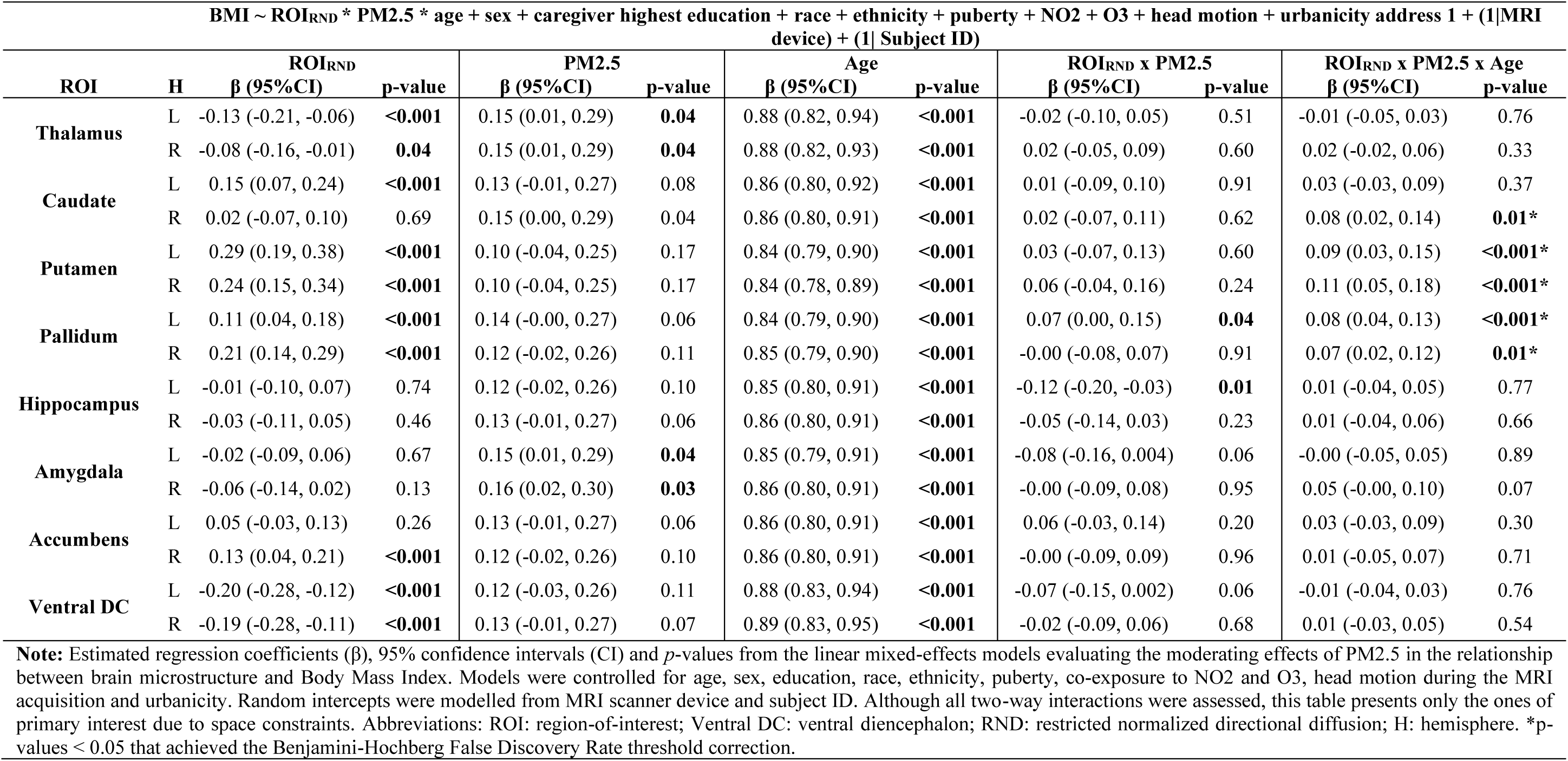
Relationship between PM2.5, subcortical restricted intracellular normalized directional diffusion (RND), and body mass index (BMI) across both sex over time.

| BMI ~ ROI<sub>RND</sub> * PM2.5 * age + sex + caregiver highest education + race + ethnicity + puberty + NO2 + O3 + head motion + urbanicity address 1 + (1\|MRI device) + (1\| Subject ID) |  |  |  |  |  |  |  |  |  |  |  |
| --- | --- | --- | --- | --- | --- | --- | --- | --- | --- | --- | --- |
| ROI | H | ROI<sub>RND</sub> |  | PM2.5 |  | Age |  | ROI<sub>RND</sub> x PM2.5 |  | ROI<sub>RND</sub> x PM2.5 x Age |  |
|  |  | β (95%CI) | p-value | β (95%CI) | p-value | β (95%CI) | p-value | β (95%CI) | p-value | β (95%CI) | p-value |
| Thalamus | L | -0.13 (-0.21, -0.06) | <0.001 | 0.15 (0.01, 0.29) | 0.04 | 0.88 (0.82, 0.94) | <0.001 | -0.02 (-0.10, 0.05) | 0.51 | -0.01 (-0.05, 0.03) | 0.76 |
|  | R | -0.08 (-0.16, -0.01) | 0.04 | 0.15 (0.01, 0.29) | 0.04 | 0.88 (0.82, 0.93) | <0.001 | 0.02 (-0.05, 0.09) | 0.60 | 0.02 (-0.02, 0.06) | 0.33 |
| Caudate | L | 0.15 (0.07, 0.24) | <0.001 | 0.13 (-0.01, 0.27) | 0.08 | 0.86 (0.80, 0.92) | <0.001 | 0.01 (-0.09, 0.10) | 0.91 | 0.03 (-0.03, 0.09) | 0.37 |
|  | R | 0.02 (-0.07, 0.10) | 0.69 | 0.15 (0.00, 0.29) | 0.04 | 0.86 (0.80, 0.91) | <0.001 | 0.02 (-0.07, 0.11) | 0.62 | 0.08 (0.02, 0.14) | 0.01* |
| Putamen | L | 0.29 (0.19, 0.38) | <0.001 | 0.10 (-0.04, 0.25) | 0.17 | 0.84 (0.79, 0.90) | <0.001 | 0.03 (-0.07, 0.13) | 0.60 | 0.09 (0.03, 0.15) | <0.001* |
|  | R | 0.24 (0.15, 0.34) | <0.001 | 0.10 (-0.04, 0.25) | 0.17 | 0.84 (0.78, 0.89) | <0.001 | 0.06 (-0.04, 0.16) | 0.24 | 0.11 (0.05, 0.18) | <0.001* |
| Pallidum | L | 0.11 (0.04, 0.18) | <0.001 | 0.14 (-0.00, 0.27) | 0.06 | 0.84 (0.79, 0.90) | <0.001 | 0.07 (0.00, 0.15) | 0.04 | 0.08 (0.04, 0.13) | <0.001* |
|  | R | 0.21 (0.14, 0.29) | <0.001 | 0.12 (-0.02, 0.26) | 0.11 | 0.85 (0.79, 0.90) | <0.001 | -0.00 (-0.08, 0.07) | 0.91 | 0.07 (0.02, 0.12) | 0.01* |
| Hippocampus | L | -0.01 (-0.10, 0.07) | 0.74 | 0.12 (-0.02, 0.26) | 0.10 | 0.85 (0.80, 0.91) | <0.001 | -0.12 (-0.20, -0.03) | 0.01 | 0.01 (-0.04, 0.05) | 0.77 |
|  | R | -0.03 (-0.11, 0.05) | 0.46 | 0.13 (-0.01, 0.27) | 0.06 | 0.86 (0.80, 0.91) | <0.001 | -0.05 (-0.14, 0.03) | 0.23 | 0.01 (-0.04, 0.06) | 0.66 |
| Amygdala | L | -0.02 (-0.09, 0.06) | 0.67 | 0.15 (0.01, 0.29) | 0.04 | 0.85 (0.79, 0.91) | <0.001 | -0.08 (-0.16, 0.004) | 0.06 | -0.00 (-0.05, 0.05) | 0.89 |
|  | R | -0.06 (-0.14, 0.02) | 0.13 | 0.16 (0.02, 0.30) | 0.03 | 0.86 (0.80, 0.91) | <0.001 | -0.00 (-0.09, 0.08) | 0.95 | 0.05 (-0.00, 0.10) | 0.07 |
| Accumbens | L | 0.05 (-0.03, 0.13) | 0.26 | 0.13 (-0.01, 0.27) | 0.06 | 0.86 (0.80, 0.91) | <0.001 | 0.06 (-0.03, 0.14) | 0.20 | 0.03 (-0.03, 0.09) | 0.30 |
|  | R | 0.13 (0.04, 0.21) | <0.001 | 0.12 (-0.02, 0.26) | 0.10 | 0.86 (0.80, 0.91) | <0.001 | -0.00 (-0.09, 0.09) | 0.96 | 0.01 (-0.05, 0.07) | 0.71 |
| Ventral DC | L | -0.20 (-0.28, -0.12) | <0.001 | 0.12 (-0.03, 0.26) | 0.11 | 0.88 (0.83, 0.94) | <0.001 | -0.07 (-0.15, 0.002) | 0.06 | -0.01 (-0.04, 0.03) | 0.76 |
|  | R | -0.19 (-0.28, -0.11) | <0.001 | 0.13 (-0.01, 0.27) | 0.07 | 0.89 (0.83, 0.95) | <0.001 | -0.02 (-0.09, 0.06) | 0.68 | 0.01 (-0.03, 0.05) | 0.54 |

**Note:** Estimated regression coefficients (β), 95% confidence intervals (CI) and *p*-values from the linear mixed-effects models evaluating the moderating effects of PM2.5 in the relationship between brain microstructure and Body Mass Index. Models were controlled for age, sex, education, race, ethnicity, puberty, co-exposure to NO2 and O3, head motion during the MRI acquisition and urbanicity. Random intercepts were modelled from MRI scanner device and subject ID. Although all two-way interactions were assessed, this table presents only the ones of primary interest due to space constraints. Abbreviations: ROI: region-of-interest; Ventral DC: ventral diencephalon; RND: restricted normalized directional diffusion; H: hemisphere. \**p*-values < 0.05 that achieved the Benjamini-Hochberg False Discovery Rate threshold correction.

Results of sex-stratified LMEs models assessing the relationship between each air pollutant and BMI revealed similar findings in males. LMEs revealed a significant interaction between PM2.5 and age, where higher exposure to PM2.5 was associated with an increase in BMI over time (β = 0.060, CI95% = [0.004, 0.115], *p*-value = 0.033). There were no significant associations between any other pollutant and BMI in males and females (see Table 2, and Figure 1).

### 3.3 Air pollution, brain subcortical microstructure, and weight gain

#### RNI

Across sexes, although RNI was associated with BMI, there were no interactions between RNI and air pollution, nor three-way interactions between RNI, air pollution and time. Similarly, this did not differ by sex. Further details including main effects and interactions of RNI and air pollutants on BMI can be found in the **Supplemental Materials** (Tables across sexes S2-S5; males: Tables S6–S9; females: Tables S10–S13).

#### RND across sexes

LMEs for NO2, O3, and Oxwt revealed that, regardless of time and air pollution exposure, high RND was significantly associated with higher BMI in some subcortical regions (e.g., bilateral putamen and pallidum). However, in other regions such as the bilateral hippocampus and amygdala, we observed the opposite relationship (i.e., lower RND was associated with higher BMI). Further details can be found in the **Supplemental Materials Tables S14–S17**. However, in the case of PM2.5, we found a significant three way interaction (RND x PM2.5 x Age) revealing that exposure to this air pollutant significantly moderated the positive association between RND and BMI over time in the right caudate nucleus (β_right_=0.077, CI95% = [0.018, 0.137], *p*=0.011, see **Table 3** and **Figure 2**) as well as the bilateral putamen (β_left_=0.094, CI95% = 0.033, 0.155, *p*=0.002; β_right_=0.114, CI95% = 0.052, 0.176, *p*<0.001, see **Table 3** and **Figure 3**) and pallidum (β_left_=0.083, CI95% = 0.038, 0.128, *p*<0.001; β_right_=0.067, CI95% = 0.016, 0.117, *p*=0.010, see **Table 3** and **Figure 4**).

**Figure 2.**
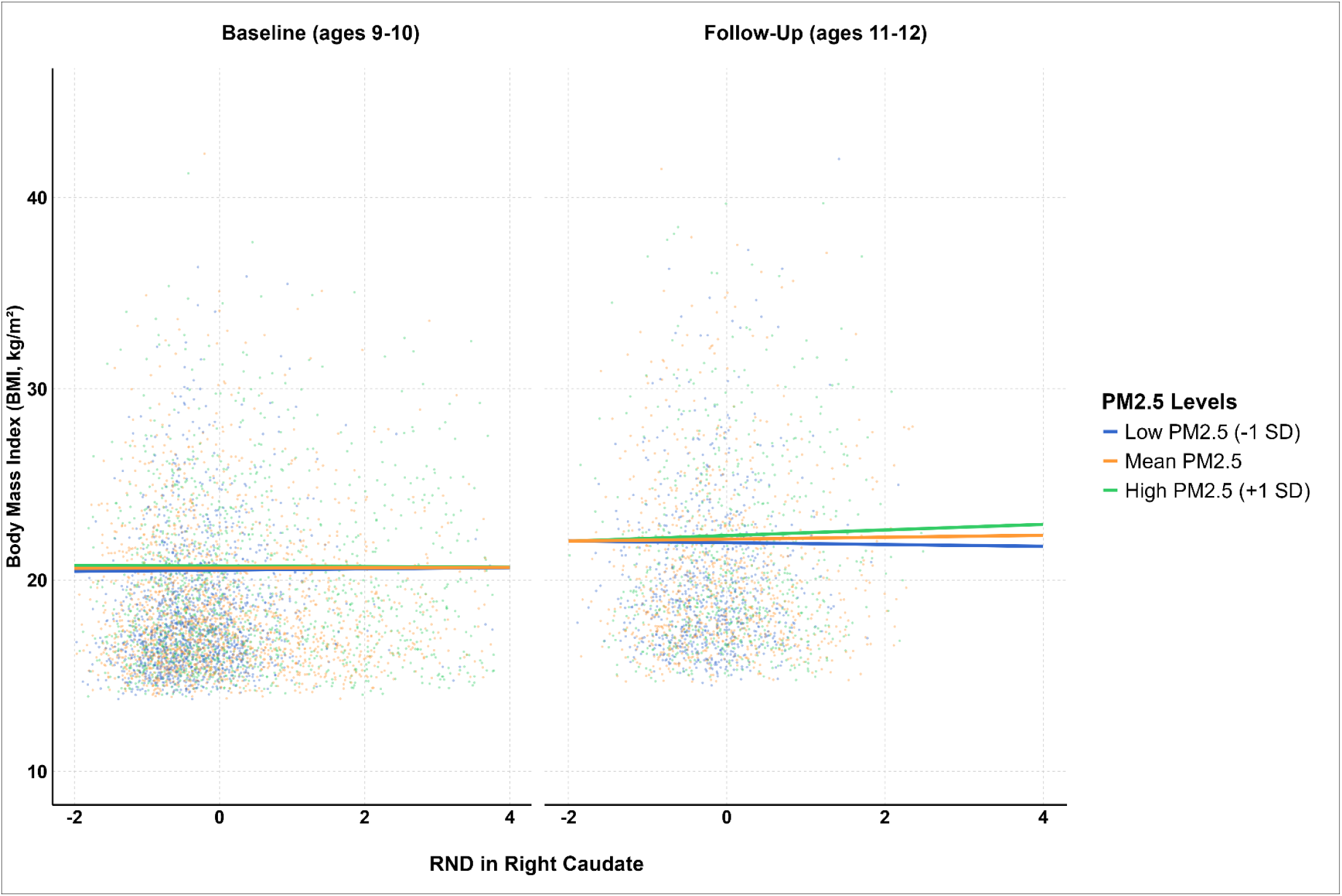
Moderating effects across both sexes at different levels of PM2.5 (at the mean and ±1SD) on the relationship between restricted normalized directional (RND) diffusion in the right caudate nucleus and Body Mass Index (BMI). Linear mixed-effects models were adjusted for age, sex, household education, ethnicity, race, puberty, in-scanner head motion, co-exposure to NO2 and O3, urbanicity at primary address. Random intercepts were modelled from MRI scanner device and subject ID.

**Figure 3.**
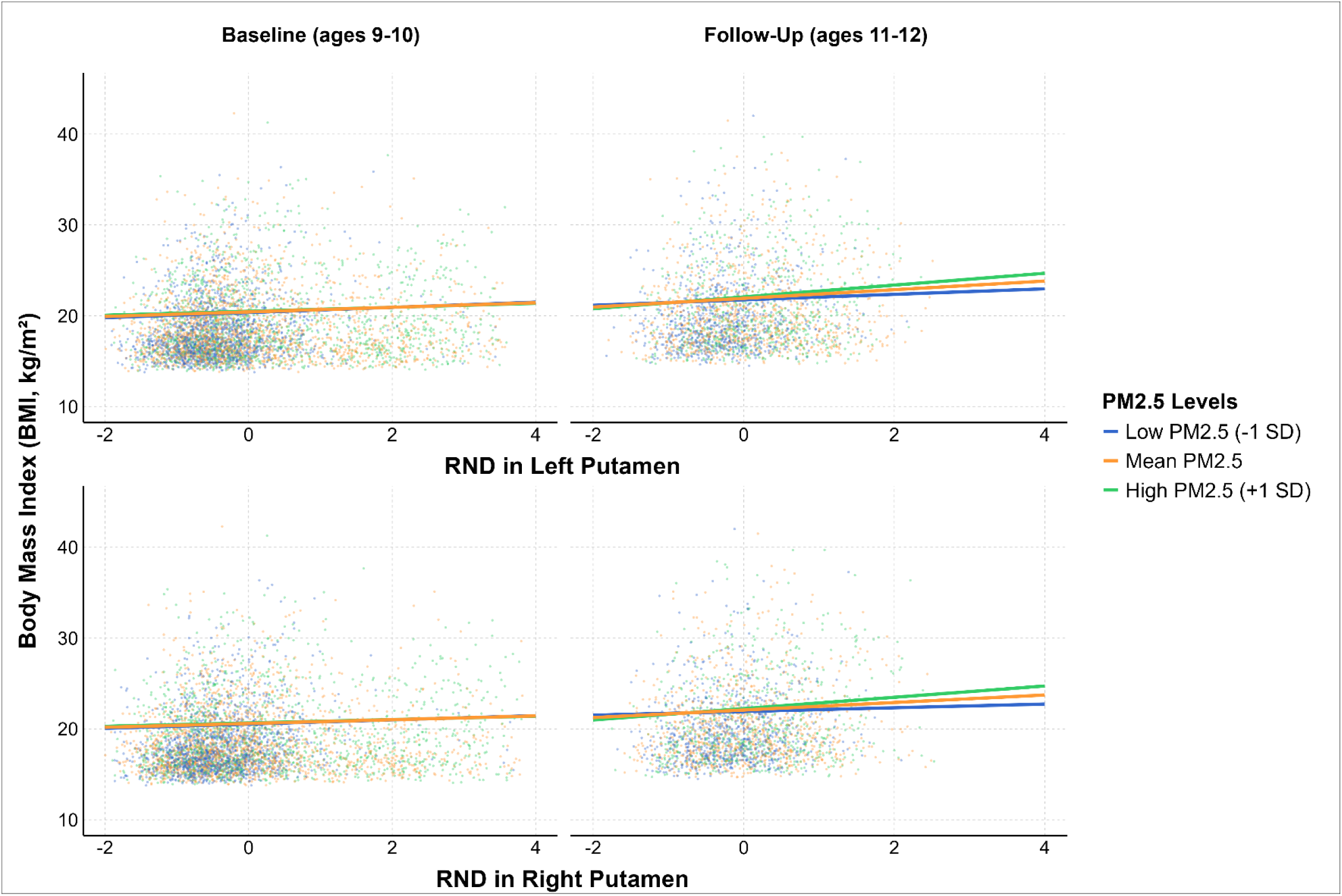
Moderating effects across both sexes at different levels of PM2.5 (at the mean and ±1SD) on the relationship between restricted normalized directional (RND) diffusion in the bilateral putamen and Body Mass Index (BMI). Linear mixed-effects models were adjusted for age, sex, household education, ethnicity, race, puberty, in-scanner head motion, co-exposure to NO2 and O3, urbanicity at primary address. Random intercepts were modelled from MRI scanner device and subject ID.

**Figure 4.**
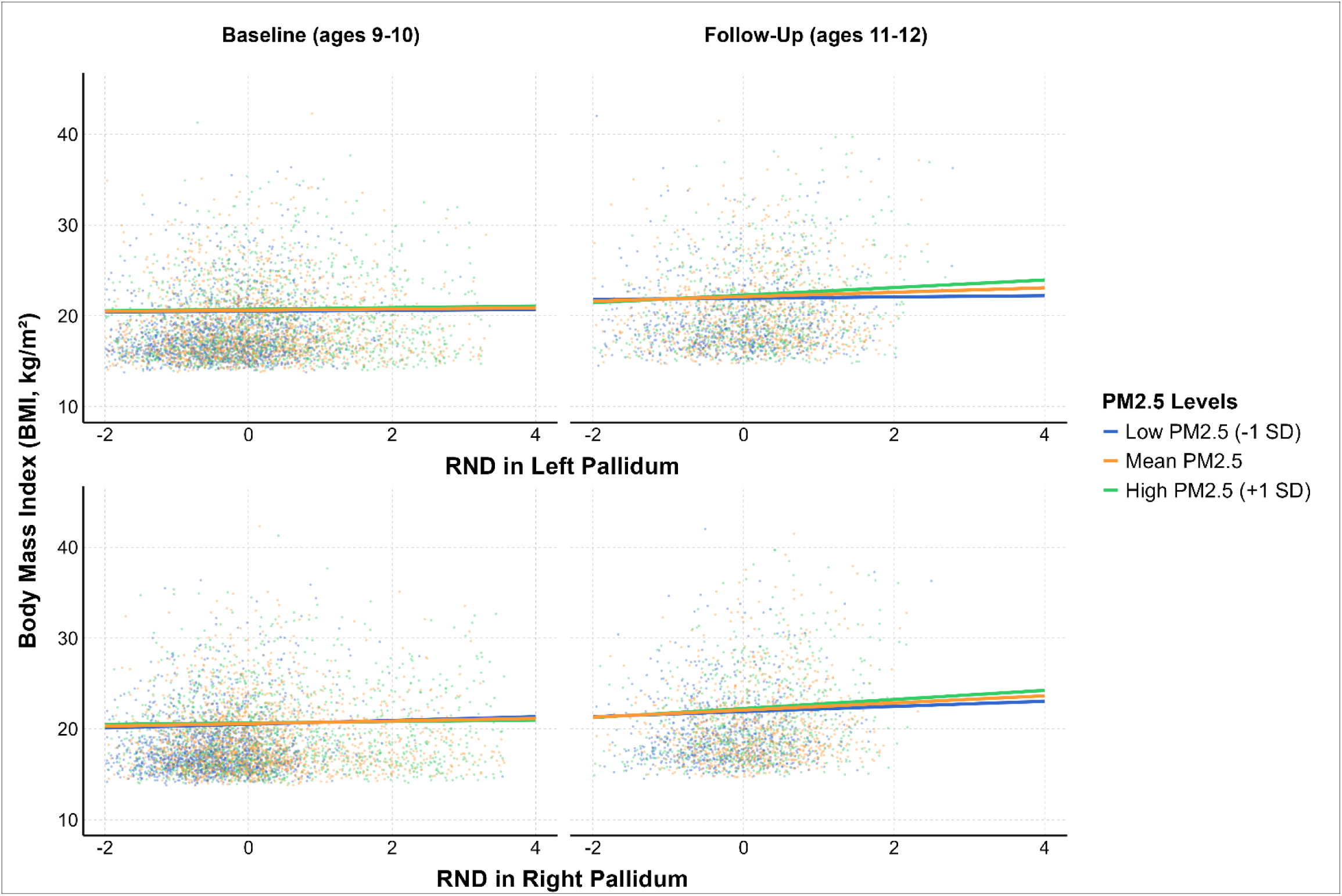
Moderating effects across both sexes at different levels of PM2.5 (at the mean and ±1SD) on the relationship between restricted normalized directional (RND) diffusion in the bilateral pallidum and Body Mass Index (BMI). Linear mixed-effects models were adjusted for age, sex, household education, ethnicity, race, puberty, in-scanner head motion, co-exposure to NO2 and O3, urbanicity at primary address. Random intercepts were modelled from MRI scanner device and subject ID.

**Table 3.** Relationship between PM2.5, subcortical restricted normalized directional diffusion (RND), and body mass index (BMI) in males over time.

| BMI ~ ROI<sub>RND</sub> * PM2.5 * age + caregiver highest education + race + ethnicity + puberty + NO2 + O3 + head motion + urbanicity address 1 + (1\|MRI device) + (1\| Subject ID) |  |  |  |  |  |  |  |  |  |  |  |
| --- | --- | --- | --- | --- | --- | --- | --- | --- | --- | --- | --- |
| ROI | H | ROI<sub>RND</sub><br>β (95%CI) | p-value | PM2.5<br>β (95%CI) | p-value | Age<br>β (95%CI) | p-value | ROI<sub>RND</sub> x PM2.5<br>β (95%CI) | p-value | ROI<sub>RND</sub> x PM2.5 x Age<br>β (95%CI) | p-value |
| Thalamus | L | -0.12 (-0.22, -0.02) | **0.02** | 0.22 (0.04, 0.40) | **0.02** | 0.92 (0.85, 0.99) | **<0.001** | 0.06 (-0.04, 0.16) | 0.24 | -0.01 (-0.06, 0.05) | 0.80 |
|  | R | -0.14 (-0.24, -0.04) | **0.01** | 0.21 (0.04, 0.39) | **0.02** | 0.93 (0.86, 1.00) | **<0.001** | 0.12 (0.02, 0.22) | **0.02** | -0.00 (-0.06, 0.05) | 0.88 |
| Caudate | L | 0.10 (-0.02, 0.22) | 0.10 | 0.19 (0.00, 0.38) | **0.05** | 0.89 (0.82, 0.96) | **<0.001** | 0.09 (-0.04, 0.22) | 0.18 | 0.07 (-0.02, 0.15) | 0.13 |
|  | R | -0.03 (-0.15, 0.08) | 0.55 | 0.21 (0.02, 0.40) | **0.03** | 0.90 (0.83, 0.96) | **<0.001** | 0.10 (-0.02, 0.22) | 0.10 | 0.10 (0.02, 0.18) | **0.02** |
| Putamen | L | 0.31 (0.18, 0.44) | **<0.001** | 0.14 (-0.05, 0.33) | 0.16 | 0.88 (0.81, 0.94) | **<0.001** | 0.08 (-0.06, 0.22) | 0.27 | 0.13 (0.04, 0.21) | **0.002\*** |
|  | R | 0.28 (0.16, 0.41) | **<0.001** | 0.15 (-0.04, 0.34) | 0.13 | 0.87 (0.80, 0.94) | **<0.001** | 0.16 (0.03, 0.29) | **0.02** | 0.18 (0.10, 0.26) | **<0.001\*** |
| Pallidum | L | 0.09 (-0.00, 0.19) | **0.05** | 0.18 (-0.00, 0.37) | 0.06 | 0.88 (0.81, 0.95) | **<0.001** | 0.05 (-0.05, 0.15) | 0.35 | 0.10 (0.04, 0.17) | **0.002\*** |
|  | R | 0.24 (0.14, 0.34) | **<0.001** | 0.19 (0.00, 0.38) | **0.05** | 0.89 (0.82, 0.96) | **<0.001** | 0.10 (-0.01, 0.21) | 0.07 | 0.15 (0.08, 0.22) | **<0.001\*** |
| Hippocampus | L | -0.06 (-0.17, 0.05) | 0.29 | 0.18 (-0.01, 0.37) | 0.07 | 0.89 (0.82, 0.96) | **<0.001** | -0.10 (-0.22, 0.02) | 0.09 | -0.02 (-0.08, 0.05) | 0.63 |
|  | R | -0.03 (-0.14, 0.08) | 0.57 | 0.20 (0.01, 0.38) | **0.04** | 0.90 (0.83, 0.96) | **<0.001** | 0.01 (-0.10, 0.13) | 0.81 | 0.04 (-0.03, 0.10) | 0.28 |
| Amygdala | L | 0.02 (-0.09, 0.13) | 0.69 | 0.21 (0.03, 0.39) | **0.03** | 0.89 (0.82, 0.96) | **<0.001** | -0.00 (-0.12, 0.12) | 0.99 | 0.03 (-0.04, 0.10) | 0.43 |
|  | R | -0.02 (-0.13, 0.08) | 0.66 | 0.20 (0.02, 0.39) | **0.03** | 0.89 (0.83, 0.96) | **<0.001** | 0.03 (-0.08, 0.15) | 0.58 | 0.05 (-0.02, 0.13) | 0.13 |
| Accumbens | L | 0.03 (-0.08, 0.14) | 0.60 | 0.17 (-0.01, 0.36) | 0.07 | 0.89 (0.82, 0.96) | **<0.001** | 0.07 (-0.05, 0.19) | 0.23 | 0.05 (-0.02, 0.13) | 0.16 |
|  | R | 0.08 (-0.04, 0.20) | 0.17 | 0.18 (-0.00, 0.37) | 0.06 | 0.89 (0.82, 0.96) | **<0.001** | 0.08 (-0.05, 0.20) | 0.24 | 0.07 (-0.01, 0.15) | 0.11 |
| Ventral DC | L | -0.16 (-0.27, -0.05) | **0.004** | 0.19 (0.00, 0.38) | **0.05** | 0.92 (0.85, 0.99) | **<0.001** | 0.00 (-0.10, 0.11) | 0.96 | 0.01 (-0.04, 0.05) | 0.79 |
|  | R | -0.15 (-0.26, -0.04) | **0.01** | 0.20 (0.02, 0.38) | **0.04** | 0.92 (0.86, 0.99) | **<0.001** | 0.06 (-0.04, 0.17) | 0.24 | 0.01 (-0.04, 0.06) | 0.60 |

**Note:** Estimated regression coefficients (β), 95% confidence intervals (CI) and *p*-values from the linear mixed-effects models evaluating the moderating effects of PM2.5 in the relationship between brain microstructure and Body Mass Index. Models were controlled for age, education, race, ethnicity, puberty, co-exposure to NO2 and O3, head motion during the MRI acquisition and urbanicity. Random intercepts were modelled from MRI scanner device and subject ID. Although all two-way interactions were assessed, this table presents only the ones of primary interest due to space constraints. Abbreviations: ROI: region-of-interest; Ventral DC: ventral diencephalon; RND: restricted normalized directional diffusion; H: hemisphere. \**p*-values < 0.05 that achieved the Benjamini-Hochberg False Discovery Rate threshold correction.

#### RND in Males

We found the same pattern of results in males. LMEs models for NO2, O3, and Oxwt, showed that, regardless of time and air pollution exposure, high RND was significantly associated with higher BMI in some subcortical regions, although other regions showed the opposite relationship (i.e., lower RND was associated with higher BMI). Further details can be found in the **Supplemental Materials Tables S18-21**. For the PM2.5 pollutant, there were significant 3-way interactions (RND x PM2.5 X Age) on BMI in the bilateral putamen (β_left_=0.126, CI95% = [0.044, 0.207], *p*=0.002; β_right_=0.177, CI95% = [0.095, 0.258], *p*< 0.001, see **Table 4** and **Figure 5**) and pallidum (β_left_=0.103, CI95% = [0.037, 0.169], *p*= 0.002; β_right_=0.148, CI95% = [0.079, 0.218], *p*<0.001, see **Table 4** and **Figure 6**). An interaction effect was also found in the right caudate; however, this result did not survive correction for multiple comparisons. Across these regions, increased exposure to PM2.5 coupled with increased RND was significantly associated with BMI increases over time.

**Figure 5.**
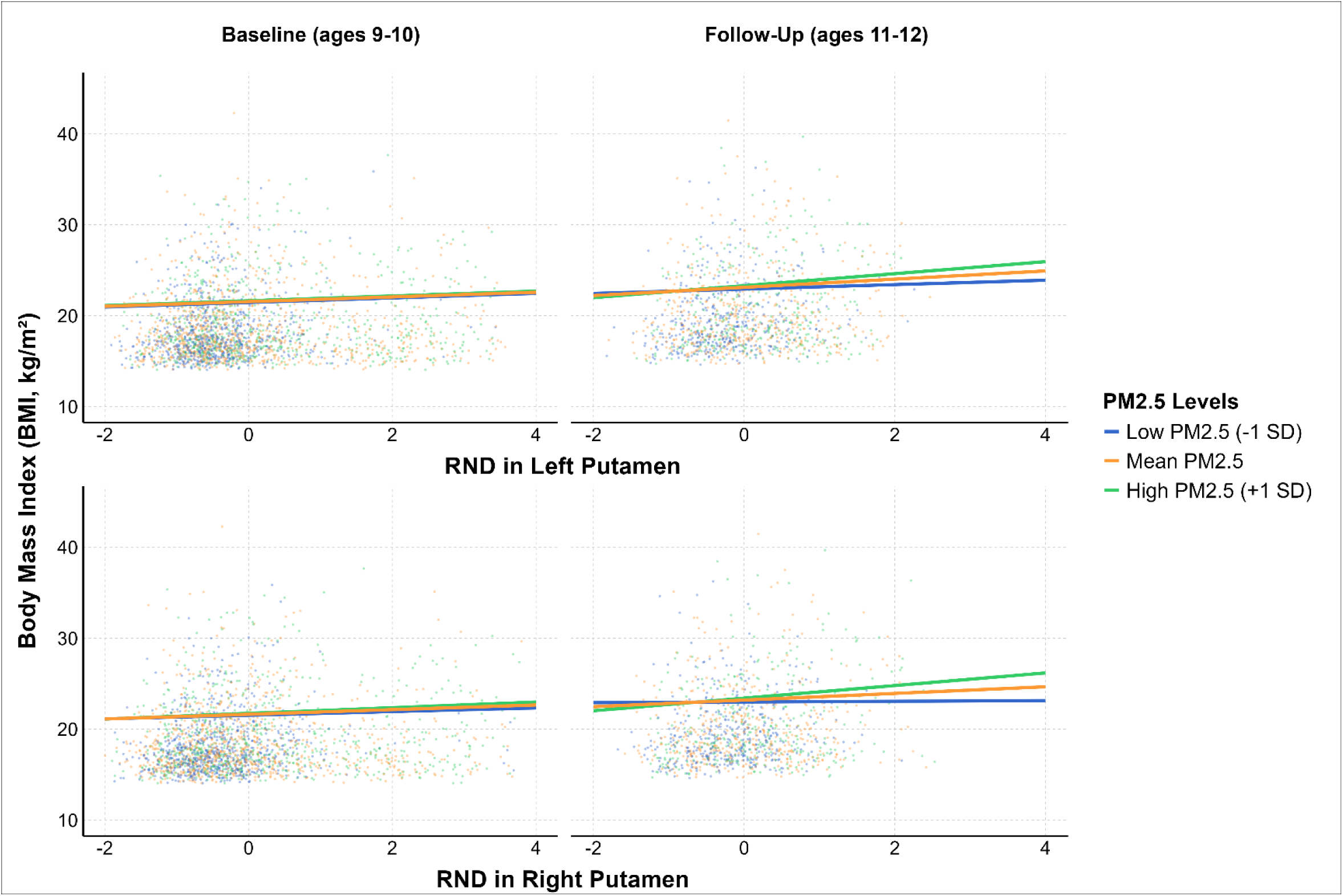
Moderating effects in males at different levels of PM2.5 (at the mean and ±1SD) on the relationship between restricted normalized directional (RND) diffusion in the bilateral putamen and Body Mass Index (BMI). Linear mixed-effects models were adjusted for age, sex, household education, ethnicity, race, puberty, in-scanner head motion, co-exposure to NO2 and O3, urbanicity at primary address. Random intercepts were modelled from MRI scanner device and subject ID.

**Figure 6.**
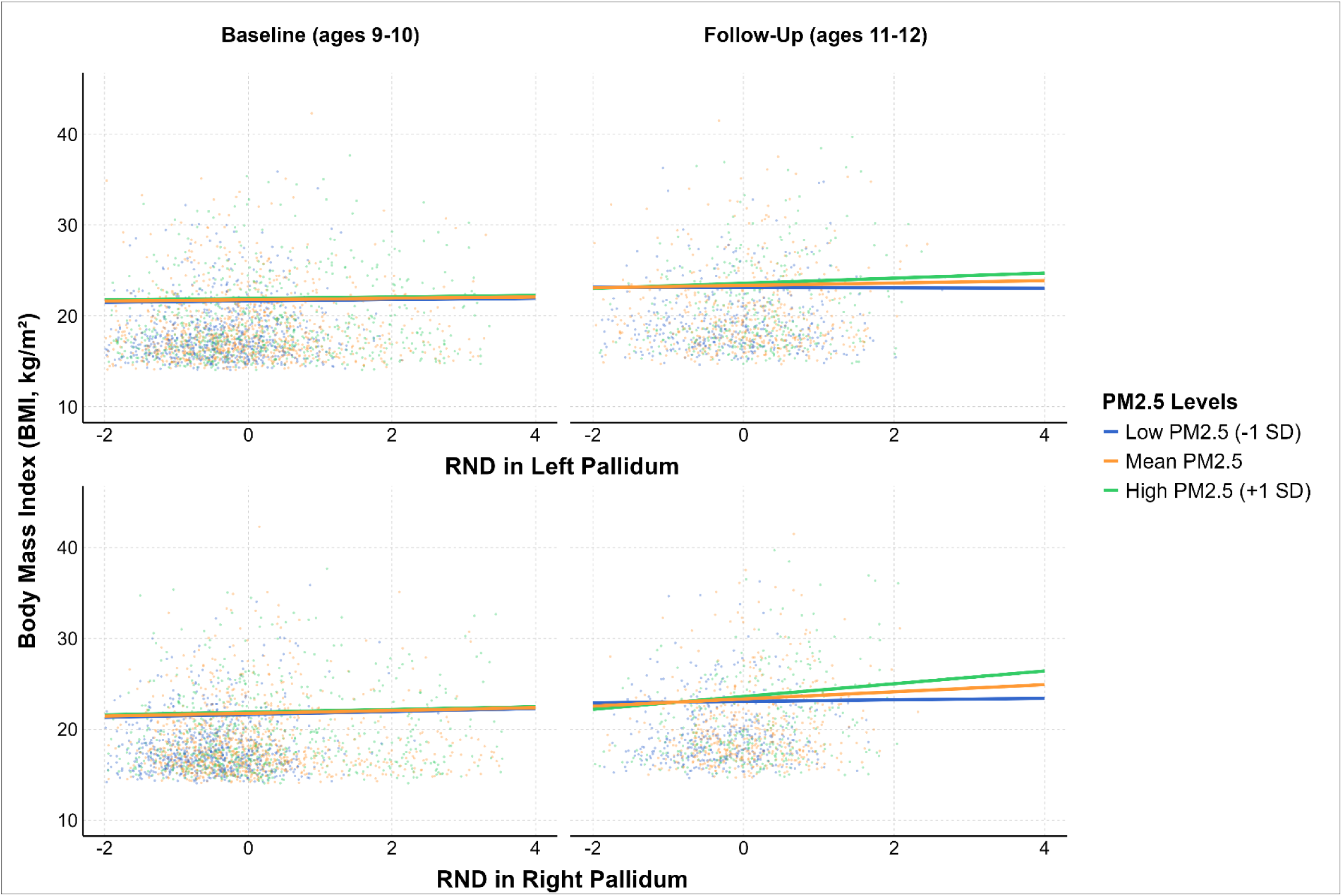
Moderating effects in males at different levels of PM2.5 (at the mean and ±1SD) on the relationship between restricted normalized directional (RND) diffusion in the bilateral pallidum and Body Mass Index (BMI). Linear mixed-effects models were adjusted for age, sex, household education, ethnicity, race, puberty, in-scanner head motion, co-exposure to NO2 and O3, urbanicity at primary address. Random intercepts were modelled from MRI scanner device and subject ID.

#### RND in Females

In females, LMEs models revealed the same main effects found in males. However, there were no significant three-way interactions between any pollutant, subcortical RND and time on BMI, see **Supplemental Materials Tables S22-25.**

## 4. DISCUSSION

In a large and diverse cohort of youth, the present study found that exposure to outdoor air pollution coupled with developmental variations in brain microstructure across regions involved in appetite and decision-making may increase weight gain risk. Importantly, our study was conducted during the transition from late childhood to early adolescence, a critical temporal window of brain and cognitive development (Vijayakumar et al., 2018; Yurgelun-Todd, 2007). Specifically, our findings showed that greater exposure to outdoor PM2.5 at 9/10-years-old and having higher RND in right caudate nucleus, and bilateral putamen and pallidum was associated with greater weight gain by 11-12-years-old. However, our sex-stratified analyses suggest that PM2.5 exposure may be more consequential for males than females, but further research is needed to explore the mechanisms driving these sex differences. No other significant associations were observed between other pollutants, microstructure development, and BMI, including any observations with RNI. Although further research is needed, our findings suggest that obesity risk can be increased due to joint effects exerted by PM2.5 exposure and accelerated development of subcortical regions of the basal ganglia (e.g., caudate, putamen, pallidum) that support reward processing and decision making, two cognitive processes with an important role in food intake behaviors (Volkow et al., 2011).

The current findings showed that RND in reward-related regions (caudate, putamen, pallidum) were associated with higher BMI, but these associations were modified both by PM2.5 exposure and age. We hypothesized that air pollution exposure coupled with subcortical microstructural development (i.e., RNI, RND) would be associated with greater weight gain over time. In the case of RNI, we did not find a moderating effect of air pollution exposure on the relationship of this biomarker with BMI over time. However, we found that, regardless to air pollution exposure and age, higher RNI levels in all subcortical regions were associated with a higher BMI, with stable effects across development. This finding is in line with previous research showing that increased RNI is positively associated with higher BMI, which has been interpreted as suggestive neuroinflammation (Li et al., 2023). Thus, if RNI reflects microstructural alterations induced by neuroinflammation, alterations in subcortical regions involved in food intake may contribute to obesity risk with sustained effects across development. However, future studies should confirm this hypothesis using more reliable markers of neuroinflammation. Additionally, we also found that exposure to PM2.5 significantly moderated the positive association between RND and weight over time in subcortex. RND is a marker of neurite density and organization, and RND increases in subcortical regions is considered normative development during the transition from childhood to adolescence (Palmer et al., 2022). Thus, considering this evidence, our results suggest that accelerated subcortical development in regions of the basal ganglia coupled with higher PM2.5 exposure may contribute to weight gain over time as children transition to adolescence.

One possible underlying mechanism linking higher BMI with joint effects of PM2.5 exposure and accelerated subcortical development could involve earlier pubertal onset. Research indicates that children with obesity tend to experience an earlier onset of puberty (Song et al., 2023). Puberty is a period that initiates during late childhood that involves several endocrine changes that lead to attain sexual maturation and reproductive capability (Patton & Viner, 2007). This period also encompasses brain maturational processes characterized by multiple biological processes that change the cytoarchitecture of gray and white matter brain tissues (e.g., cell growth, myelination, synaptic pruning). Recent research has demonstrated that advances in pubertal development are associated with increased brain maturation (Holm et al., 2023), as indexed by brain age, a neuroimaging biomarker that reflects the biological age of the brain based on neuroimaging features (Cole & Franke, 2017). Thus, although our statistical models accounted for current pubertal status at each visit, inter-individual differences in the timing of pubertal onset and brain maturation before the temporal windows examined may still be present among the youth evaluated. These factors, in combination with neurotoxic effects induced by PM2.5 exposure, could contribute to BMI changes over time. However, this hypothesis should be addressed in further longitudinal research to disentangle the temporal order of these associations and their implications for childhood obesity risk.

From a neuroanatomical perspective, we found accelerated brain maturation in caudate nucleus, pallidum and putamen, three regions that play a key role in reward processing and decision-making behaviors (Hiebert et al., 2014; Smith et al., 2009) and functionally involved in cognitive process related to food intake control (Volkow et al., 2011). Considering our findings, altered development of these regions supporting reward processing and decision-making may increase food-seeking reward motivated behaviors. Thus, it is plausible that joint effects of PM2.5 exposure and accelerated maturation of brain regions that support these cognitive processes can lead to a higher impulsivity. Studies in animal models have shown some support for this, as exposure to concentrated ambient particulate matter was associated with an enhanced bias towards immediate rewards and higher levels of impulsivity (Allen et al., 2013). Moreover, the tendency to act impulsively has been shown to correlate with unhealthy eating patterns (Booth et al., 2018) and greater BMI in youth (Van Den Berg et al., 2011). Therefore, it may be plausible that air pollution exposure coupled with altered timing of brain development in reward processing structures may contribute to impulsive food choices, although further research is needed.

Importantly, our sex-stratified models suggest that the moderating effect of PM2.5 on the brain-weight relationship may be more pronounced in males. Previous research has shown sex-specific associations between brain structure and air pollution (Cotter et al., 2024; Peterson et al., 2022). However, these differences appear to depend on the specific air pollutant and the MRI metric evaluated. Consequently, due to inconsistent findings in the literature, the neurobiological factors that may place one sex at a higher risk than the other in response to air pollutant exposure remain unclear. This underscores the need for further research to disentangle the underlying mechanisms driving these differential effects. Additionally, sex differences in impulsive behaviors could help explain why the moderating effects of PM2.5 on the brain-weight relationship are more significant in males. While it is well established that males tend to display higher levels of impulsivity than females throughout development (Hosseini-Kamkar & Bruce Morton, 2014), this trait may interact differently with environmental factors like air pollution exposure. For instance, it has been demonstrated that exposure NO2 (Loftus et al., 2020) or PM2.5 (Smolker et al., 2024) is associated with increased externalizing behaviors in children like aggression, attention problems, and impulsivity in children, although more research is needed as these findings were not recently replicated (Campbell et al., 2024; Jorcano et al., 2019). Moreover, recent findings from the ABCD Study® revealed that, in males, fat and added sugar intake is strongly influenced by sensation-seeking impulsive behaviors (Adise, Boutelle, et al., 2024). Within this context, it is plausible that PM2.5 exposure coupled with altered development of basal ganglia regions involved in reward processing could exacerbate impulsivity behaviors in males, which could contribute to display unhealthy eating patterns.

This study has several strengths that contribute to advancing our understanding of how air pollution may influence the association between brain microstructure and weight gain in youth. With a large and diverse sample, this is the first longitudinal study to examine how air pollution exposure moderates the relationship between brain microstructural development in regions involved in food intake control and weight gain during the critical transition from late childhood to early adolescence. In addition to commonly studied pollutants (PM2.5, NO2, and O3), we also evaluated Oxwt, a metric reflecting the combined oxidative potential of NO2 and O3. Furthermore, we evaluated novel neuroimaging markers of cellular density (RNI) and neurite density and organization (RND), which may offer deeper neurobiological insights than macrostructural neuroimaging metrics (e.g., gray matter volume).

However, this study is not without limitations. All air pollutants evaluated in the current study were based on geocoded residential addresses of participants, using the annual average concentration from 2016. However, it should be noted that baseline data of the ABCD Study was collected between 2016 and 2018, when participants were 9-10 years old. Therefore, there might be some temporal mismatch between air pollution estimates and the timing of data collection for each participant, which could compromise the precision of our findings. Unfortunately, air pollutant estimates for subsequent visits are not available in the latest ABCD data release, which limits our ability to evaluate how changes in exposure relate to microstructural alterations over time. Nevertheless, it is worth noting that prior research has shown that annual averages of PM2.5 remain relatively stable spatially in the years leading up to 2016 (Di et al., 2019). Incorporating air pollution data from before and after the evaluated time window (9 to 12 years old) would provide insights into ongoing and cumulative exposure, helping to explain individual differences in brain development and obesity risk. Additionally, air pollution estimates evaluated in the present study were limited to participants’ primary residential addresses due to a high rate of missing data from additional reported addresses. Future studies should include exposure measures from other areas where participants spend significant time, such as schools. Moreover, due to the inclusion of participants with overweight and obesity at baseline, the current study does not allow us to establish the causal mechanisms underlying the moderating effects of air pollution on the relationship between brain microstructure and the development of unhealthy weight. Despite this, our findings advance the current understanding of how exposure to air pollution could interact with altered neurodevelopment of brain regions involved in regulating food intake behaviors, and how these joint associations may then contribute to overweight and obesity. Finally, the effect sizes in this study were small, but this aligns with expectations in large-scale studies evaluating brain development (Owens et al., 2021).

## 5. CONCLUSIONS

In summary, this study provides some support that exposure to PM2.5 coupled with accelerated brain development in basal ganglia regions involved in appetite and decision-making (caudate nucleus, putamen, and pallidum) may increase weight gain risk from ages 9 to 12 years-old. Importantly, the moderating effects of PM2.5 exposure on the relationship between brain development and weight gain are more consequential for males, highlighting potential sex-specific environmental and neurobiological susceptibilities that should be jointly evaluated in future research. Given the rising burden of both childhood obesity and environmental pollution, further multidisciplinary longitudinal research is essential to uncover with higher specificity the underlying mechanisms driving these associations. This knowledge will be important for developing targeted public health interventions aimed at mitigating these environmental health risks at critical windows of brain and cognitive development, which would be important for preventing long-term health effects at later lifespan stages. A deeper understanding of these associations will contribute to a more resilient, sustainable, and healthy future.

## Data availability

Data used in the preparation of this article were obtained from the ABCD Study® (https://abcdstudy.org/), which is held in the NIMH Data Archive (NDA) and is available for any researcher wishing to use it. The ABCD Study® data repository grows and changes over time. The ABCD Study® data used in this report were obtained from https://dx.doi.org/10.15154/z563-zd24. The code used to prepare the data, perform the analysis, and plot the results can be found in (https://github.com/Adise-lab/abcd-air-pollution-bmi)

## Acknowledgements

We would like to thank the participants of the ABCD Study® and the research assistants who collected the data.

Funding acknowledgements: The data used in the preparation of this article were obtained from the Adolescent Brain Cognitive Development^SM^ Study® (https://abcdstudy.org/), which is held in the NIMH Data Archive (NDA). The ABCD Study^®^ is supported by the National Institutes of Health and National Institute on Drug Abuse and additional federal partners under award numbers U01DA041048, U01DA050989, U01DA051016, U01DA041022, U01DA051018, U01DA051037, U01DA050987, U01DA041174, U01DA041106, U01DA041117, U01DA041028, U01DA041134, U01DA050988, U01DA051039, U01DA041156, U01DA041025, U01DA041120, U01DA051038, U01DA041148, U01DA041093, U01DA041089, U24DA041123, U24DA041147. A full list of supporters is available at https://abcdstudy.org/federal-partners/. A listing of participating sites and a complete listing of the study investigators can be found at https://abcdstudy.org/principal-investigators/. The ABCD Study^®^ consortium investigators designed and implemented the study and/or provided data but did not necessarily participate in analysis or writing of this report. The ABCD Study^®^ data repository grows and changes over time. The ABCD Study® data used in this report were obtained from https://dx.doi.org/10.15154/z563-zd24. Additional support for this work was made possible from National Institute of Environmental Health Sciences (NIEHS): R01-ES032295 and R01-ES031074. Additionally, SA was supported by funding from the National Institutes of Health: National Institute of Diabetes and Digestive and Kidney Diseases (NIH NIDDK) (K01 DK135847), The Southern California Center for Latino Health (Funded by The National Institute on Minority Health and Health Disparities, P50MD017344) and funding by The Saban Research Institute at Children’s Hospital of Los Angeles. JOG was supported by The Saban Research Institute at Children’s Hospital Los Angeles (RCDF-2024-000016305). This manuscript reflects the views of the authors and may not reflect the opinions or views of the NIH, Southern California Center for Latino Health, The Saban Research Institute or other ABCD Study^®^ consortium investigators.

