## Supplemental Materials for "Sex-specific Effects of Outdoor Air Pollution on Subcortical Microstructure and Weight Gain: Findings from the ABCD Study®"

| | |
|---|---|
| ● **Supplemental Methods** | Pg 3-6 |
| ● **Supplemental Tables** | Pg 7-31 |
| ● Table S1: Exclusion criteria | Pg 7 |
| ● Tables S2: Main effects and interactions of linear mixed effects models assessing moderating effects of PM2.5 on the relationship between RNI and BMI across both sexes. | Pg 8 |
| ● Tables S3: Main effects and interactions of linear mixed effects models assessing moderating effects of NO2 on the relationship between RNI and BMI across both sexes. | Pg 9 |
| ● Tables S4: Main effects and interactions of linear mixed effects models assessing moderating effects of O3 on the relationship between RNI and BMI across both sexes. | Pg 10 |
| ● Tables S5: Main effects and interactions of linear mixed effects models assessing moderating effects of Oxwt on the relationship between RNI and BMI across both sexes. | Pg 11 |
| ● Tables S6: Main effects and interactions of linear mixed effects models assessing moderating effects of PM2.5 on the relationship between RNI and BMI in males. | Pg 12 |
| ● Tables S7: Main effects and interactions of linear mixed effects models assessing moderating effects of NO2 on the relationship between RNI and BMI in males. | Pg 13 |
| ● Tables S8: Main effects and interactions of linear mixed effects models assessing moderating effects of O3 on the relationship between RNI and BMI in males. | Pg 14 |
| ● Tables S9: Main effects and interactions of linear mixed effects models assessing moderating effects of Oxwt on the relationship between RNI and BMI in males. | Pg 15 |
| ● Tables S10: Main effects and interactions of linear mixed effects models assessing moderating effects of PM2.5 on the relationship between RNI and BMI in females. | Pg 16 |
| ● Tables S11: Main effects and interactions of linear mixed effects models assessing moderating effects of NO2 on the relationship between RNI and BMI in females. | Pg 17 |
| ● Tables S12: Main effects and interactions of linear mixed effects models assessing moderating effects of O3 on the relationship between RNI and BMI in females. | Pg 18 |
| ● Tables S13: Main effects and interactions of linear mixed effects models assessing moderating effects of Oxwt on the relationship between RNI and BMI in females. | Pg 19 |
| ● Tables S14: Main effects and interactions of linear mixed effects models assessing moderating effects of PM2.5 on the relationship between RND and BMI across both sexes. | Pg 20 |
| ● Tables S15: Main effects and interactions of linear mixed effects models assessing moderating effects of NO2 on the relationship between RND and BMI across both sexes. | Pg 21 |
| ● Tables S16: Main effects and interactions of linear mixed effects models assessing moderating effects of O3 on the relationship between RND and BMI across both sexes. | Pg 22 |

| | |
|---|---|
| • Tables S17: Main effects and interactions of linear mixed effects models assessing moderating effects of Oxwt on the relationship between RND and BMI across both sexes. | Pg 23 |
| • Tables S18: Main effects and interactions of linear mixed effects models assessing moderating effects of PM2.5 on the relationship between RND and BMI in males. | Pg 24 |
| • Tables S19: Main effects and interactions of linear mixed effects models assessing moderating effects of NO2 on the relationship between RND and BMI in males. | Pg 25 |
| • Tables S20: Main effects and interactions of linear mixed effects models assessing moderating effects of O3 on the relationship between RND and BMI in males. | Pg 26 |
| • Tables S21: Main effects and interactions of linear mixed effects models assessing moderating effects of Oxwt on the relationship between RND and BMI in males. | Pg 27 |
| • Tables S22: Main effects and interactions of linear mixed effects models assessing moderating effects of PM2.5 on the relationship between RNI and BMI in females. | Pg 28 |
| • Tables S23: Main effects and interactions of linear mixed effects models assessing moderating effects of NO2 on the relationship between RNI and BMI in females. | Pg 29 |
| • Tables S24: Main effects and interactions of linear mixed effects models assessing moderating effects of O3 on the relationship between RNI and BMI in females. | Pg 30 |
| • Tables S25: Main effects and interactions of linear mixed effects models assessing moderating effects of Oxwt on the relationship between RNI and BMI in females. | Pg 31 |
| • **Supplemental References** | Pg 32-33 |

### Supplemental Methods

#### Exclusion criteria

The number of subjects available at baseline and year 2 after applying each exclusion criteria detailed below are available in **Supplementary Table 1**.

Participants were excluded if they had major medical or neurological conditions, psychiatric disorders, intellectual disabilities, and alcohol or substance abuse. Moreover, to avoid potential prenatal influences in brain development caused by maternal drug exposure (Thompson et al., 2009), participants were also excluded if their biological mothers reported consumption of alcohol or other drugs after becoming aware of the pregnancy (if data were available). Further exclusion criteria included a history of traumatic head injury to avoid the influence of such injuries on brain structure (Tasker, 2006), intersex status, or the use of medications that could affect weight (e.g., insulin, anti-hyperglycaemic drugs, or weight loss medications). To avoid potential differences in brain development and body growth due to prematurity (Ji et al., 2024), participants born more than 12 weeks premature, or with a birth weight less than 1,200 grams were excluded.

A quality control assessment was performed on the anthropometric data to identify inconsistencies. Height measurements were compared across baseline, first year, and second year follow-ups. Participants with implausible height measurements were excluded (i.e., participant's follow-up height less than the previous year; not plausible growth rates over time [e.g., more than 8 inches in a single year]). Participants with weight loss of more than 20 kilograms or with implausible weight (i.e., very low weight) were flagged, subsequently reviewed for potential errors, and excluded as appropriate. Moreover, participants with underweight during baseline and follow-up were excluded to prevent potential issues with restrictive feeding or medical issues. To avoid inaccuracies associated with at-home estimates of anthropometrics, participants with remote assessments were excluded. Additionally, participants with missing data, or those whose visits occurred during COVID-19 restrictions (before 2020/03/19) were also excluded.

Regarding MRI data, participants were excluded if there were image artifacts preventing radiology reading or if incidental findings were reported by a Board-Certified Neuroradiologist. The ABCD Data Analysis, Informatics, and Resource Center (DAIRC) were responsible for identifying images that contained artifacts, clinical findings, or poor quality after processing. This center managed the preprocessing and quality control of the data and provided exclusion recommendations. Participants were excluded under the following conditions: raw DWI and T1 data did not pass initial quality control assessments; there was no available B0 unwarp data; incomplete DWI scans; errors occurred in registering DWI to the T1w structural images; there were failures in FreeSurfer and DWI post-processing; the dorsal cut-off score was higher than 47 and the ventral cut-off score higher than 54; or derived results were unavailable. This quality control criterion was coded in the ABCD dictionary as imgincl_dmri_include. In addition to this quality control assessment, tabulated data in each brain region was inspected to detect outliers (i.e., three standard deviations above or below the mean). Moreover, participants were also excluded if their head motion during the DWI acquisition, measured as the mean framewise displacement, exceeded 1.7 mm. Additionally, to avoid brain structure alterations related to COVID-19 (van Drunen et al., 2023), participants who underwent the MRI session after pandemic restrictions were excluded. Participants with missing data on age, sex, education, MRI scanner type, urbanicity, PM2.5, NO2, O3, number of years living at address 1, percentage of time spent at address 1, or those who reported implausible data on the percentage of time children spent at the primary address (e.g., 1% of time) were also excluded. In addition, participants who had data available at year 2 but not at baseline were excluded. As some of the collecting sites from the ABCD study oversampled siblings in the recruitment process, and to avoid issues with independence, the current statistical analyses only included one participant from each family, selected at random but prioritizing participants who had data available at both baseline and follow-up to maximize the sample size.

**Physical assessments**

Height and weight measurements were assessed by a trained researcher. Participants were weighed to the nearest 0.1 lb while wearing light clothing and stocking feet, using a physician's scale (Detecto model 439, Webb City, MO). Height was recorded on the same scale, accurate to the nearest 0.1 inch. Each measurement was taken twice and averaged, with a third measurement added if there was a significant difference between the first two.

**Pubertal assessment.**

At each visit, both the attending caregiver and the youth completed the Pubertal Development Scale (PDS) (Petersen et al., 1988). This questionnaire consisted of sex-specific versions that assessed physical changes associated with puberty across 5 questions including growth in height, body hair, and whether they noticed skin changes. The male-specific form included questions about deepening of the voice and facial hair growth. The form for females included questions about breast development and menstruation (e.g., has it begun? at what age did menstruation begin?). Response options were: 1) not yet started; 2) barely started; 3) definitely started; 4) seems complete; 5) I do not know (or missing). Menstruation responses were coded as yes and no, where yes corresponded to 4 points (no 1 point). A puberty score was calculated using the sum of the responses.

Puberty scores for boys consisted of adding the scores for body hair growth, deepening of the voice, and facial hair. Scores were converted into Tanner staging as follows: Pre-pubescent = 3; early puberty = >3 and <6; mid puberty = >5 and <9; late puberty >8 and <12; post puberty ≥ 12. Puberty scores for females were scored using body hair growth, breast development and menarche. Scores were converted into Tanner staging as follows: pre-pubescent =3; early puberty =3 and no menarche; mid puberty = 4 and < 8 and no menarche; late puberty <8 and menarche; post puberty 8 and menarche.

Participants in which missing values for any of the items needed to calculate Tanner staging, were excluded. Because not all caregivers are aware of the youth's pubertal growth, and that youth may overestimate, an average of the caregiver and youth tanner stage (at each time point, respectively was utilized). In rare cases, Tanner staging only existed for one responder (e.g., caregiver, youth), so no average existed as Tanner staging was based on only one report.

#### Magnetic Resonance Imaging acquisition and preprocessing

Magnetic Resonance Imaging (MRI) data were acquired with 3T MRI scanners across 21 sites ($n_{scanners} = 31$). Brain structural imaging data were collected through T1-weighted and multi-shell Diffusion Weighted Imaging (DWI) sequences. A detailed description of the preprocessing pipeline employed can be found in **Supplemental Materials**. The preprocessing pipeline of DWI data included eddy current (Zhuang et al., 2006) and head motion correction (Hagler et al., 2019) adjustment of gradients for head motion (Hagler et al., 2019; Leemans and Jones, 2009), robust tensor fit to identify and exclude dark frames due to abrupt head motion (Chang et al., 2005), B0 distortion and gradient distortion correction using opposite phase encoding pairs of b=0s (Holland et al., 2010; Jovicich et al., 2006), b0 registration to T1w structural images using mutual information (Wells et al., 1996) and cubic interpolation to resample at the 1.7 isotropic resolution. The ABCD DAIRC was responsible for the preprocessing of these data. The multiple b-value DWI acquisition allowed the estimation of an RSI model (White et al., 2013, 2014). A linear estimation approach was employed to estimate mixtures of “restricted” and “hindered” diffusion within individual voxels. The RSI to model was employed to estimate two volume fractions, representing intracellular (restricted) and extracellular (hindered) diffusion, with separate fibre orientation density (FOD) functions, modeled as fourth-order spherical harmonic functions, allowing for multiple diffusion orientations within a single voxel. The longitudinal diffusion was held constant for both fractions, with a value of $1\times10^{-3}$ mm$^2$/s. For the restricted fraction, the transverse diffusion was modeled as 0, and for the hindered fraction, the transverse diffusion was modeled as $0.9 \times 10^{-3}$ mm$^2$/s. The quantification of the restricted normalized diffusion provides biological insights about the structure of the brain tissues such as the cellular and neurite density within grey matter tissues.

#### Outdoor Air pollution estimates

Caregivers provided their residential addresses, which were used to link to annual average estimates of air pollution (Fan et al., 2021). Annual average air pollution estimates were collected in 2016, which corresponded to the start of the ABCD Study enrolment (i.e., when children were 9-10 years old). Air pollution data was estimated daily over the 2016 calendar year at a 1-km$^2$ resolution using hybrid spatiotemporal models that combine satellite-based aerosol optical depth models, land-use regression, and chemical transport model outputs (Di et al., 2019; Requia et al., 2020). Additionally, the DAIRC geocoded the latitude and longitude of baseline residential addresses, with up to three residential addresses collected for each participant, using the Google Maps Application Programming Interface (Fan et al., 2021). Urbanicity of participant’s addresses was assessed with a categorical variable indicating if participants’ addresses were in census tracts considered to be urban (50,000 or more people), urban clusters (at least 2,500 and less than 50,000 people), or rural (less than 2,500 people per tract, and/or not included in urban areas or clusters). These categories were established according to the Census Tract Urban Classification and were based on 2010 census estimates (Fan et al., 2021). The annual average pollution concentrations were linked to the youth’s geocoded residential addresses. Daily exposure estimates were calculated over the 2016 calendar period for fine particulate matter (PM2.5) in micrograms per cubic meter (μg/m$^3$), nitrogen dioxide (NO2) in parts per billion (ppb), and 8-hour maximum ozone (O3) levels in ppb, all at a 1 km$^2$ resolution. Due to their potential to induce oxidative stress when combined, the redox-weighted oxidative capacity ($O_x^{wt}$) was estimated as the ratio of the weighted redox potentials of NO2 and O3 using the following formula: [(1.07 * NO2) + (2.075 * O3)] / 3.145 (Williams et al., 2014). Further information about the methods and protocols employed to estimate air pollution data in the ABCD Study® are described elsewhere (Badilla et al., 2024; Fan et al., 2021). Given the high percentage of missing data for pollutants associated with the second and third addresses in the current sample (address 2: 89.32%; address 3: 98.19%), this study focused exclusively on the primary residential address.

**Associations between air pollutants and BMI.**

The relationship between BMI and each air pollutant (i.e., PM2.5, NO2, O3, and Oxwt) was evaluated across sexes and also in each sex by means of linear mixed effects models (LME) adjusted for household education, race, ethnicity, puberty and urbanicity according to the Census Tract Urban Classification. To account also for co-exposure, models for PM2.5 additionally included NO2 and O3 as covariates; models for NO2 included PM2.5 and O3; models for O3 included PM2.5 and NO2; and models for Oxwt included PM2.5. To control the potential influence of site effects, site and participants identifiers (IDs) were included in the model as crossed random effects. The model formula employed were the following:

**PM2.5**

BMI ~ PM2.5 * age + sex + caregiver highest education + race + ethnicity + puberty + NO2 + O3 + urbanicity at primary address + (1| Site_ID) + (1| subject_ID)

**NO2**

BMI ~ NO2 * age + sex + caregiver highest education + race + ethnicity + puberty + PM2.5 + O3 + urbanicity at primary address + (1| Site_ID) + (1| subject_ID)

**O3**

BMI ~ O3 * age + sex + caregiver highest education + race + ethnicity + puberty + PM2.5 + NO2 + urbanicity at primary address + (1| Site_ID) + (1| subject_ID)

**Oxwt**

BMI ~ Oxwt * age + sex + caregiver highest education + race + ethnicity + puberty + PM2.5 + urbanicity at primary address + (1| Site_ID) + (1| subject_ID)

**Note:** Sex-stratified models were the same but without the sex covariate. Results of these analyses can be found in the main manuscript.

### Supplemental Tables

**Supplementary Table 1.** The number of subjects available at baseline and year 2 after applying each exclusion criteria.

| Criteria | Description | n |
|---|---|---|
| 1 | Medical conditions, psychiatric disorders, drugs, head injuries, intersex, medications, gestational age, low birthweight | Baseline: 9193 |
| | | Year 2: 8502 |
| 2 | Anthropometric data: shrink cases, wrong data in growth, excessive weight loss, missing data, visits conducted remotely or during COVID-19 restrictions. | Baseline: 9073 |
| | | Year 2: 7155 |
| 3 | Exclude underweight in both visits. | Baseline: 8745 |
| | | Year 2: 5599 |
| 4 | Imaging artifacts, incidental findings, missing data in head motion or excessive head motion during MRI acquisition. | Baseline: 6535 |
| | | Year 2: 4033 |
| 5 | MRI acquisition during COVID-19 restrictions. | Baseline: 6535 |
| | | Year 2: 4033 |
| 6 | Missing data: age, sex, education, MRI scanner, race, ethnicity, puberty, urbanicity, PM2.5, NO2, O3. | Baseline: 5369 |
| | | Year 2: 3404 |
| 7 | Exclude participants in year 2 who do not have data at baseline | Baseline: 5369 |
| | | Year 2: 2652 |
| 8 | No siblings | **Baseline: 4802** |
| | | **Year 2: 2439** |

**Supplementary Table 2.** Main effects and interactions for the PM2.5 pollutant across both sexes in RNI.

| | | BMI ~ $ROI_{RNI}$ * PM2.5 * age + sex + caregiver highest education + race + ethnicity + puberty + NO2 + O3 + head motion + urbanicity address 1 + (1\|MRI device) + (1\| Subject ID) | | | | | | | | | | | | |
|---|---|---|---|---|---|---|---|---|---|---|---|---|---|---|
| | | | **$ROI_{RNI}$** | | **PM2.5** | | **Age** | | **$ROI_{RNI}$ X PM2.5** | | **$ROI_{RNI}$ x Age** | | **$ROI_{RNI}$ x PM2.5 x Age** | |
| **ROI** | **H** | **n** | **β (95%CI)** | **p-value** | **β (95%CI)** | **p-value** | **β (95%CI)** | **p-value** | **β (95%CI)** | **p-value** | **β (95%CI)** | **p-value** | **β (95%CI)** | **p-value** |
| **Thalamus** | L | 7073 | 0.186 (0.098, 0.274) | <0.001 | 0.120 (-0.028, 0.267) | 0.113 | 0.808 (0.747, 0.869) | <0.001 | 0.002 (-0.079, 0.084) | 0.954 | -0.020 (-0.063, 0.023) | 0.357 | 0.005 (-0.033, 0.044) | 0.795 |
| | R | 7099 | 0.226 (0.139, 0.314) | <0.001 | 0.101 (-0.053, 0.255) | 0.198 | 0.794 (0.733, 0.856) | <0.001 | 0.049 (-0.031, 0.130) | 0.229 | -0.017 (-0.061, 0.027) | 0.442 | 0.030 (-0.009, 0.070) | 0.129 |
| **Caudate** | L | 7166 | 0.297 (0.228, 0.366) | <0.001 | 0.135 (-0.007, 0.276) | 0.064 | 0.781 (0.723, 0.839) | <0.001 | 0.048 (-0.019, 0.116) | 0.158 | 0.044 (0.003, 0.086) | 0.038 | -0.031 (-0.073, 0.010) | 0.140 |
| | R | 7151 | 0.193 (0.123, 0.262) | <0.001 | 0.117 (-0.026, 0.260) | 0.111 | 0.809 (0.751, 0.867) | <0.001 | 0.011 (-0.055, 0.078) | 0.741 | 0.031 (-0.012, 0.073) | 0.155 | 0.008 (-0.034, 0.049) | 0.710 |
| **Putamen** | L | 7142 | 0.353 (0.269, 0.438) | <0.001 | 0.119 (-0.033, 0.270) | 0.126 | 0.715 (0.651, 0.780) | <0.001 | 0.013 (-0.061, 0.087) | 0.731 | 0.029 (-0.013, 0.071) | 0.172 | -0.019 (-0.060, 0.021) | 0.345 |
| | R | 7149 | 0.359 (0.278, 0.440) | <0.001 | 0.113 (-0.033, 0.259) | 0.132 | 0.704 (0.640, 0.769) | <0.001 | -0.016 (-0.088, 0.055) | 0.654 | 0.028 (-0.014, 0.070) | 0.189 | -0.006 (-0.048, 0.036) | 0.786 |
| **Pallidum** | L | 7085 | 0.348 (0.254, 0.442) | <0.001 | 0.102 (-0.049, 0.253) | 0.186 | 0.707 (0.640, 0.775) | <0.001 | 0.019 (-0.061, 0.098) | 0.649 | 0.004 (-0.037, 0.046) | 0.837 | -0.008 (-0.045, 0.028) | 0.657 |
| | R | 7085 | 0.384 (0.289, 0.480) | <0.001 | 0.107 (-0.048, 0.261) | 0.179 | 0.696 (0.628, 0.764) | <0.001 | 0.056 (-0.027, 0.139) | 0.186 | -0.011 (-0.053, 0.030) | 0.598 | -0.011 (-0.047, 0.026) | 0.559 |
| **Hippocampus** | L | 7137 | 0.180 (0.105, 0.255) | <0.001 | 0.150 (0.005, 0.295) | 0.045 | 0.812 (0.753, 0.871) | <0.001 | -0.019 (-0.089, 0.052) | 0.606 | -0.015 (-0.059, 0.029) | 0.506 | -0.012 (-0.056, 0.032) | 0.585 |
| | R | 7150 | 0.203 (0.124, 0.281) | <0.001 | 0.170 (0.028, 0.313) | 0.020 | 0.801 (0.741, 0.862) | <0.001 | -0.006 (-0.080, 0.068) | 0.873 | -0.020 (-0.062, 0.023) | 0.358 | 0.006 (-0.034, 0.047) | 0.760 |
| **Amygdala** | L | 7126 | 0.050 (-0.022, 0.121) | 0.176 | 0.162 (0.022, 0.302) | 0.025 | 0.858 (0.799, 0.916) | <0.001 | -0.022 (-0.089, 0.046) | 0.525 | -0.039 (-0.083, 0.005) | 0.083 | -0.010 (-0.052, 0.032) | 0.637 |
| | R | 7130 | 0.035 (-0.040, 0.109) | 0.364 | 0.187 (0.049, 0.324) | 0.009 | 0.860 (0.801, 0.919) | <0.001 | -0.065 (-0.134, 0.003) | 0.063 | -0.044 (-0.087, -0.001) | 0.047 | 0.006 (-0.032, 0.044) | 0.756 |
| **Accumbens** | L | 7111 | 0.455 (0.373, 0.536) | <0.001 | 0.073 (-0.080, 0.227) | 0.347 | 0.714 (0.653, 0.774) | <0.001 | -0.023 (-0.094, 0.049) | 0.532 | 0.044 (0.001, 0.087) | 0.047 | -0.037 (-0.080, 0.005) | 0.084 |
| | R | 7106 | 0.372 (0.292, 0.452) | <0.001 | 0.101 (-0.048, 0.250) | 0.186 | 0.735 (0.675, 0.796) | <0.001 | -0.017 (-0.088, 0.054) | 0.638 | 0.044 (0.002, 0.086) | 0.041 | -0.038 (-0.078, 0.003) | 0.068 |
| **Ventral DC** | L | 7087 | 0.182 (0.098, 0.267) | <0.001 | 0.157 (0.012, 0.303) | 0.036 | 0.819 (0.759, 0.878) | <0.001 | -0.052 (-0.126, 0.022) | 0.167 | -0.033 (-0.077, 0.010) | 0.133 | -0.001 (-0.037, 0.034) | 0.945 |
| | R | 7101 | 0.168 (0.083, 0.252) | <0.001 | 0.157 (0.012, 0.303) | 0.036 | 0.826 (0.767, 0.885) | <0.001 | -0.035 (-0.110, 0.040) | 0.361 | -0.038 (-0.081, 0.006) | 0.089 | 0.005 (-0.031, 0.041) | 0.780 |

**Note:** Estimated regression coefficients (β), 95% confidence intervals (CI) and *p*-values from the linear mixed-effects models evaluating the moderating effects of PM2.5 in the relationship between brain microstructure and Body Mass Index. Models were controlled for age, sex, education, race, ethnicity, puberty, co-exposure to other air pollutants, head motion during the MRI acquisition and urbanicity. Random intercepts were modelled from MRI scanner device and subject ID. Although all two-way interactions were assessed, this table presents only the ones of primary interest due to space constraints. Abbreviations: ROI: region-of-interest; Ventral DC: ventral diencephalon; RSI: restricted spectrum imaging; RNI: restricted normalized isotropic diffusion; H: hemisphere. *p-values < 0.05 of two-way and three-way interactions that achieved the Benjamini-Hochberg False Discovery Rate threshold correction.

**Supplementary Table 3.** Main effects and interactions for the NO2 pollutant across both sexes in RNI.

| | | BMI ~ $ROI_{RNI}$ * NO2 * age + sex + caregiver highest education + race + ethnicity + puberty + PM2.5 + O3 + head motion + urbanicity address 1 + (1\|MRI device) + (1\| Subject ID) | | | | | | | | | | | | |
|---|---|---|---|---|---|---|---|---|---|---|---|---|---|---|
| | | | $ROI_{RNI}$ | | NO2 | | Age | | $ROI_{RNI}$ X NO2 | | $ROI_{RNI}$ x Age | | $ROI_{RNI}$ x NO2 x Age | |
| **ROI** | **H** | **n** | **β (95%CI)** | **p-value** | **β (95%CI)** | **p-value** | **β (95%CI)** | **p-value** | **β (95%CI)** | **p-value** | **β (95%CI)** | **p-value** | **β (95%CI)** | **p-value** |
| **Thalamus** | L | 7073 | 0.191 (0.103, 0.279) | <0.001 | 0.078 (-0.072, 0.229) | 0.308 | 0.806 (0.745, 0.867) | <0.001 | 0.025 (-0.064, 0.114) | 0.580 | -0.018 (-0.062, 0.025) | 0.413 | 0.015 (-0.028, 0.058) | 0.486 |
| | R | 7099 | 0.239 (0.152, 0.325) | <0.001 | 0.099 (-0.056, 0.254) | 0.213 | 0.790 (0.729, 0.851) | <0.001 | 0.060 (-0.027, 0.146) | 0.178 | -0.015 (-0.059, 0.029) | 0.510 | 0.020 (-0.023, 0.063) | 0.365 |
| **Caudate** | L | 7166 | 0.295 (0.226, 0.365) | <0.001 | 0.064 (-0.081, 0.210) | 0.389 | 0.773 (0.716, 0.831) | <0.001 | 0.016 (-0.053, 0.085) | 0.645 | 0.048 (0.007, 0.090) | 0.023 | -0.052 (-0.094, -0.011) | 0.014 |
| | R | 7151 | 0.193 (0.124, 0.262) | <0.001 | 0.070 (-0.076, 0.217) | 0.348 | 0.807 (0.749, 0.865) | <0.001 | 0.015 (-0.055, 0.085) | 0.678 | 0.033 (-0.010, 0.075) | 0.131 | -0.028 (-0.069, 0.013) | 0.187 |
| **Putamen** | L | 7142 | 0.376 (0.291, 0.461) | <0.001 | 0.102 (-0.053, 0.257) | 0.200 | 0.707 (0.643, 0.771) | <0.001 | 0.107 (0.023, 0.191) | 0.012 | 0.030 (-0.012, 0.071) | 0.157 | -0.0001 (0.040, 0.040) | 0.996 |
| | R | 7149 | 0.367 (0.287, 0.448) | <0.001 | 0.095 (-0.055, 0.246) | 0.217 | 0.704 (0.640, 0.768) | <0.001 | 0.064 (-0.018, 0.146) | 0.124 | 0.032 (-0.010, 0.074) | 0.139 | -0.013 (-0.053, 0.027) | 0.529 |
| **Pallidum** | L | 7085 | 0.367 (0.273, 0.461) | <0.001 | 0.083 (-0.072, 0.238) | 0.293 | 0.700 (0.633, 0.768) | <0.001 | 0.125 (0.033, 0.218) | 0.008 | 0.006 (-0.035, 0.048) | 0.760 | 0.017 (-0.025, 0.058) | 0.435 |
| | R | 7085 | 0.396 (0.300, 0.491) | <0.001 | 0.113 (-0.043, 0.270) | 0.158 | 0.691 (0.623, 0.759) | <0.001 | 0.070 (-0.023, 0.163) | 0.138 | -0.011 (-0.053, 0.031) | 0.596 | 0.016 (-0.024, 0.056) | 0.435 |
| **Hippocampus** | L | 7137 | 0.187 (0.113, 0.261) | <0.001 | 0.074 (-0.077, 0.224) | 0.338 | 0.810 (0.751, 0.869) | <0.001 | 0.014 (-0.065, 0.093) | 0.724 | -0.017 (-0.061, 0.027) | 0.448 | -0.004 (-0.049, 0.040) | 0.845 |
| | R | 7150 | 0.208 (0.131, 0.286) | <0.001 | 0.060 (-0.086, 0.207) | 0.420 | 0.798 (0.738, 0.859) | <0.001 | -0.018 (-0.100, 0.063) | 0.661 | -0.022 (-0.065, 0.020) | 0.307 | 0.018 (-0.024, 0.060) | 0.403 |
| **Amygdala** | L | 7126 | 0.060 (-0.011, 0.131) | 0.099 | 0.043 (-0.102, 0.188) | 0.563 | 0.857 (0.798, 0.916) | <0.001 | 0.060 (-0.013, 0.132) | 0.108 | -0.041 (-0.085, 0.003) | 0.068 | 0.003 (-0.042, 0.049) | 0.881 |
| | R | 7130 | 0.033 (-0.042, 0.107) | 0.389 | 0.042 (-0.101, 0.185) | 0.565 | 0.856 (0.797, 0.915) | <0.001 | -0.030 (-0.110, 0.049) | 0.456 | -0.044 (-0.088, -0.001) | 0.047 | 0.003 (-0.040, 0.046) | 0.884 |
| **Accumbens** | L | 7111 | 0.460 (0.379, 0.541) | <0.001 | 0.125 (-0.031, 0.282) | 0.117 | 0.707 (0.647, 0.767) | <0.001 | 0.069 (-0.011, 0.149) | 0.092 | 0.043 (0.0001, 0.086) | 0.049 | -0.027 (-0.071, 0.016) | 0.217 |
| | R | 7106 | 0.380 (0.300, 0.460) | <0.001 | 0.094 (-0.059, 0.247) | 0.230 | 0.728 (0.668, 0.789) | <0.001 | 0.047 (-0.032, 0.126) | 0.242 | 0.044 (0.002, 0.086) | 0.041 | -0.013 (-0.054, 0.028) | 0.530 |
| **Ventral DC** | L | 7087 | 0.190 (0.105, 0.275) | <0.001 | 0.077 (-0.073, 0.227) | 0.316 | 0.820 (0.760, 0.879) | <0.001 | 0.039 (-0.045, 0.124) | 0.364 | -0.028 (-0.073, 0.017) | 0.226 | 0.023 (-0.022, 0.069) | 0.312 |
| | R | 7101 | 0.174 (0.089, 0.258) | <0.001 | 0.069 (-0.080, 0.219) | 0.365 | 0.825 (0.766, 0.884) | <0.001 | 0.007 (-0.078, 0.092) | 0.866 | -0.035 (-0.080, 0.010) | 0.125 | 0.016 (-0.028, 0.060) | 0.470 |

**Note:** Estimated regression coefficients (β), 95% confidence intervals (CI) and *p*-values from the linear mixed-effects models evaluating the moderating effects of NO2 in the relationship between brain microstructure and Body Mass Index. Models were controlled for age, sex, education, race, ethnicity, puberty, co-exposure to other air pollutants, head motion during the MRI acquisition and urbanicity. Random intercepts were modelled from MRI scanner device and subject ID. Although all two-way interactions were assessed, this table presents only the ones of primary interest due to space constraints. Abbreviations: ROI: region-of-interest; Ventral DC: ventral diencephalon; RSI: restricted spectrum imaging; RNI: restricted normalized isotropic diffusion; H: hemisphere. *p-values < 0.05 of two-way and three-way interactions that achieved the Benjamini-Hochberg False Discovery Rate threshold correction.

**Supplementary Table 4.** Main effects and interactions for the O3 pollutant across both sexes in RNI.

| | | BMI ~ $ROI_{RNI}$ * O3 * age + sex + caregiver highest education + race + ethnicity + puberty + PM2.5 + NO2 + head motion + urbanicity address 1 + (1\|MRI device) + (1\| Subject ID) | | | | | | | | | | | | |
|---|---|---|---|---|---|---|---|---|---|---|---|---|---|---|
| | | | **$ROI_{RNI}$** | | **O3** | | **Age** | | **$ROI_{RNI}$ X O3** | | **$ROI_{RNI}$ x Age** | | **$ROI_{RNI}$ x O3 x Age** | |
| **ROI** | **H** | **n** | **β (95%CI)** | **p-value** | **β (95%CI)** | **p-value** | **β (95%CI)** | **p-value** | **β (95%CI)** | **p-value** | **β (95%CI)** | **p-value** | **β (95%CI)** | **p-value** |
| **Thalamus** | L | 7073 | 0.186 (0.099, 0.274) | <0.001 | 0.002 (-0.121, 0.125) | 0.977 | 0.804 (0.743, 0.865) | <0.001 | -0.009 (-0.095, 0.077) | 0.837 | -0.022 (-0.066, 0.021) | 0.312 | 0.011 (-0.033, 0.055) | 0.624 |
| | R | 7099 | 0.226 (0.139, 0.312) | <0.001 | -0.003 (-0.127, 0.122) | 0.963 | 0.791 (0.730, 0.852) | <0.001 | -0.046 (-0.128, 0.037) | 0.278 | -0.018 (-0.062, 0.026) | 0.417 | -0.012 (-0.057, 0.032) | 0.594 |
| **Caudate** | L | 7166 | 0.294 (0.225, 0.363) | <0.001 | 0.018 (-0.102, 0.137) | 0.770 | 0.774 (0.717, 0.832) | <0.001 | 0.013 (-0.056, 0.082) | 0.707 | 0.053 (0.011, 0.095) | 0.013 | -0.013 (-0.054, 0.029) | 0.547 |
| | R | 7151 | 0.190 (0.121, 0.259) | <0.001 | 0.024 (-0.096, 0.145) | 0.691 | 0.806 (0.748, 0.865) | <0.001 | -0.054 (-0.122, 0.015) | 0.127 | 0.033 (-0.009, 0.076) | 0.122 | -0.006 (-0.049, 0.037) | 0.783 |
| **Putamen** | L | 7142 | 0.358 (0.274, 0.442) | <0.001 | 0.001 (-0.122, 0.125) | 0.983 | 0.709 (0.644, 0.773) | <0.001 | -0.025 (-0.108, 0.058) | 0.551 | 0.032 (-0.009, 0.074) | 0.129 | -0.002 (-0.044, 0.039) | 0.917 |
| | R | 7149 | 0.354 (0.274, 0.434) | <0.001 | 0.023 (-0.099, 0.145) | 0.713 | 0.704 (0.639, 0.768) | <0.001 | 0.029 (-0.051, 0.110) | 0.476 | 0.033 (-0.009, 0.075) | 0.120 | -0.004 (-0.045, 0.038) | 0.869 |
| **Pallidum** | L | 7085 | 0.348 (0.255, 0.442) | <0.001 | -0.013 (-0.137, 0.110) | 0.834 | 0.703 (0.635, 0.771) | <0.001 | -0.041 (-0.130, 0.047) | 0.361 | 0.002 (-0.039, 0.043) | 0.923 | 0.028 (-0.015, 0.071) | 0.196 |
| | R | 7085 | 0.383 (0.288, 0.477) | <0.001 | -0.025 (-0.149, 0.100) | 0.699 | 0.689 (0.621, 0.757) | <0.001 | -0.021 (-0.110, 0.069) | 0.648 | -0.016 (-0.058, 0.025) | 0.437 | 0.042 (-0.0004, 0.085) | 0.053 |
| **Hippocampus** | L | 7137 | 0.181 (0.106, 0.255) | <0.001 | -0.001 (-0.123, 0.121) | 0.985 | 0.809 (0.750, 0.868) | <0.001 | 0.004 (-0.072, 0.080) | 0.915 | -0.015 (-0.059, 0.028) | 0.493 | 0.025 (-0.018, 0.068) | 0.260 |
| | R | 7150 | 0.205 (0.127, 0.283) | <0.001 | 0.013 (-0.107, 0.133) | 0.832 | 0.798 (0.737, 0.858) | <0.001 | -0.015 (-0.092, 0.062) | 0.697 | -0.023 (-0.066, 0.019) | 0.280 | -0.00006 (-0.042, 0.042) | 0.998 |
| **Amygdala** | L | 7126 | 0.053 (-0.019, 0.124) | 0.148 | 0.019 (-0.100, 0.139) | 0.753 | 0.853 (0.794, 0.912) | <0.001 | 0.027 (-0.045, 0.099) | 0.461 | -0.040 (-0.084, 0.004) | 0.077 | 0.039 (-0.005, 0.084) | 0.085 |
| | R | 7130 | 0.029 (-0.045, 0.103) | 0.442 | 0.040 (-0.079, 0.159) | 0.511 | 0.859 (0.799, 0.918) | <0.001 | 0.029 (-0.044, 0.102) | 0.439 | -0.045 (-0.088, -0.002) | 0.040 | 0.010 (-0.033, 0.053) | 0.649 |
| **Accumbens** | L | 7111 | 0.450 (0.370, 0.531) | <0.001 | 0.017 (-0.106, 0.141) | 0.786 | 0.709 (0.649, 0.769) | <0.001 | 0.020 (-0.056, 0.097) | 0.602 | 0.047 (0.005, 0.090) | 0.030 | -0.003 (-0.046, 0.040) | 0.891 |
| | R | 7106 | 0.374 (0.294, 0.453) | <0.001 | 0.032 (-0.090, 0.155) | 0.605 | 0.730 (0.670, 0.790) | <0.001 | 0.060 (-0.017, 0.138) | 0.127 | 0.045 (0.003, 0.087) | 0.034 | -0.003 (-0.045, 0.039) | 0.899 |
| **Ventral DC** | L | 7087 | 0.176 (0.092, 0.260) | <0.001 | 0.008 (-0.114, 0.130) | 0.896 | 0.817 (0.758, 0.876) | <0.001 | 0.002 (-0.078, 0.083) | 0.956 | -0.037 (-0.080, 0.006) | 0.095 | 0.023 (-0.022, 0.068) | 0.314 |
| | R | 7101 | 0.171 (0.087, 0.256) | <0.001 | -0.005 (-0.127, 0.118) | 0.939 | 0.821 (0.762, 0.880) | <0.001 | 0.086 (0.005, 0.167) | 0.037 | -0.043 (-0.087, -0.00005) | 0.050 | 0.049 (0.004, 0.094) | 0.031 |

**Note:** Estimated regression coefficients (β), 95% confidence intervals (CI) and *p*-values from the linear mixed-effects models evaluating the moderating effects of O3 in the relationship between brain microstructure and Body Mass Index. Models were controlled for age, sex, education, race, ethnicity, puberty, co-exposure to other air pollutants, head motion during the MRI acquisition and urbanicity. Random intercepts were modelled from MRI scanner device and subject ID. Although all two-way interactions were assessed, this table presents only the ones of primary interest due to space constraints. Abbreviations: ROI: region-of-interest; Ventral DC: ventral diencephalon; RSI: restricted spectrum imaging; RNI: restricted normalized isotropic diffusion; H: hemisphere. *p-values < 0.05 of two-way and three-way interactions that achieved the Benjamini-Hochberg False Discovery Rate threshold correction.

**Supplementary Table 5.** Main effects and interactions for the Oxwt measure across both sexes in RNI.

| | | BMI ~ $ROI_{RNI}$ * Oxwt * age + sex + caregiver highest education + race + ethnicity + puberty + PM2.5 + head motion + urbanicity address 1 + (1\|MRI device) + (1\| Subject ID) | | | | | | | | | | | | |
|---|---|---|---|---|---|---|---|---|---|---|---|---|---|---|
| | | | $ROI_{RNI}$ | | Oxwt | | Age | | $ROI_{RNI}$ X Oxwt | | $ROI_{RNI}$ x Age | | $ROI_{RNI}$ x Oxwt x Age | |
| **ROI** | **H** | **n** | **β (95%CI)** | **p-value** | **β (95%CI)** | **p-value** | **β (95%CI)** | **p-value** | **β (95%CI)** | **p-value** | **β (95%CI)** | **p-value** | **β (95%CI)** | **p-value** |
| **Thalamus** | L | 7073 | 0.184 (0.097, 0.272) | <0.001 | 0.032 (-0.095, 0.158) | 0.626 | 0.805 (0.745, 0.866) | <0.001 | 0.004 (-0.084, 0.092) | 0.933 | -0.021 (-0.064, 0.022) | 0.335 | 0.022 (-0.023, 0.068) | 0.334 |
| | R | 7099 | 0.225 (0.139, 0.312) | <0.001 | 0.030 (-0.099, 0.159) | 0.650 | 0.791 (0.730, 0.852) | <0.001 | -0.012 (-0.095, 0.072) | 0.786 | -0.019 (-0.063, 0.025) | 0.402 | 0.006 (-0.038, 0.051) | 0.784 |
| **Caudate** | L | 7166 | 0.295 (0.226, 0.364) | <0.001 | 0.046 (-0.078, 0.169) | 0.471 | 0.775 (0.717, 0.833) | <0.001 | 0.019 (-0.050, 0.088) | 0.597 | 0.050 (0.008, 0.091) | 0.020 | -0.043 (-0.085, -0.00009) | 0.050 |
| | R | 7151 | 0.189 (0.120, 0.258) | <0.001 | 0.052 (-0.073, 0.177) | 0.417 | 0.808 (0.750, 0.866) | <0.001 | -0.040 (-0.109, 0.030) | 0.265 | 0.032 (-0.011, 0.074) | 0.143 | -0.020 (-0.063, 0.023) | 0.355 |
| **Putamen** | L | 7142 | 0.359 (0.275, 0.443) | <0.001 | 0.042 (-0.087, 0.171) | 0.521 | 0.710 (0.646, 0.775) | <0.001 | 0.034 (-0.050, 0.117) | 0.427 | 0.032 (-0.009, 0.074) | 0.127 | 0.001 (-0.041, 0.044) | 0.954 |
| | R | 7149 | 0.356 (0.276, 0.436) | <0.001 | 0.061 (-0.065, 0.187) | 0.344 | 0.705 (0.641, 0.769) | <0.001 | 0.058 (-0.023, 0.139) | 0.162 | 0.032 (-0.009, 0.074) | 0.130 | -0.008 (-0.050, 0.035) | 0.722 |
| **Pallidum** | L | 7085 | 0.350 (0.257, 0.443) | <0.001 | 0.017 (-0.112, 0.145) | 0.798 | 0.703 (0.636, 0.771) | <0.001 | 0.025 (-0.064, 0.115) | 0.579 | 0.004 (-0.037, 0.046) | 0.833 | 0.036 (-0.007, 0.079) | 0.099 |
| | R | 7085 | 0.379 (0.284, 0.473) | <0.001 | 0.018 (-0.111, 0.148) | 0.781 | 0.693 (0.625, 0.761) | <0.001 | 0.020 (-0.073, 0.112) | 0.678 | -0.012 (-0.054, 0.029) | 0.560 | 0.050 (0.007, 0.093) | 0.022 |
| **Hippocampus** | L | 7137 | 0.183 (0.109, 0.258) | <0.001 | 0.029 (-0.097, 0.156) | 0.648 | 0.809 (0.750, 0.868) | <0.001 | 0.009 (-0.067, 0.084) | 0.825 | -0.016 (-0.059, 0.028) | 0.483 | 0.020 (-0.023, 0.064) | 0.361 |
| | R | 7150 | 0.205 (0.127, 0.283) | <0.001 | 0.035 (-0.089, 0.160) | 0.579 | 0.797 (0.737, 0.857) | <0.001 | -0.023 (-0.102, 0.055) | 0.558 | -0.023 (-0.066, 0.019) | 0.280 | 0.012 (-0.030, 0.054) | 0.573 |
| **Amygdala** | L | 7126 | 0.058 (-0.014, 0.129) | 0.114 | 0.037 (-0.087, 0.162) | 0.556 | 0.855 (0.796, 0.914) | <0.001 | 0.049 (-0.022, 0.119) | 0.177 | -0.040 (-0.085, 0.004) | 0.072 | 0.035 (-0.009, 0.079) | 0.121 |
| | R | 7130 | 0.031 (-0.044, 0.106) | 0.415 | 0.055 (-0.069, 0.179) | 0.387 | 0.858 (0.798, 0.917) | <0.001 | 0.007 (-0.068, 0.081) | 0.856 | -0.044 (-0.087, -0.001) | 0.045 | 0.011 (-0.032, 0.053) | 0.619 |
| **Accumbens** | L | 7111 | 0.452 (0.371, 0.532) | <0.001 | 0.067 (-0.062, 0.196) | 0.308 | 0.710 (0.650, 0.769) | <0.001 | 0.050 (-0.027, 0.127) | 0.207 | 0.046 (0.003, 0.088) | 0.037 | -0.017 (-0.061, 0.027) | 0.441 |
| | R | 7106 | 0.379 (0.299, 0.458) | <0.001 | 0.070 (-0.057, 0.198) | 0.279 | 0.729 (0.669, 0.789) | <0.001 | 0.075 (-0.003, 0.153) | 0.058 | 0.044 (0.002, 0.086) | 0.040 | -0.010 (-0.052, 0.033) | 0.659 |
| **Ventral DC** | L | 7087 | 0.181 (0.097, 0.265) | <0.001 | 0.037 (-0.089, 0.164) | 0.561 | 0.818 (0.759, 0.877) | <0.001 | 0.018 (-0.063, 0.099) | 0.668 | -0.031 (-0.075, 0.012) | 0.161 | 0.037 (-0.009, 0.082) | 0.112 |
| | R | 7101 | 0.176 (0.091, 0.260) | <0.001 | 0.030 (-0.096, 0.156) | 0.642 | 0.824 (0.765, 0.883) | <0.001 | 0.071 (-0.010, 0.153) | 0.087 | -0.035 (-0.079, 0.008) | 0.114 | 0.050 (0.006, 0.095) | 0.025 |

**Note:** Estimated regression coefficients (β), 95% confidence intervals (CI) and *p*-values from the linear mixed-effects models evaluating the moderating effects of Oxwt in the relationship between brain microstructure and Body Mass Index. Models were controlled for age, sex, education, race, ethnicity, puberty, co-exposure to other air pollutants, head motion during the MRI acquisition and urbanicity. Random intercepts were modelled from MRI scanner device and subject ID. Although all two-way interactions were assessed, this table presents only the ones of primary interest due to space constraints. Abbreviations: ROI: region-of-interest; Ventral DC: ventral diencephalon; RSI: restricted spectrum imaging; RNI: restricted normalized isotropic diffusion; H: hemisphere. *p-values < 0.05 of two-way and three-way interactions that achieved the Benjamini-Hochberg False Discovery Rate threshold correction.

**Supplementary Table 6.** Main effects and interactions for the PM2.5 pollutant for males in RNI.

| | | BMI ~ $ROI_{RNI}$ * PM2.5 * age + caregiver highest education + race + ethnicity + puberty + NO2 + O3 + head motion + urbanicity address 1 + (1\|MRI device) + (1\| Subject ID) | | | | | | | | | | | | |
|---|---|---|---|---|---|---|---|---|---|---|---|---|---|---|
| | | | $ROI_{RNI}$ | | PM2.5 | | Age | | $ROI_{RNI}$ X PM2.5 | | $ROI_{RNI}$ x Age | | $ROI_{RNI}$ x PM2.5 x Age | |
| **ROI** | **H** | **n** | **β (95%CI)** | **p-value** | **β (95%CI)** | **p-value** | **β (95%CI)** | **p-value** | **β (95%CI)** | **p-value** | **β (95%CI)** | **p-value** | **β (95%CI)** | **p-value** |
| **Thalamus** | L | 3550 | 0.171 (0.052, 0.290) | 0.005 | 0.156 (-0.038, 0.351) | 0.118 | 0.851 (0.775, 0.927) | <0.001 | 0.105 (-0.005, 0.215) | 0.063 | -0.031 (-0.091, 0.028) | 0.303 | 0.020 (-0.033, 0.073) | 0.459 |
| | R | 3562 | 0.121 (0.004, 0.239) | 0.043 | 0.144 (-0.057, 0.345) | 0.163 | 0.864 (0.788, 0.940) | <0.001 | 0.163 (0.051, 0.275) | 0.004 | -0.049 (-0.110, 0.012) | 0.114 | 0.036 (-0.018, 0.090) | 0.186 |
| **Caudate** | L | 3600 | 0.231 (0.136, 0.326) | <0.001 | 0.199 (0.016, 0.382) | 0.035 | 0.842 (0.770, 0.913) | <0.001 | 0.055 (-0.037, 0.147) | 0.244 | 0.017 (-0.040, 0.075) | 0.555 | -0.022 (-0.078, 0.034) | 0.447 |
| | R | 3598 | 0.127 (0.032, 0.221) | 0.009 | 0.184 (-0.002, 0.371) | 0.054 | 0.870 (0.798, 0.941) | <0.001 | 0.086 (-0.009, 0.180) | 0.075 | -0.043 (-0.101, 0.015) | 0.148 | 0.017 (-0.042, 0.076) | 0.569 |
| **Putamen** | L | 3585 | 0.300 (0.184, 0.416) | <0.001 | 0.188 (-0.005, 0.380) | 0.059 | 0.772 (0.691, 0.853) | <0.001 | 0.097 (-0.005, 0.198) | 0.062 | -0.050 (-0.108, 0.009) | 0.099 | -0.001 (-0.059, 0.057) | 0.978 |
| | R | 3589 | 0.375 (0.263, 0.487) | <0.001 | 0.181 (-0.009, 0.370) | 0.064 | 0.740 (0.660, 0.820) | <0.001 | 0.049 (-0.051, 0.149) | 0.334 | -0.044 (-0.102, 0.015) | 0.144 | 0.015 (-0.044, 0.074) | 0.609 |
| **Pallidum** | L | 3554 | 0.351 (0.223, 0.478) | <0.001 | 0.159 (-0.038, 0.356) | 0.117 | 0.744 (0.658, 0.831) | <0.001 | 0.052 (-0.058, 0.163) | 0.352 | -0.023 (-0.080, 0.033) | 0.417 | -0.002 (-0.052, 0.049) | 0.949 |
| | R | 3549 | 0.407 (0.276, 0.538) | <0.001 | 0.181 (-0.020, 0.382) | 0.081 | 0.722 (0.635, 0.809) | <0.001 | 0.101 (-0.014, 0.215) | 0.085 | -0.036 (-0.094, 0.022) | 0.229 | -0.012 (-0.063, 0.039) | 0.651 |
| **Hippocampus** | L | 3595 | 0.073 (-0.029, 0.175) | 0.161 | 0.201 (0.013, 0.389) | 0.038 | 0.888 (0.815, 0.960) | <0.001 | 0.028 (-0.069, 0.124) | 0.571 | -0.050 (-0.110, 0.010) | 0.101 | 0.003 (-0.056, 0.062) | 0.924 |
| | R | 3593 | 0.123 (0.014, 0.232) | 0.027 | 0.220 (0.035, 0.405) | 0.021 | 0.862 (0.787, 0.937) | <0.001 | 0.020 (-0.082, 0.121) | 0.704 | -0.048 (-0.106, 0.011) | 0.111 | 0.024 (-0.029, 0.076) | 0.377 |
| **Amygdala** | L | 3584 | 0.050 (-0.047, 0.147) | 0.314 | 0.183 (-0.003, 0.370) | 0.056 | 0.898 (0.825, 0.970) | <0.001 | 0.045 (-0.050, 0.140) | 0.351 | -0.076 (-0.137, -0.014) | 0.016 | 0.024 (-0.036, 0.083) | 0.436 |
| | R | 3587 | 0.038 (-0.065, 0.140) | 0.473 | 0.227 (0.048, 0.406) | 0.014 | 0.894 (0.821, 0.967) | <0.001 | -0.042 (-0.140, 0.056) | 0.400 | -0.088 (-0.147, -0.028) | 0.004 | 0.016 (-0.036, 0.069) | 0.545 |
| **Accumbens** | L | 3563 | 0.423 (0.312, 0.533) | <0.001 | 0.142 (-0.060, 0.345) | 0.171 | 0.764 (0.689, 0.839) | <0.001 | 0.043 (-0.056, 0.142) | 0.394 | -0.009 (-0.068, 0.050) | 0.765 | -0.024 (-0.082, 0.034) | 0.423 |
| | R | 3568 | 0.343 (0.233, 0.453) | <0.001 | 0.167 (-0.030, 0.364) | 0.098 | 0.784 (0.709, 0.859) | <0.001 | 0.035 (-0.063, 0.134) | 0.480 | -0.020 (-0.077, 0.037) | 0.494 | -0.028 (-0.084, 0.028) | 0.323 |
| **Ventral DC** | L | 3555 | 0.215 (0.104, 0.325) | <0.001 | 0.218 (0.031, 0.405) | 0.024 | 0.847 (0.773, 0.920) | <0.001 | -0.027 (-0.123, 0.069) | 0.580 | -0.029 (-0.089, 0.031) | 0.347 | 0.018 (-0.030, 0.067) | 0.465 |
| | R | 3562 | 0.173 (0.061, 0.285) | 0.003 | 0.209 (0.019, 0.399) | 0.033 | 0.860 (0.787, 0.933) | <0.001 | -0.002 (-0.102, 0.099) | 0.976 | -0.037 (-0.098, 0.023) | 0.230 | 0.003 (-0.047, 0.052) | 0.915 |

**Note:** Estimated regression coefficients (β), 95% confidence intervals (CI) and *p*-values from the linear mixed-effects models evaluating the moderating effects of PM2.5 in the relationship between brain microstructure and Body Mass Index. Models were controlled for age, education, race, ethnicity, puberty, co-exposure to other air pollutants, head motion during the MRI acquisition and urbanicity. Random intercepts were modelled from MRI scanner device and subject ID. Although all two-way interactions were assessed, this table presents only the ones of primary interest due to space constraints. Abbreviations: ROI: region-of-interest; Ventral DC: ventral diencephalon; RSI: restricted spectrum imaging; RNI: restricted normalized isotropic diffusion; H: hemisphere. *p-values < 0.05 of two-way and three-way interactions that achieved the Benjamini-Hochberg False Discovery Rate threshold correction.

**Supplementary Table 7.** Main effects and interactions for the NO2 pollutant for males in RNI.

| | | BMI ~ $ROI_{RNI}$ * NO2 * age + caregiver highest education + race + ethnicity + puberty + PM2.5 + O3 + head motion + urbanicity address 1 + (1\|MRI device) + (1\| Subject ID) | | | | | | | | | | | | |
|---|---|---|---|---|---|---|---|---|---|---|---|---|---|---|
| | | | $ROI_{RNI}$ | | NO2 | | Age | | $ROI_{RNI}$ X NO2 | | $ROI_{RNI}$ x Age | | $ROI_{RNI}$ x NO2 x Age | |
| **ROI** | **H** | **n** | **β (95%CI)** | **p-value** | **β (95%CI)** | **p-value** | **β (95%CI)** | **p-value** | **β (95%CI)** | **p-value** | **β (95%CI)** | **p-value** | **β (95%CI)** | **p-value** |
| **Thalamus** | L | 3550 | 0.170 (0.051, 0.289) | 0.005 | 0.009 (-0.183, 0.201) | 0.930 | 0.845 (0.768, 0.921) | <0.001 | 0.023 (-0.098, 0.144) | 0.711 | -0.039 (-0.099, 0.022) | 0.211 | 0.002 (-0.056, 0.060) | 0.950 |
| | R | 3562 | 0.139 (0.023, 0.255) | 0.020 | 0.022 (-0.174, 0.219) | 0.826 | 0.856 (0.780, 0.932) | <0.001 | 0.099 (-0.022, 0.219) | 0.109 | -0.054 (-0.116, 0.008) | 0.087 | 0.026 (-0.032, 0.085) | 0.376 |
| **Caudate** | L | 3600 | 0.231 (0.136, 0.327) | <0.001 | 0.015 (-0.174, 0.203) | 0.880 | 0.831 (0.759, 0.902) | <0.001 | 0.015 (-0.077, 0.107) | 0.755 | 0.019 (-0.039, 0.076) | 0.526 | -0.060 (-0.116, -0.005) | 0.032 |
| | R | 3598 | 0.137 (0.042, 0.232) | 0.005 | 0.007 (-0.183, 0.198) | 0.940 | 0.858 (0.786, 0.929) | <0.001 | 0.048 (-0.047, 0.143) | 0.321 | -0.045 (-0.103, 0.014) | 0.134 | -0.008 (-0.063, 0.047) | 0.774 |
| **Putamen** | L | 3585 | 0.325 (0.208, 0.442) | <0.001 | 0.027 (-0.168, 0.223) | 0.785 | 0.761 (0.681, 0.842) | <0.001 | 0.115 (0.0003, 0.229) | 0.049 | -0.049 (-0.108, 0.011) | 0.108 | -0.005 (-0.061, 0.052) | 0.875 |
| | R | 3589 | 0.383 (0.271, 0.494) | <0.001 | 0.036 (-0.158, 0.230) | 0.720 | 0.734 (0.654, 0.814) | <0.001 | 0.079 (-0.033, 0.192) | 0.167 | -0.045 (-0.104, 0.014) | 0.135 | -0.005 (-0.061, 0.050) | 0.846 |
| **Pallidum** | L | 3554 | 0.367 (0.239, 0.495) | <0.001 | 0.031 (-0.170, 0.232) | 0.763 | 0.734 (0.647, 0.820) | <0.001 | 0.107 (-0.022, 0.236) | 0.103 | -0.025 (-0.082, 0.032) | 0.393 | 0.008 (-0.048, 0.063) | 0.792 |
| | R | 3549 | 0.409 (0.278, 0.541) | <0.001 | 0.062 (-0.140, 0.264) | 0.550 | 0.716 (0.629, 0.802) | <0.001 | 0.057 (-0.070, 0.185) | 0.378 | -0.042 (-0.100, 0.017) | 0.162 | 0.004 (-0.051, 0.058) | 0.893 |
| **Hippocampus** | L | 3595 | 0.088 (-0.014, 0.190) | 0.092 | 0.008 (-0.185, 0.202) | 0.935 | 0.880 (0.808, 0.953) | <0.001 | 0.051 (-0.058, 0.160) | 0.359 | -0.054 (-0.114, 0.006) | 0.080 | 0.006 (-0.054, 0.066) | 0.845 |
| | R | 3593 | 0.130 (0.022, 0.239) | 0.019 | -0.002 (-0.190, 0.187) | 0.985 | 0.853 (0.779, 0.928) | <0.001 | -0.009 (-0.123, 0.105) | 0.875 | -0.055 (-0.113, 0.004) | 0.068 | 0.009 (-0.048, 0.066) | 0.764 |
| **Amygdala** | L | 3584 | 0.066 (-0.032, 0.163) | 0.187 | -0.005 (-0.196, 0.185) | 0.956 | 0.896 (0.824, 0.969) | <0.001 | 0.141 (0.041, 0.241) | 0.006 | -0.076 (-0.138, -0.015) | 0.015 | 0.035 (-0.028, 0.098) | 0.282 |
| | R | 3587 | 0.033 (-0.069, 0.136) | 0.526 | -0.024 (-0.209, 0.161) | 0.802 | 0.889 (0.816, 0.962) | <0.001 | -0.002 (-0.113, 0.109) | 0.973 | -0.086 (-0.146, -0.025) | 0.006 | 0.030 (-0.029, 0.088) | 0.320 |
| **Accumbens** | L | 3563 | 0.439 (0.328, 0.550) | <0.001 | 0.079 (-0.128, 0.286) | 0.457 | 0.749 (0.675, 0.824) | <0.001 | 0.056 (-0.054, 0.166) | 0.320 | -0.013 (-0.072, 0.047) | 0.671 | -0.043 (-0.103, 0.016) | 0.151 |
| | R | 3568 | 0.356 (0.245, 0.467) | <0.001 | 0.043 (-0.159, 0.246) | 0.674 | 0.772 (0.698, 0.847) | <0.001 | 0.034 (-0.073, 0.141) | 0.530 | -0.019 (-0.077, 0.038) | 0.512 | -0.010 (-0.067, 0.046) | 0.721 |
| **Ventral DC** | L | 3555 | 0.217 (0.104, 0.329) | <0.001 | -0.001 (-0.191, 0.189) | 0.995 | 0.842 (0.768, 0.915) | <0.001 | 0.014 (-0.101, 0.130) | 0.807 | -0.026 (-0.089, 0.037) | 0.424 | 0.032 (-0.030, 0.094) | 0.313 |
| | R | 3562 | 0.177 (0.065, 0.290) | 0.002 | 0.009 (-0.182, 0.200) | 0.925 | 0.854 (0.781, 0.927) | <0.001 | 0.003 (-0.113, 0.118) | 0.966 | -0.044 (-0.108, 0.020) | 0.176 | 0.005 (-0.057, 0.066) | 0.884 |

**Note:** Estimated regression coefficients (β), 95% confidence intervals (CI) and *p*-values from the linear mixed-effects models evaluating the moderating effects of NO2 in the relationship between brain microstructure and Body Mass Index. Models were controlled for age, education, race, ethnicity, puberty, co-exposure to other air pollutants, head motion during the MRI acquisition and urbanicity. Random intercepts were modelled from MRI scanner device and subject ID. Although all two-way interactions were assessed, this table presents only the ones of primary interest due to space constraints. Abbreviations: ROI: region-of-interest; Ventral DC: ventral diencephalon; RSI: restricted spectrum imaging; RNI: restricted normalized isotropic diffusion; H: hemisphere. *p-values < 0.05 of two-way and three-way interactions that achieved the Benjamini-Hochberg False Discovery Rate threshold correction.

**Supplementary Table 8.** Main effects and interactions for the O3 pollutant for males in RNI.

| | | BMI ~ $ROI_{RNI}$ * O3 * age + caregiver highest education + race + ethnicity + puberty + PM2.5 + NO2 + head motion + urbanicity address 1 + (1\|MRI device) + (1\| Subject ID) | | | | | | | | | | | | |
|---|---|---|---|---|---|---|---|---|---|---|---|---|---|---|
| | | | $ROI_{RNI}$ | | O3 | | Age | | $ROI_{RNI}$ X O3 | | $ROI_{RNI}$ x Age | | $ROI_{RNI}$ x O3 x Age | |
| ROI | H | n | β (95%CI) | p-value | β (95%CI) | p-value | β (95%CI) | p-value | β (95%CI) | p-value | β (95%CI) | p-value | β (95%CI) | p-value |
| Thalamus | L | 3550 | 0.165 (0.047, 0.283) | 0.006 | 0.027 (-0.137, 0.192) | 0.745 | 0.847 (0.771, 0.923) | <0.001 | 0.007 (-0.112, 0.126) | 0.912 | -0.035 (-0.094, 0.025) | 0.255 | -0.013 (-0.073, 0.047) | 0.666 |
| | R | 3562 | 0.123 (0.007, 0.238) | 0.039 | 0.030 (-0.136, 0.196) | 0.725 | 0.861 (0.785, 0.937) | <0.001 | -0.035 (-0.152, 0.081) | 0.552 | -0.057 (-0.118, 0.004) | 0.067 | -0.035 (-0.098, 0.027) | 0.265 |
| Caudate | L | 3600 | 0.225 (0.130, 0.321) | <0.001 | 0.034 (-0.127, 0.194) | 0.680 | 0.834 (0.762, 0.905) | <0.001 | -0.020 (-0.114, 0.075) | 0.683 | 0.026 (-0.031, 0.083) | 0.367 | -0.016 (-0.073, 0.040) | 0.571 |
| | R | 3598 | 0.128 (0.033, 0.222) | 0.008 | 0.034 (-0.129, 0.197) | 0.681 | 0.861 (0.790, 0.933) | <0.001 | -0.060 (-0.156, 0.036) | 0.221 | -0.044 (-0.102, 0.015) | 0.143 | 0.003 (-0.056, 0.061) | 0.932 |
| Putamen | L | 3585 | 0.307 (0.191, 0.422) | <0.001 | 0.023 (-0.143, 0.189) | 0.784 | 0.765 (0.684, 0.845) | <0.001 | -0.026 (-0.143, 0.090) | 0.659 | -0.046 (-0.105, 0.012) | 0.121 | -0.007 (-0.066, 0.053) | 0.822 |
| | R | 3589 | 0.374 (0.263, 0.484) | <0.001 | 0.058 (-0.106, 0.223) | 0.486 | 0.733 (0.654, 0.813) | <0.001 | 0.054 (-0.059, 0.167) | 0.348 | -0.046 (-0.104, 0.012) | 0.122 | -0.017 (-0.076, 0.042) | 0.576 |
| Pallidum | L | 3554 | 0.346 (0.220, 0.473) | <0.001 | 0.006 (-0.161, 0.173) | 0.943 | 0.740 (0.654, 0.826) | <0.001 | -0.071 (-0.196, 0.053) | 0.262 | -0.026 (-0.083, 0.030) | 0.361 | 0.011 (-0.049, 0.071) | 0.722 |
| | R | 3549 | 0.398 (0.268, 0.528) | <0.001 | -0.009 (-0.177, 0.159) | 0.920 | 0.715 (0.629, 0.802) | <0.001 | 0.011 (-0.119, 0.141) | 0.869 | -0.040 (-0.097, 0.018) | 0.177 | 0.040 (-0.019, 0.100) | 0.183 |
| Hippocampus | L | 3595 | 0.075 (-0.027, 0.177) | 0.151 | 0.023 (-0.141, 0.186) | 0.787 | 0.883 (0.810, 0.955) | <0.001 | -0.014 (-0.120, 0.092) | 0.792 | -0.053 (-0.113, 0.007) | 0.083 | -0.009 (-0.068, 0.051) | 0.775 |
| | R | 3593 | 0.127 (0.019, 0.236) | 0.022 | 0.054 (-0.108, 0.215) | 0.515 | 0.857 (0.783, 0.932) | <0.001 | -0.042 (-0.151, 0.066) | 0.444 | -0.054 (-0.113, 0.004) | 0.069 | -0.050 (-0.107, 0.007) | 0.089 |
| Amygdala | L | 3584 | 0.061 (-0.036, 0.158) | 0.217 | 0.053 (-0.109, 0.215) | 0.523 | 0.894 (0.822, 0.966) | <0.001 | 0.086 (-0.012, 0.184) | 0.087 | -0.084 (-0.145, -0.023) | 0.007 | -0.027 (-0.089, 0.034) | 0.381 |
| | R | 3587 | 0.036 (-0.065, 0.137) | 0.486 | 0.085 (-0.075, 0.245) | 0.298 | 0.892 (0.819, 0.965) | <0.001 | 0.075 (-0.026, 0.177) | 0.145 | -0.094 (-0.153, -0.035) | 0.002 | -0.021 (-0.081, 0.038) | 0.487 |
| Accumbens | L | 3563 | 0.426 (0.316, 0.535) | <0.001 | 0.026 (-0.141, 0.194) | 0.758 | 0.756 (0.681, 0.830) | <0.001 | -0.005 (-0.110, 0.101) | 0.930 | -0.004 (-0.063, 0.054) | 0.891 | 0.009 (-0.050, 0.069) | 0.756 |
| | R | 3568 | 0.348 (0.238, 0.458) | <0.001 | 0.060 (-0.107, 0.226) | 0.481 | 0.777 (0.703, 0.852) | <0.001 | 0.104 (-0.006, 0.215) | 0.064 | -0.018 (-0.075, 0.038) | 0.526 | -0.001 (-0.060, 0.057) | 0.962 |
| Ventral DC | L | 3555 | 0.214 (0.103, 0.325) | <0.001 | 0.048 (-0.116, 0.211) | 0.567 | 0.842 (0.769, 0.915) | <0.001 | 0.061 (-0.048, 0.170) | 0.272 | -0.036 (-0.096, 0.024) | 0.237 | -0.012 (-0.074, 0.051) | 0.714 |
| | R | 3562 | 0.185 (0.074, 0.297) | 0.001 | 0.034 (-0.130, 0.198) | 0.688 | 0.855 (0.783, 0.928) | <0.001 | 0.157 (0.046, 0.269) | 0.006 | -0.046 (-0.105, 0.014) | 0.135 | 0.023 (-0.041, 0.087) | 0.475 |

**Note:** Estimated regression coefficients (β), 95% confidence intervals (CI) and *p*-values from the linear mixed-effects models evaluating the moderating effects of O3 in the relationship between brain microstructure and Body Mass Index. Models were controlled for age, education, race, ethnicity, puberty, co-exposure to other air pollutants, head motion during the MRI acquisition and urbanicity. Random intercepts were modelled from MRI scanner device and subject ID. Although all two-way interactions were assessed, this table presents only the ones of primary interest due to space constraints. Abbreviations: ROI: region-of-interest; Ventral DC: ventral diencephalon; RSI: restricted spectrum imaging; RNI: restricted normalized isotropic diffusion; H: hemisphere. *p-values < 0.05 of two-way and three-way interactions that achieved the Benjamini-Hochberg False Discovery Rate threshold correction.

**Supplementary Table 9.** Main effects and interactions for the Oxwt measure in males in RNI.

| | | BMI ~ $ROI_{RNI}$ * Oxwt * age + caregiver highest education + race + ethnicity + puberty + PM2.5 + head motion + urbanicity address 1 + (1\|MRI device) + (1\| Subject ID) | | | | | | | | | | | | |
|---|---|---|---|---|---|---|---|---|---|---|---|---|---|---|
| | | | $ROI_{RNI}$ | | Oxwt | | Age | | $ROI_{RNI}$ X Oxwt | | $ROI_{RNI}$ x Age | | $ROI_{RNI}$ x Oxwt x Age | |
| **ROI** | **H** | **n** | **β (95%CI)** | **p-value** | **β (95%CI)** | **p-value** | **β (95%CI)** | **p-value** | **β (95%CI)** | **p-value** | **β (95%CI)** | **p-value** | **β (95%CI)** | **p-value** |
| **Thalamus** | L | 3550 | 0.166 (0.048, 0.284) | 0.006 | 0.030 (-0.140, 0.201) | 0.727 | 0.846 (0.771, 0.922) | <0.001 | 0.019 (-0.104, 0.142) | 0.759 | -0.039 (-0.099, 0.021) | 0.199 | -0.014 (-0.076, 0.048) | 0.661 |
| | R | 3562 | 0.126 (0.010, 0.242) | 0.034 | 0.035 (-0.137, 0.207) | 0.692 | 0.858 (0.782, 0.934) | <0.001 | 0.024 (-0.093, 0.142) | 0.686 | -0.060 (-0.122, 0.001) | 0.056 | -0.015 (-0.078, 0.048) | 0.647 |
| **Caudate** | L | 3600 | 0.229 (0.134, 0.325) | <0.001 | 0.038 (-0.129, 0.205) | 0.657 | 0.833 (0.762, 0.905) | <0.001 | -0.012 (-0.107, 0.084) | 0.808 | 0.021 (-0.037, 0.078) | 0.481 | -0.055 (-0.112, 0.002) | 0.061 |
| | R | 3598 | 0.129 (0.034, 0.224) | 0.008 | 0.036 (-0.134, 0.206) | 0.681 | 0.861 (0.789, 0.933) | <0.001 | -0.023 (-0.120, 0.073) | 0.637 | -0.045 (-0.104, 0.013) | 0.129 | -0.003 (-0.060, 0.054) | 0.922 |
| **Putamen** | L | 3585 | 0.306 (0.190, 0.421) | <0.001 | 0.041 (-0.133, 0.214) | 0.647 | 0.767 (0.686, 0.847) | <0.001 | 0.044 (-0.073, 0.162) | 0.460 | -0.047 (-0.106, 0.011) | 0.115 | -0.009 (-0.070, 0.053) | 0.786 |
| | R | 3589 | 0.374 (0.263, 0.485) | <0.001 | 0.074 (-0.098, 0.245) | 0.399 | 0.736 (0.656, 0.816) | <0.001 | 0.089 (-0.024, 0.202) | 0.122 | -0.047 (-0.105, 0.011) | 0.114 | -0.017 (-0.077, 0.043) | 0.575 |
| **Pallidum** | L | 3554 | 0.347 (0.220, 0.474) | <0.001 | 0.017 (-0.157, 0.190) | 0.849 | 0.740 (0.654, 0.826) | <0.001 | -0.004 (-0.133, 0.124) | 0.945 | -0.027 (-0.083, 0.030) | 0.360 | 0.015 (-0.045, 0.076) | 0.620 |
| | R | 3549 | 0.395 (0.266, 0.524) | <0.001 | 0.020 (-0.154, 0.195) | 0.821 | 0.719 (0.633, 0.805) | <0.001 | 0.044 (-0.090, 0.178) | 0.521 | -0.037 (-0.095, 0.021) | 0.213 | 0.037 (-0.023, 0.097) | 0.228 |
| **Hippocampus** | L | 3595 | 0.078 (-0.025, 0.181) | 0.137 | 0.026 (-0.144, 0.196) | 0.767 | 0.883 (0.811, 0.955) | <0.001 | 0.015 (-0.091, 0.121) | 0.787 | -0.054 (-0.114, 0.006) | 0.077 | -0.005 (-0.066, 0.055) | 0.867 |
| | R | 3593 | 0.124 (0.015, 0.233) | 0.025 | 0.048 (-0.121, 0.216) | 0.580 | 0.855 (0.780, 0.929) | <0.001 | -0.041 (-0.154, 0.072) | 0.475 | -0.057 (-0.116, 0.001) | 0.054 | -0.041 (-0.100, 0.018) | 0.170 |
| **Amygdala** | L | 3584 | 0.064 (-0.033, 0.162) | 0.197 | 0.047 (-0.122, 0.216) | 0.587 | 0.898 (0.826, 0.970) | <0.001 | 0.143 (0.047, 0.240) | 0.004 | -0.084 (-0.145, -0.023) | 0.007 | -0.005 (-0.067, 0.056) | 0.863 |
| | R | 3587 | 0.031 (-0.072, 0.134) | 0.554 | 0.064 (-0.105, 0.232) | 0.460 | 0.893 (0.820, 0.966) | <0.001 | 0.064 (-0.038, 0.167) | 0.220 | -0.092 (-0.152, -0.033) | 0.002 | -0.001 (-0.061, 0.058) | 0.963 |
| **Accumbens** | L | 3563 | 0.428 (0.318, 0.538) | <0.001 | 0.060 (-0.115, 0.235) | 0.500 | 0.755 (0.681, 0.830) | <0.001 | 0.026 (-0.083, 0.135) | 0.641 | -0.008 (-0.066, 0.051) | 0.800 | -0.022 (-0.084, 0.041) | 0.495 |
| | R | 3568 | 0.359 (0.248, 0.470) | <0.001 | 0.081 (-0.093, 0.256) | 0.360 | 0.774 (0.699, 0.849) | <0.001 | 0.106 (-0.005, 0.216) | 0.061 | -0.020 (-0.078, 0.037) | 0.485 | -0.010 (-0.070, 0.050) | 0.752 |
| **Ventral DC** | L | 3555 | 0.216 (0.104, 0.329) | <0.001 | 0.046 (-0.124, 0.217) | 0.595 | 0.842 (0.769, 0.916) | <0.001 | 0.065 (-0.048, 0.178) | 0.261 | -0.035 (-0.097, 0.026) | 0.258 | 0.008 (-0.057, 0.073) | 0.815 |
| | R | 3562 | 0.190 (0.077, 0.303) | 0.001 | 0.044 (-0.126, 0.215) | 0.611 | 0.858 (0.786, 0.931) | <0.001 | 0.136 (0.024, 0.249) | 0.018 | -0.041 (-0.102, 0.020) | 0.188 | 0.018 (-0.046, 0.082) | 0.574 |

**Note:** Estimated regression coefficients (β), 95% confidence intervals (CI) and *p*-values from the linear mixed-effects models evaluating the moderating effects of Oxwt in the relationship between brain microstructure and Body Mass Index. Models were controlled for age, education, race, ethnicity, puberty, co-exposure to other air pollutants, head motion during the MRI acquisition and urbanicity. Random intercepts were modelled from MRI scanner device and subject ID. Although all two-way interactions were assessed, this table presents only the ones of primary interest due to space constraints. Abbreviations: ROI: region-of-interest; Ventral DC: ventral diencephalon; RSI: restricted spectrum imaging; RNI: restricted normalized isotropic diffusion; H: hemisphere. *p-values < 0.05 of two-way and three-way interactions that achieved the Benjamini-Hochberg False Discovery Rate threshold correction.

**Supplementary Table 10.** Main effects and interactions for the PM2.5 pollutant for females in RNI.

| | | BMI ~ $ROI_{RNI}$ * PM2.5 * age + caregiver highest education + race + ethnicity + puberty + NO2 + O3 + head motion + urbanicity address 1 + (1\|MRI device) + (1\| Subject ID) | | | | | | | | | | | | |
|---|---|---|---|---|---|---|---|---|---|---|---|---|---|---|
| | | | $ROI_{RNI}$ | | PM2.5 | | Age | | $ROI_{RNI}$ X PM2.5 | | $ROI_{RNI}$ x Age | | $ROI_{RNI}$ x PM2.5 x Age | |
| **ROI** | **H** | **n** | **β (95%CI)** | **p-value** | **β (95%CI)** | **p-value** | **β (95%CI)** | **p-value** | **β (95%CI)** | **p-value** | **β (95%CI)** | **p-value** | **β (95%CI)** | **p-value** |
| **Thalamus** | L | 3523 | 0.164 (0.044, 0.285) | 0.007 | 0.137 (-0.038, 0.312) | 0.128 | 0.722 (0.621, 0.824) | <0.001 | -0.131 (-0.243, -0.020) | 0.021 | 0.007 (-0.057, 0.070) | 0.839 | -0.006 (-0.063, 0.050) | 0.823 |
| | R | 3537 | 0.260 (0.141, 0.380) | <0.001 | 0.116 (-0.066, 0.299) | 0.214 | 0.694 (0.592, 0.795) | <0.001 | -0.097 (-0.206, 0.012) | 0.081 | 0.025 (-0.039, 0.089) | 0.445 | 0.022 (-0.036, 0.079) | 0.463 |
| **Caudate** | L | 3566 | 0.353 (0.253, 0.453) | <0.001 | 0.079 (-0.096, 0.254) | 0.376 | 0.661 (0.563, 0.759) | <0.001 | 0.038 (-0.060, 0.135) | 0.450 | 0.081 (0.019, 0.143) | 0.010 | -0.040 (-0.102, 0.022) | 0.208 |
| | R | 3553 | 0.240 (0.141, 0.339) | <0.001 | 0.079 (-0.097, 0.255) | 0.382 | 0.694 (0.596, 0.792) | <0.001 | -0.068 (-0.161, 0.024) | 0.147 | 0.118 (0.055, 0.181) | <0.001 | -0.002 (-0.061, 0.057) | 0.940 |
| **Putamen** | L | 3557 | 0.353 (0.235, 0.472) | <0.001 | 0.102 (-0.083, 0.287) | 0.283 | 0.598 (0.490, 0.706) | <0.001 | -0.084 (-0.190, 0.023) | 0.123 | 0.126 (0.065, 0.186) | <0.001 | -0.041 (-0.098, 0.015) | 0.154 |
| | R | 3560 | 0.311 (0.197, 0.425) | <0.001 | 0.105 (-0.072, 0.281) | 0.247 | 0.603 (0.495, 0.712) | <0.001 | -0.089 (-0.191, 0.014) | 0.090 | 0.123 (0.061, 0.185) | <0.001 | -0.031 (-0.092, 0.029) | 0.307 |
| **Pallidum** | L | 3531 | 0.280 (0.152, 0.407) | <0.001 | 0.111 (-0.067, 0.288) | 0.225 | 0.637 (0.527, 0.746) | <0.001 | -0.049 (-0.157, 0.058) | 0.367 | 0.053 (-0.009, 0.114) | 0.094 | -0.020 (-0.074, 0.034) | 0.468 |
| | R | 3536 | 0.275 (0.148, 0.402) | <0.001 | 0.104 (-0.078, 0.287) | 0.265 | 0.647 (0.538, 0.756) | <0.001 | -0.016 (-0.127, 0.095) | 0.776 | 0.033 (-0.028, 0.093) | 0.293 | -0.011 (-0.063, 0.041) | 0.683 |
| **Hippocampus** | L | 3542 | 0.277 (0.170, 0.385) | <0.001 | 0.116 (-0.063, 0.294) | 0.207 | 0.676 (0.575, 0.777) | <0.001 | -0.092 (-0.195, 0.011) | 0.081 | 0.032 (-0.032, 0.097) | 0.326 | -0.033 (-0.098, 0.032) | 0.323 |
| | R | 3557 | 0.257 (0.147, 0.366) | <0.001 | 0.149 (-0.027, 0.325) | 0.099 | 0.697 (0.596, 0.799) | <0.001 | -0.062 (-0.167, 0.044) | 0.252 | 0.013 (-0.049, 0.075) | 0.681 | -0.013 (-0.076, 0.049) | 0.679 |
| **Amygdala** | L | 3542 | 0.052 (-0.050, 0.154) | 0.316 | 0.170 (0.001, 0.340) | 0.050 | 0.762 (0.663, 0.862) | <0.001 | -0.096 (-0.191, -0.001) | 0.047 | 0.010 (-0.054, 0.074) | 0.759 | -0.046 (-0.106, 0.013) | 0.125 |
| | R | 3543 | 0.032 (-0.072, 0.136) | 0.547 | 0.175 (0.005, 0.345) | 0.046 | 0.775 (0.675, 0.875) | <0.001 | -0.113 (-0.207, -0.019) | 0.018 | 0.011 (-0.053, 0.075) | 0.736 | -0.016 (-0.072, 0.041) | 0.584 |
| **Accumbens** | L | 3548 | 0.438 (0.324, 0.553) | <0.001 | 0.037 (-0.148, 0.221) | 0.698 | 0.622 (0.521, 0.722) | <0.001 | -0.108 (-0.209, -0.008) | 0.034 | 0.122 (0.058, 0.185) | <0.001 | -0.051 (-0.113, 0.011) | 0.108 |
| | R | 3538 | 0.368 (0.258, 0.479) | <0.001 | 0.078 (-0.102, 0.257) | 0.398 | 0.638 (0.537, 0.738) | <0.001 | -0.092 (-0.192, 0.008) | 0.071 | 0.129 (0.068, 0.191) | <0.001 | -0.044 (-0.103, 0.015) | 0.146 |
| **Ventral DC** | L | 3532 | 0.113 (-0.004, 0.229) | 0.059 | 0.144 (-0.032, 0.320) | 0.111 | 0.751 (0.652, 0.850) | <0.001 | -0.107 (-0.209, -0.004) | 0.043 | -0.025 (-0.088, 0.039) | 0.444 | -0.021 (-0.073, 0.031) | 0.436 |
| | R | 3539 | 0.132 (0.016, 0.249) | 0.027 | 0.154 (-0.021, 0.329) | 0.087 | 0.749 (0.650, 0.848) | <0.001 | -0.098 (-0.201, 0.005) | 0.063 | -0.032 (-0.095, 0.032) | 0.327 | 0.006 (-0.047, 0.060) | 0.813 |

**Note:** Estimated regression coefficients (β), 95% confidence intervals (CI) and *p*-values from the linear mixed-effects models evaluating the moderating effects of PM2.5 in the relationship between brain microstructure and Body Mass Index. Models were controlled for age, education, race, ethnicity, puberty, co-exposure to other air pollutants, head motion during the MRI acquisition and urbanicity. Random intercepts were modelled from MRI scanner device and subject ID. Although all two-way interactions were assessed, this table presents only the ones of primary interest due to space constraints. Abbreviations: ROI: region-of-interest; Ventral DC: ventral diencephalon; RSI: restricted spectrum imaging; RNI: restricted normalized isotropic diffusion; H: hemisphere. *p-values < 0.05 of two-way and three-way interactions that achieved the Benjamini-Hochberg False Discovery Rate threshold correction.

**Supplementary Table 11.** Main effects and interactions for the NO2 pollutant for females in RNI.

| | | BMI ~ ROI$_{RNI}$ * NO2 * age + caregiver highest education + race + ethnicity + puberty + PM2.5 + O3 + head motion + urbanicity address 1 + (1\|MRI device) + (1\| Subject ID) | | | | | | | | | | | | |
|---|---|---|---|---|---|---|---|---|---|---|---|---|---|---|
| | | | ROI$_{RNI}$ | | NO2 | | Age | | ROI$_{RNI}$ X NO2 | | ROI$_{RNI}$ x Age | | ROI$_{RNI}$ x NO2 x Age | |
| **ROI** | **H** | **n** | **β (95%CI)** | **p-value** | **β (95%CI)** | **p-value** | **β (95%CI)** | **p-value** | **β (95%CI)** | **p-value** | **β (95%CI)** | **p-value** | **β (95%CI)** | **p-value** |
| **Thalamus** | L | 3523 | 0.161 (0.040, 0.282) | 0.009 | 0.003 (-0.177, 0.183) | 0.971 | 0.731 (0.630, 0.833) | <0.001 | 0.016 (-0.110, 0.142) | 0.803 | 0.014 (-0.050, 0.078) | 0.668 | 0.027 (-0.036, 0.091) | 0.400 |
| | R | 3537 | 0.256 (0.138, 0.375) | <0.001 | 0.030 (-0.158, 0.217) | 0.756 | 0.699 (0.597, 0.800) | <0.001 | 0.008 (-0.113, 0.128) | 0.902 | 0.032 (-0.032, 0.095) | 0.326 | 0.009 (-0.055, 0.072) | 0.790 |
| **Caudate** | L | 3566 | 0.352 (0.253, 0.451) | <0.001 | 0.011 (-0.170, 0.191) | 0.906 | 0.661 (0.563, 0.758) | <0.001 | 0.005 (-0.099, 0.109) | 0.925 | 0.086 (0.025, 0.147) | 0.006 | -0.056 (-0.119, 0.008) | 0.085 |
| | R | 3553 | 0.232 (0.134, 0.329) | <0.001 | 0.031 (-0.149, 0.212) | 0.734 | 0.706 (0.609, 0.804) | <0.001 | -0.058 (-0.161, 0.045) | 0.271 | 0.124 (0.062, 0.186) | <0.001 | -0.067 (-0.129, -0.006) | 0.033 |
| **Putamen** | L | 3557 | 0.358 (0.240, 0.476) | <0.001 | 0.028 (-0.162, 0.219) | 0.772 | 0.602 (0.495, 0.710) | <0.001 | 0.099 (-0.025, 0.223) | 0.117 | 0.120 (0.060, 0.180) | <0.001 | -0.015 (-0.074, 0.044) | 0.619 |
| | R | 3560 | 0.313 (0.199, 0.426) | <0.001 | 0.016 (-0.166, 0.199) | 0.860 | 0.613 (0.505, 0.720) | <0.001 | 0.035 (-0.085, 0.155) | 0.569 | 0.125 (0.063, 0.186) | <0.001 | -0.036 (-0.096, 0.023) | 0.234 |
| **Pallidum** | L | 3531 | 0.293 (0.166, 0.420) | <0.001 | -0.018 (-0.201, 0.166) | 0.852 | 0.642 (0.532, 0.751) | <0.001 | 0.116 (-0.014, 0.247) | 0.080 | 0.052 (-0.009, 0.112) | 0.096 | 0.022 (-0.041, 0.085) | 0.489 |
| | R | 3536 | 0.292 (0.165, 0.419) | <0.001 | -0.001 (-0.186, 0.185) | 0.995 | 0.650 (0.541, 0.759) | <0.001 | 0.082 (-0.049, 0.212) | 0.221 | 0.037 (-0.024, 0.097) | 0.239 | 0.024 (-0.036, 0.084) | 0.427 |
| **Hippocampus** | L | 3542 | 0.268 (0.163, 0.373) | <0.001 | 0.028 (-0.158, 0.214) | 0.768 | 0.683 (0.582, 0.784) | <0.001 | -0.047 (-0.161, 0.067) | 0.420 | 0.032 (-0.032, 0.096) | 0.329 | -0.027 (-0.094, 0.041) | 0.442 |
| | R | 3557 | 0.258 (0.150, 0.366) | <0.001 | 0.019 (-0.164, 0.202) | 0.838 | 0.705 (0.604, 0.806) | <0.001 | -0.044 (-0.160, 0.071) | 0.455 | 0.015 (-0.047, 0.076) | 0.641 | 0.025 (-0.038, 0.088) | 0.441 |
| **Amygdala** | L | 3542 | 0.048 (-0.053, 0.148) | 0.352 | 0.004 (-0.171, 0.178) | 0.965 | 0.775 (0.675, 0.874) | <0.001 | -0.028 (-0.133, 0.077) | 0.596 | 0.004 (-0.060, 0.068) | 0.907 | -0.045 (-0.112, 0.023) | 0.194 |
| | R | 3543 | 0.024 (-0.078, 0.126) | 0.647 | 0.013 (-0.162, 0.188) | 0.883 | 0.782 (0.682, 0.882) | <0.001 | -0.060 (-0.173, 0.052) | 0.292 | 0.001 (-0.061, 0.064) | 0.966 | -0.035 (-0.099, 0.029) | 0.282 |
| **Accumbens** | L | 3548 | 0.419 (0.307, 0.531) | <0.001 | 0.043 (-0.147, 0.232) | 0.661 | 0.627 (0.527, 0.727) | <0.001 | 0.051 (-0.064, 0.166) | 0.382 | 0.118 (0.055, 0.181) | <0.001 | -0.016 (-0.079, 0.047) | 0.622 |
| | R | 3538 | 0.357 (0.247, 0.466) | <0.001 | 0.021 (-0.163, 0.205) | 0.822 | 0.642 (0.542, 0.742) | <0.001 | 0.028 (-0.087, 0.144) | 0.629 | 0.124 (0.062, 0.185) | <0.001 | -0.025 (-0.085, 0.035) | 0.413 |
| **Ventral DC** | L | 3532 | 0.112 (-0.004, 0.228) | 0.060 | 0.010 (-0.171, 0.190) | 0.916 | 0.763 (0.664, 0.862) | <0.001 | 0.048 (-0.071, 0.168) | 0.429 | -0.024 (-0.088, 0.041) | 0.473 | 0.011 (-0.055, 0.077) | 0.741 |
| | R | 3539 | 0.124 (0.009, 0.238) | 0.035 | -0.009 (-0.187, 0.170) | 0.923 | 0.758 (0.659, 0.857) | <0.001 | -0.003 (-0.123, 0.117) | 0.962 | -0.023 (-0.087, 0.041) | 0.476 | 0.023 (-0.041, 0.087) | 0.480 |

**Note:** Estimated regression coefficients (β), 95% confidence intervals (CI) and *p*-values from the linear mixed-effects models evaluating the moderating effects of NO2 in the relationship between brain microstructure and Body Mass Index. Models were controlled for age, education, race, ethnicity, puberty, co-exposure to other air pollutants, head motion during the MRI acquisition and urbanicity. Random intercepts were modelled from MRI scanner device and subject ID. Although all two-way interactions were assessed, this table presents only the ones of primary interest due to space constraints. Abbreviations: ROI: region-of-interest; Ventral DC: ventral diencephalon; RSI: restricted spectrum imaging; RNI: restricted normalized isotropic diffusion; H: hemisphere. *p-values < 0.05 of two-way and three-way interactions that achieved the Benjamini-Hochberg False Discovery Rate threshold correction.

**Supplementary Table 12.** Main effects and interactions for the O3 pollutant for females in RNI.

| | | BMI ~ $ROI_{RNI}$ * O3 * age + caregiver highest education + race + ethnicity + puberty + PM2.5 + NO2 + head motion + urbanicity address 1 + (1\|MRI device) + (1\| Subject ID) | | | | | | | | | | | | |
|---|---|---|---|---|---|---|---|---|---|---|---|---|---|---|
| | | | $ROI_{RNI}$ | | O3 | | Age | | $ROI_{RNI}$ X O3 | | $ROI_{RNI}$ x Age | | $ROI_{RNI}$ x O3 x Age | |
| **ROI** | **H** | **n** | **β (95%CI)** | **p-value** | **β (95%CI)** | **p-value** | **β (95%CI)** | **p-value** | **β (95%CI)** | **p-value** | **β (95%CI)** | **p-value** | **β (95%CI)** | **p-value** |
| **Thalamus** | L | 3523 | 0.157 (0.036, 0.277) | 0.011 | 0.042 (-0.118, 0.201) | 0.611 | 0.725 (0.623, 0.827) | <0.001 | -0.024 (-0.144, 0.096) | 0.694 | 0.002 (-0.062, 0.065) | 0.956 | 0.037 (-0.028, 0.102) | 0.268 |
| | R | 3537 | 0.250 (0.131, 0.368) | <0.001 | 0.016 (-0.147, 0.180) | 0.847 | 0.695 (0.593, 0.796) | <0.001 | -0.067 (-0.179, 0.046) | 0.247 | 0.028 (-0.035, 0.092) | 0.387 | 0.008 (-0.056, 0.072) | 0.815 |
| **Caudate** | L | 3566 | 0.350 (0.251, 0.449) | <0.001 | 0.051 (-0.108, 0.209) | 0.531 | 0.659 (0.561, 0.756) | <0.001 | 0.040 (-0.060, 0.141) | 0.433 | 0.089 (0.028, 0.150) | 0.005 | -0.015 (-0.077, 0.047) | 0.633 |
| | R | 3553 | 0.234 (0.136, 0.332) | <0.001 | 0.065 (-0.094, 0.225) | 0.424 | 0.697 (0.599, 0.795) | <0.001 | -0.043 (-0.141, 0.055) | 0.392 | 0.123 (0.061, 0.185) | <0.001 | -0.028 (-0.092, 0.035) | 0.381 |
| **Putamen** | L | 3557 | 0.342 (0.225, 0.459) | <0.001 | 0.040 (-0.125, 0.204) | 0.636 | 0.598 (0.490, 0.705) | <0.001 | -0.043 (-0.161, 0.075) | 0.474 | 0.125 (0.065, 0.185) | <0.001 | -0.005 (-0.064, 0.055) | 0.875 |
| | R | 3560 | 0.300 (0.187, 0.413) | <0.001 | 0.059 (-0.102, 0.219) | 0.474 | 0.608 (0.500, 0.715) | <0.001 | 0.008 (-0.107, 0.124) | 0.886 | 0.131 (0.070, 0.192) | <0.001 | -0.009 (-0.069, 0.052) | 0.779 |
| **Pallidum** | L | 3531 | 0.277 (0.151, 0.403) | <0.001 | 0.030 (-0.131, 0.190) | 0.717 | 0.638 (0.529, 0.748) | <0.001 | -0.031 (-0.155, 0.093) | 0.626 | 0.048 (-0.013, 0.109) | 0.125 | 0.038 (-0.023, 0.100) | 0.223 |
| | R | 3536 | 0.275 (0.149, 0.401) | <0.001 | 0.021 (-0.140, 0.182) | 0.802 | 0.648 (0.539, 0.757) | <0.001 | -0.052 (-0.173, 0.069) | 0.397 | 0.026 (-0.034, 0.087) | 0.392 | 0.042 (-0.020, 0.103) | 0.184 |
| **Hippocampus** | L | 3542 | 0.270 (0.164, 0.376) | <0.001 | 0.020 (-0.142, 0.182) | 0.805 | 0.680 (0.579, 0.781) | <0.001 | 0.036 (-0.072, 0.145) | 0.511 | 0.031 (-0.033, 0.095) | 0.337 | 0.056 (-0.008, 0.119) | 0.087 |
| | R | 3557 | 0.251 (0.142, 0.360) | <0.001 | 0.020 (-0.139, 0.180) | 0.803 | 0.701 (0.599, 0.802) | <0.001 | 0.017 (-0.091, 0.125) | 0.754 | 0.010 (-0.052, 0.073) | 0.743 | 0.044 (-0.017, 0.105) | 0.158 |
| **Amygdala** | L | 3542 | 0.040 (-0.060, 0.141) | 0.433 | 0.053 (-0.103, 0.210) | 0.505 | 0.764 (0.665, 0.863) | <0.001 | -0.013 (-0.116, 0.090) | 0.807 | -0.001 (-0.064, 0.063) | 0.986 | 0.097 (0.032, 0.163) | 0.004 |
| | R | 3543 | 0.017 (-0.086, 0.120) | 0.745 | 0.066 (-0.091, 0.222) | 0.411 | 0.780 (0.680, 0.881) | <0.001 | 0.001 (-0.102, 0.105) | 0.980 | 0.005 (-0.057, 0.067) | 0.872 | 0.032 (-0.030, 0.094) | 0.306 |
| **Accumbens** | L | 3548 | 0.411 (0.300, 0.523) | <0.001 | 0.044 (-0.118, 0.206) | 0.596 | 0.624 (0.524, 0.724) | <0.001 | 0.032 (-0.077, 0.140) | 0.565 | 0.122 (0.059, 0.184) | <0.001 | -0.014 (-0.076, 0.048) | 0.651 |
| | R | 3538 | 0.353 (0.244, 0.461) | <0.001 | 0.051 (-0.109, 0.211) | 0.534 | 0.639 (0.538, 0.739) | <0.001 | 0.020 (-0.087, 0.128) | 0.713 | 0.127 (0.065, 0.188) | <0.001 | -0.014 (-0.075, 0.047) | 0.654 |
| **Ventral DC** | L | 3532 | 0.095 (-0.020, 0.210) | 0.108 | 0.036 (-0.123, 0.195) | 0.657 | 0.755 (0.656, 0.854) | <0.001 | -0.061 (-0.176, 0.054) | 0.300 | -0.035 (-0.098, 0.028) | 0.280 | 0.054 (-0.011, 0.119) | 0.102 |
| | R | 3539 | 0.113 (-0.001, 0.228) | 0.053 | 0.030 (-0.128, 0.189) | 0.709 | 0.750 (0.651, 0.849) | <0.001 | 0.011 (-0.102, 0.124) | 0.851 | -0.036 (-0.099, 0.027) | 0.267 | 0.066 (0.003, 0.129) | 0.039 |

**Note:** Estimated regression coefficients (β), 95% confidence intervals (CI) and *p*-values from the linear mixed-effects models evaluating the moderating effects of O3 in the relationship between brain microstructure and Body Mass Index. Models were controlled for age, education, race, ethnicity, puberty, co-exposure to other air pollutants, head motion during the MRI acquisition and urbanicity. Random intercepts were modelled from MRI scanner device and subject ID. Although all two-way interactions were assessed, this table presents only the ones of primary interest due to space constraints. Abbreviations: ROI: region-of-interest; Ventral DC: ventral diencephalon; RSI: restricted spectrum imaging; RNI: restricted normalized isotropic diffusion; H: hemisphere. *p-values < 0.05 of two-way and three-way interactions that achieved the Benjamini-Hochberg False Discovery Rate threshold correction.

**Supplementary Table 13.** Main effects and interactions for the Oxwt measure in females in RNI.

| | | BMI ~ $ROI_{RNI}$ * Oxwt * age + caregiver highest education + race + ethnicity + puberty + PM2.5 + head motion + urbanicity address 1 + (1\|MRI device) + (1\| Subject ID) | | | | | | | | | | | | |
|---|---|---|---|---|---|---|---|---|---|---|---|---|---|---|
| | | | $ROI_{RNI}$ | | Oxwt | | Age | | $ROI_{RNI}$ X Oxwt | | $ROI_{RNI}$ x Age | | $ROI_{RNI}$ x Oxwt x Age | |
| **ROI** | **H** | **n** | **β (95%CI)** | **p-value** | **β (95%CI)** | **p-value** | **β (95%CI)** | **p-value** | **β (95%CI)** | **p-value** | **β (95%CI)** | **p-value** | **β (95%CI)** | **p-value** |
| **Thalamus** | L | 3523 | 0.163 (0.042, 0.284) | 0.008 | 0.034 (-0.131, 0.199) | 0.689 | 0.726 (0.624, 0.827) | <0.001 | -0.007 (-0.131, 0.116) | 0.908 | 0.006 (-0.057, 0.069) | 0.845 | 0.057 (-0.010, 0.124) | 0.097 |
| | R | 3537 | 0.254 (0.135, 0.372) | <0.001 | 0.020 (-0.148, 0.188) | 0.819 | 0.696 (0.594, 0.797) | <0.001 | -0.059 (-0.174, 0.057) | 0.320 | 0.027 (-0.036, 0.090) | 0.403 | 0.018 (-0.046, 0.082) | 0.577 |
| **Caudate** | L | 3566 | 0.353 (0.254, 0.452) | <0.001 | 0.052 (-0.112, 0.215) | 0.537 | 0.660 (0.563, 0.758) | <0.001 | 0.033 (-0.067, 0.133) | 0.521 | 0.086 (0.024, 0.147) | 0.006 | -0.041 (-0.105, 0.023) | 0.208 |
| | R | 3553 | 0.235 (0.137, 0.333) | <0.001 | 0.074 (-0.090, 0.239) | 0.377 | 0.702 (0.604, 0.799) | <0.001 | -0.072 (-0.172, 0.028) | 0.161 | 0.122 (0.060, 0.184) | <0.001 | -0.062 (-0.126, 0.002) | 0.059 |
| **Putamen** | L | 3557 | 0.348 (0.231, 0.466) | <0.001 | 0.046 (-0.124, 0.215) | 0.596 | 0.600 (0.493, 0.708) | <0.001 | 0.008 (-0.110, 0.127) | 0.892 | 0.123 (0.063, 0.182) | <0.001 | -0.007 (-0.067, 0.053) | 0.818 |
| | R | 3560 | 0.307 (0.194, 0.420) | <0.001 | 0.060 (-0.106, 0.226) | 0.479 | 0.609 (0.501, 0.717) | <0.001 | 0.022 (-0.095, 0.140) | 0.707 | 0.128 (0.067, 0.189) | <0.001 | -0.022 (-0.083, 0.039) | 0.488 |
| **Pallidum** | L | 3531 | 0.288 (0.162, 0.415) | <0.001 | 0.014 (-0.152, 0.180) | 0.866 | 0.636 (0.526, 0.745) | <0.001 | 0.029 (-0.096, 0.154) | 0.647 | 0.048 (-0.013, 0.109) | 0.123 | 0.049 (-0.013, 0.111) | 0.118 |
| | R | 3536 | 0.283 (0.157, 0.409) | <0.001 | 0.013 (-0.153, 0.180) | 0.876 | 0.646 (0.537, 0.755) | <0.001 | -0.003 (-0.130, 0.123) | 0.961 | 0.028 (-0.032, 0.089) | 0.356 | 0.058 (-0.003, 0.120) | 0.063 |
| **Hippocampus** | L | 3542 | 0.273 (0.167, 0.379) | <0.001 | 0.033 (-0.135, 0.201) | 0.698 | 0.680 (0.580, 0.781) | <0.001 | -0.003 (-0.111, 0.105) | 0.956 | 0.030 (-0.034, 0.094) | 0.360 | 0.035 (-0.028, 0.099) | 0.274 |
| | R | 3557 | 0.258 (0.149, 0.367) | <0.001 | 0.026 (-0.140, 0.191) | 0.761 | 0.700 (0.599, 0.801) | <0.001 | -0.009 (-0.117, 0.100) | 0.875 | 0.012 (-0.050, 0.074) | 0.711 | 0.054 (-0.006, 0.115) | 0.080 |
| **Amygdala** | L | 3542 | 0.042 (-0.059, 0.143) | 0.415 | 0.054 (-0.110, 0.217) | 0.519 | 0.765 (0.666, 0.865) | <0.001 | -0.038 (-0.140, 0.063) | 0.461 | -0.003 (-0.067, 0.061) | 0.920 | 0.058 (-0.007, 0.122) | 0.080 |
| | R | 3543 | 0.023 (-0.081, 0.127) | 0.665 | 0.065 (-0.098, 0.228) | 0.436 | 0.780 (0.679, 0.880) | <0.001 | -0.037 (-0.143, 0.069) | 0.496 | 0.005 (-0.057, 0.067) | 0.871 | 0.012 (-0.050, 0.073) | 0.715 |
| **Accumbens** | L | 3548 | 0.417 (0.305, 0.529) | <0.001 | 0.059 (-0.108, 0.227) | 0.488 | 0.625 (0.525, 0.725) | <0.001 | 0.049 (-0.058, 0.156) | 0.373 | 0.118 (0.056, 0.181) | <0.001 | -0.017 (-0.079, 0.045) | 0.596 |
| | R | 3538 | 0.358 (0.249, 0.467) | <0.001 | 0.055 (-0.110, 0.220) | 0.511 | 0.640 (0.539, 0.740) | <0.001 | 0.032 (-0.077, 0.141) | 0.565 | 0.123 (0.062, 0.185) | <0.001 | -0.022 (-0.083, 0.038) | 0.473 |
| **Ventral DC** | L | 3532 | 0.106 (-0.010, 0.223) | 0.074 | 0.033 (-0.132, 0.198) | 0.693 | 0.756 (0.657, 0.855) | <0.001 | -0.030 (-0.145, 0.084) | 0.603 | -0.027 (-0.090, 0.036) | 0.402 | 0.058 (-0.006, 0.121) | 0.076 |
| | R | 3539 | 0.127 (0.011, 0.243) | 0.032 | 0.022 (-0.142, 0.186) | 0.794 | 0.751 (0.652, 0.850) | <0.001 | 0.008 (-0.108, 0.124) | 0.897 | -0.029 (-0.092, 0.034) | 0.364 | 0.070 (0.008, 0.132) | 0.026 |

**Note:** Estimated regression coefficients (β), 95% confidence intervals (CI) and *p*-values from the linear mixed-effects models evaluating the moderating effects of Oxwt in the relationship between brain microstructure and Body Mass Index. Models were controlled for age, education, race, ethnicity, puberty, co-exposure to other air pollutants, head motion during the MRI acquisition and urbanicity. Random intercepts were modelled from MRI scanner device and subject ID. Although all two-way interactions were assessed, this table presents only the ones of primary interest due to space constraints. Abbreviations: ROI: region-of-interest; Ventral DC: ventral diencephalon; RSI: restricted spectrum imaging; RNI: restricted normalized isotropic diffusion; H: hemisphere. *p-values < 0.05 of two-way and three-way interactions that achieved the Benjamini-Hochberg False Discovery Rate threshold correction.

**Supplementary Table 14.** Main effects and interactions for the PM2.5 pollutant across both sexes in RND.

| | | BMI ~ $ROI_{RND}$ * PM2.5 * age + sex + caregiver highest education + race + ethnicity + puberty + NO2 + O3 + head motion + urbanicity address 1 + (1\|MRI device) + (1\| Subject ID) | | | | | | | | | | | | |
|---|---|---|---|---|---|---|---|---|---|---|---|---|---|---|
| | | | $ROI_{RND}$ | | PM2.5 | | Age | | $ROI_{RND}$ X PM2.5 | | $ROI_{RND}$ x Age | | $ROI_{RND}$ x PM2.5 x Age | |
| **ROI** | **H** | **n** | **β (95%CI)** | **p-value** | **β (95%CI)** | **p-value** | **β (95%CI)** | **p-value** | **β (95%CI)** | **p-value** | **β (95%CI)** | **p-value** | **β (95%CI)** | **p-value** |
| **Thalamus** | L | 7217 | -0.132 (-0.207, -0.056) | 0.001 | 0.146 (0.006, 0.285) | 0.042 | 0.881 (0.825, 0.937) | <0.001 | -0.025 (-0.099, 0.049) | 0.510 | -0.025 (-0.067, 0.017) | 0.249 | -0.006 (-0.045, 0.033) | 0.765 |
| | R | 7198 | -0.082 (-0.158, -0.006) | 0.035 | 0.147 (0.008, 0.285) | 0.040 | 0.876 (0.820, 0.932) | <0.001 | 0.020 (-0.054, 0.095) | 0.595 | -0.001 (-0.044, 0.041) | 0.949 | 0.020 (-0.020, 0.059) | 0.335 |
| **Caudate** | L | 7097 | 0.155 (0.071, 0.238) | <0.001 | 0.128 (-0.012, 0.268) | 0.076 | 0.859 (0.803, 0.915) | <0.001 | 0.005 (-0.086, 0.097) | 0.911 | 0.094 (0.041, 0.148) | 0.001 | 0.028 (-0.034, 0.090) | 0.374 |
| | R | 7072 | 0.017 (-0.067, 0.101) | 0.690 | 0.145 (0.005, 0.286) | 0.044 | 0.856 (0.800, 0.912) | <0.001 | 0.023 (-0.066, 0.112) | 0.616 | 0.028 (-0.025, 0.080) | 0.308 | 0.077 (0.018, 0.137) | **0.011** |
| **Putamen** | L | 7102 | 0.286 (0.189, 0.383) | <0.001 | 0.102 (-0.044, 0.249) | 0.172 | 0.842 (0.786, 0.899) | <0.001 | 0.027 (-0.073, 0.127) | 0.598 | 0.114 (0.061, 0.167) | <0.001 | 0.094 (0.033, 0.155) | **0.002** |
| | R | 7054 | 0.244 (0.152, 0.336) | <0.001 | 0.101 (-0.043, 0.245) | 0.172 | 0.837 (0.781, 0.894) | <0.001 | 0.059 (-0.039, 0.157) | 0.235 | 0.101 (0.047, 0.156) | <0.001 | 0.114 (0.052, 0.176) | **<0.001** |
| **Pallidum** | L | 7122 | 0.110 (0.042, 0.178) | 0.002 | 0.136 (-0.002, 0.274) | 0.055 | 0.843 (0.787, 0.899) | <0.001 | 0.074 (0.002, 0.146) | 0.043 | 0.099 (0.052, 0.146) | <0.001 | 0.083 (0.038, 0.128) | **<0.001** |
| | R | 7081 | 0.213 (0.141, 0.286) | <0.001 | 0.117 (-0.024, 0.258) | 0.106 | 0.848 (0.791, 0.905) | <0.001 | -0.004 (-0.081, 0.072) | 0.913 | 0.139 (0.087, 0.191) | <0.001 | 0.067 (0.016, 0.117) | **0.010** |
| **Hippocampus** | L | 7138 | -0.014 (-0.097, 0.069) | 0.740 | 0.120 (-0.022, 0.263) | 0.100 | 0.854 (0.799, 0.910) | <0.001 | -0.116 (-0.202, -0.029) | 0.009 | 0.012 (-0.035, 0.059) | 0.617 | 0.007 (-0.041, 0.055) | 0.771 |
| | R | 7136 | -0.030 (-0.109, 0.049) | 0.456 | 0.134 (-0.006, 0.274) | 0.062 | 0.855 (0.800, 0.911) | <0.001 | -0.052 (-0.137, 0.033) | 0.227 | -0.022 (-0.069, 0.024) | 0.348 | 0.011 (-0.038, 0.060) | 0.656 |
| **Amygdala** | L | 7129 | -0.017 (-0.093, 0.060) | 0.667 | 0.146 (0.006, 0.286) | 0.042 | 0.850 (0.794, 0.905) | <0.001 | -0.077 (-0.158, 0.004) | 0.063 | -0.005 (-0.051, 0.042) | 0.840 | -0.004 (-0.054, 0.047) | 0.888 |
| | R | 7163 | -0.061 (-0.139, 0.018) | 0.131 | 0.157 (0.018, 0.296) | 0.028 | 0.856 (0.801, 0.911) | <0.001 | -0.003 (-0.085, 0.080) | 0.952 | 0.0002 (-0.046, 0.047) | 0.991 | 0.048 (-0.004, 0.099) | 0.069 |
| **Accumbens** | L | 7081 | 0.047 (-0.034, 0.128) | 0.260 | 0.133 (-0.006, 0.273) | 0.063 | 0.858 (0.802, 0.914) | <0.001 | 0.056 (-0.030, 0.142) | 0.201 | 0.063 (0.013, 0.114) | 0.015 | 0.029 (-0.027, 0.085) | 0.303 |
| | R | 7072 | 0.125 (0.038, 0.212) | 0.005 | 0.121 (-0.020, 0.262) | 0.096 | 0.857 (0.800, 0.913) | <0.001 | -0.002 (-0.094, 0.089) | 0.964 | 0.073 (0.022, 0.124) | 0.005 | 0.011 (-0.046, 0.069) | 0.707 |
| **Ventral DC** | L | 7170 | -0.201 (-0.282, -0.120) | <0.001 | 0.117 (-0.028, 0.262) | 0.115 | 0.883 (0.827, 0.938) | <0.001 | -0.073 (-0.148, 0.002) | 0.058 | -0.035 (-0.077, 0.007) | 0.106 | -0.005 (-0.041, 0.030) | 0.762 |
| | R | 7164 | -0.195 (-0.275, -0.114) | <0.001 | 0.131 (-0.011, 0.273) | 0.072 | 0.889 (0.834, 0.945) | <0.001 | -0.016 (-0.092, 0.060) | 0.680 | -0.055 (-0.097, -0.012) | 0.011 | 0.011 (-0.025, 0.048) | 0.542 |

**Note:** Estimated regression coefficients (β), 95% confidence intervals (CI) and *p*-values from the linear mixed-effects models evaluating the moderating effects of PM2.5 in the relationship between brain microstructure and Body Mass Index. Models were controlled for age, sex, education, race, ethnicity, puberty, co-exposure to other air pollutants, head motion during the MRI acquisition and urbanicity. Random intercepts were modelled from MRI scanner device and subject ID. Although all two-way interactions were assessed, this table presents only the ones of primary interest due to space constraints. Abbreviations: ROI: region-of-interest; Ventral DC: ventral diencephalon; RSI: restricted spectrum imaging; RND: restricted normalized directional diffusion; H: hemisphere. *p-values < 0.05 of two-way and three-way interactions that achieved the Benjamini-Hochberg False Discovery Rate threshold correction.

**Supplementary Table 15.** Main effects and interactions for the NO2 pollutant across both sexes in RND.

| | | BMI ~ $ROI_{RND}$ * NO2 * age + sex + caregiver highest education + race + ethnicity + puberty + PM2.5 + O3 + head motion + urbanicity address 1 + (1\|MRI device) + (1\| Subject ID) | | | | | | | | | | | | |
|---|---|---|---|---|---|---|---|---|---|---|---|---|---|---|
| | | | $ROI_{RND}$ | | NO2 | | Age | | $ROI_{RND}$ X NO2 | | $ROI_{RND}$ x Age | | $ROI_{RND}$ x NO2 x Age | |
| **ROI** | **H** | **n** | **β (95%CI)** | **p-value** | **β (95%CI)** | **p-value** | **β (95%CI)** | **p-value** | **β (95%CI)** | **p-value** | **β (95%CI)** | **p-value** | **β (95%CI)** | **p-value** |
| **Thalamus** | L | 7217 | -0.141 (-0.216, -0.066) | <0.001 | 0.058 (-0.086, 0.201) | 0.431 | 0.884 (0.828, 0.940) | <0.001 | 0.010 (-0.068, 0.087) | 0.809 | -0.024 (-0.066, 0.018) | 0.267 | -0.029 (-0.072, 0.014) | 0.182 |
| | R | 7198 | -0.084 (-0.159, -0.008) | 0.030 | 0.066 (-0.078, 0.210) | 0.369 | 0.877 (0.821, 0.934) | <0.001 | 0.025 (-0.053, 0.104) | 0.527 | -0.001 (-0.044, 0.042) | 0.960 | -0.016 (-0.059, 0.027) | 0.464 |
| **Caudate** | L | 7097 | 0.143 (0.061, 0.224) | 0.001 | 0.078 (-0.066, 0.222) | 0.289 | 0.859 (0.803, 0.914) | <0.001 | 0.063 (-0.017, 0.143) | 0.124 | 0.107 (0.055, 0.159) | <0.001 | -0.025 (-0.081, 0.032) | 0.390 |
| | R | 7072 | -0.0004 (-0.083, 0.082) | 0.992 | 0.059 (-0.086, 0.204) | 0.428 | 0.859 (0.804, 0.914) | <0.001 | 0.007 (-0.077, 0.091) | 0.871 | 0.045 (-0.007, 0.096) | 0.089 | -0.053 (-0.108, 0.002) | 0.057 |
| **Putamen** | L | 7102 | 0.272 (0.178, 0.365) | <0.001 | 0.105 (-0.044, 0.253) | 0.168 | 0.856 (0.801, 0.911) | <0.001 | 0.118 (0.024, 0.212) | 0.014 | 0.135 (0.084, 0.187) | <0.001 | 0.014 (-0.040, 0.069) | 0.607 |
| | R | 7054 | 0.227 (0.138, 0.316) | <0.001 | 0.089 (-0.058, 0.235) | 0.239 | 0.849 (0.794, 0.904) | <0.001 | 0.097 (0.006, 0.188) | 0.036 | 0.127 (0.074, 0.179) | <0.001 | -0.002 (-0.058, 0.054) | 0.942 |
| **Pallidum** | L | 7122 | 0.099 (0.031, 0.166) | 0.004 | 0.058 (-0.085, 0.201) | 0.427 | 0.847 (0.792, 0.903) | <0.001 | 0.102 (0.032, 0.171) | 0.004 | 0.103 (0.057, 0.149) | <0.001 | 0.020 (-0.025, 0.065) | 0.385 |
| | R | 7081 | 0.189 (0.119, 0.260) | <0.001 | 0.054 (-0.090, 0.198) | 0.461 | 0.858 (0.802, 0.914) | <0.001 | 0.077 (0.005, 0.149) | 0.036 | 0.154 (0.103, 0.204) | <0.001 | 0.026 (-0.026, 0.078) | 0.327 |
| **Hippocampus** | L | 7138 | -0.048 (-0.128, 0.032) | 0.238 | 0.050 (-0.093, 0.192) | 0.496 | 0.861 (0.805, 0.916) | <0.001 | 0.038 (-0.047, 0.123) | 0.382 | 0.019 (-0.028, 0.066) | 0.424 | -0.041 (-0.089, 0.008) | 0.099 |
| | R | 7136 | -0.051 (-0.128, 0.027) | 0.198 | 0.055 (-0.088, 0.198) | 0.450 | 0.857 (0.802, 0.912) | <0.001 | 0.022 (-0.057, 0.101) | 0.581 | -0.013 (-0.059, 0.033) | 0.572 | -0.010 (-0.056, 0.036) | 0.666 |
| **Amygdala** | L | 7129 | -0.029 (-0.104, 0.047) | 0.454 | 0.056 (-0.088, 0.200) | 0.447 | 0.854 (0.798, 0.909) | <0.001 | 0.078 (0.003, 0.154) | 0.043 | 0.005 (-0.041, 0.050) | 0.838 | -0.009 (-0.055, 0.038) | 0.711 |
| | R | 7163 | -0.070 (-0.147, 0.008) | 0.080 | 0.057 (-0.087, 0.201) | 0.441 | 0.860 (0.805, 0.915) | <0.001 | 0.010 (-0.069, 0.088) | 0.807 | 0.011 (-0.035, 0.057) | 0.631 | -0.044 (-0.090, 0.002) | 0.061 |
| **Accumbens** | L | 7081 | 0.046 (-0.034, 0.125) | 0.260 | 0.072 (-0.072, 0.216) | 0.331 | 0.858 (0.803, 0.913) | <0.001 | 0.034 (-0.047, 0.116) | 0.410 | 0.071 (0.022, 0.120) | 0.005 | -0.014 (-0.066, 0.037) | 0.585 |
| | R | 7072 | 0.118 (0.033, 0.203) | 0.006 | 0.077 (-0.069, 0.222) | 0.303 | 0.857 (0.802, 0.912) | <0.001 | 0.046 (-0.042, 0.134) | 0.309 | 0.082 (0.033, 0.131) | 0.001 | -0.019 (-0.072, 0.033) | 0.471 |
| **Ventral DC** | L | 7170 | -0.213 (-0.293, -0.132) | <0.001 | 0.068 (-0.078, 0.214) | 0.361 | 0.885 (0.829, 0.941) | <0.001 | 0.011 (-0.069, 0.092) | 0.779 | -0.038 (-0.081, 0.006) | 0.089 | -0.002 (-0.044, 0.040) | 0.936 |
| | R | 7164 | -0.195 (-0.276, -0.114) | <0.001 | 0.076 (-0.069, 0.222) | 0.305 | 0.889 (0.833, 0.945) | <0.001 | 0.016 (-0.064, 0.096) | 0.691 | -0.059 (-0.103, -0.016) | 0.007 | 0.0001 (-0.041, 0.041) | 0.996 |

**Note:** Estimated regression coefficients (β), 95% confidence intervals (CI) and *p*-values from the linear mixed-effects models evaluating the moderating effects of NO2 in the relationship between brain microstructure and Body Mass Index. Models were controlled for age, sex, education, race, ethnicity, puberty, co-exposure to other air pollutants, head motion during the MRI acquisition and urbanicity. Random intercepts were modelled from MRI scanner device and subject ID. Although all two-way interactions were assessed, this table presents only the ones of primary interest due to space constraints. Abbreviations: ROI: region-of-interest; Ventral DC: ventral diencephalon; RSI: restricted spectrum imaging; RND: restricted normalized directional diffusion; H: hemisphere. *p-values < 0.05 of two-way and three-way interactions that achieved the Benjamini-Hochberg False Discovery Rate threshold correction.

**Supplementary Table 16.** Main effects and interactions for the O3 pollutant across both sexes in RND.

| | | BMI ~ ROI$_{RND}$ * O3 * age + sex + caregiver highest education + race + ethnicity + puberty + PM2.5 + NO2 + head motion + urbanicity address 1 + (1\|MRI device) + (1\| Subject ID) | | | | | | | | | | | | |
|---|---|---|---|---|---|---|---|---|---|---|---|---|---|---|
| | | | ROI$_{RND}$ | | O3 | | Age | | ROI$_{RND}$ X O3 | | ROI$_{RND}$ x Age | | ROI$_{RND}$ x O3 x Age | |
| ROI | H | n | β (95%CI) | p-value | β (95%CI) | p-value | β (95%CI) | p-value | β (95%CI) | p-value | β (95%CI) | p-value | β (95%CI) | p-value |
| Thalamus | L | 7217 | -0.141 (-0.216, -0.066) | <0.001 | 0.024 (-0.094, 0.143) | 0.686 | 0.880 (0.824, 0.935) | <0.001 | 0.032 (-0.042, 0.107) | 0.397 | -0.020 (-0.062, 0.022) | 0.355 | -0.009 (-0.052, 0.033) | 0.672 |
| | R | 7198 | -0.085 (-0.161, -0.009) | 0.028 | 0.028 (-0.091, 0.147) | 0.650 | 0.873 (0.817, 0.929) | <0.001 | -0.046 (-0.122, 0.030) | 0.232 | 0.002 (-0.040, 0.045) | 0.923 | -0.026 (-0.069, 0.017) | 0.233 |
| Caudate | L | 7097 | 0.156 (0.072, 0.239) | <0.001 | 0.060 (-0.059, 0.179) | 0.324 | 0.858 (0.802, 0.913) | <0.001 | 0.038 (-0.042, 0.118) | 0.352 | 0.100 (0.047, 0.154) | <0.001 | -0.017 (-0.071, 0.037) | 0.539 |
| | R | 7072 | -0.004 (-0.087, 0.080) | 0.934 | 0.034 (-0.085, 0.154) | 0.573 | 0.860 (0.805, 0.916) | <0.001 | -0.050 (-0.130, 0.029) | 0.215 | 0.043 (-0.010, 0.095) | 0.110 | -0.036 (-0.089, 0.017) | 0.184 |
| Putamen | L | 7102 | 0.282 (0.186, 0.377) | <0.001 | 0.061 (-0.061, 0.182) | 0.326 | 0.847 (0.791, 0.902) | <0.001 | 0.055 (-0.030, 0.140) | 0.205 | 0.128 (0.076, 0.180) | <0.001 | -0.020 (-0.072, 0.032) | 0.449 |
| | R | 7054 | 0.242 (0.152, 0.333) | <0.001 | 0.072 (-0.049, 0.193) | 0.245 | 0.837 (0.781, 0.893) | <0.001 | 0.052 (-0.031, 0.135) | 0.221 | 0.116 (0.063, 0.170) | <0.001 | -0.039 (-0.092, 0.014) | 0.150 |
| Pallidum | L | 7122 | 0.099 (0.031, 0.167) | 0.005 | 0.051 (-0.068, 0.170) | 0.402 | 0.851 (0.795, 0.908) | <0.001 | -0.011 (-0.076, 0.053) | 0.729 | 0.101 (0.055, 0.147) | <0.001 | -0.029 (-0.075, 0.017) | 0.216 |
| | R | 7081 | 0.214 (0.141, 0.286) | <0.001 | 0.055 (-0.065, 0.175) | 0.369 | 0.850 (0.794, 0.906) | <0.001 | 0.066 (-0.001, 0.133) | 0.055 | 0.148 (0.097, 0.199) | <0.001 | -0.036 (-0.087, 0.015) | 0.164 |
| Hippocampus | L | 7138 | -0.048 (-0.128, 0.033) | 0.244 | 0.040 (-0.079, 0.159) | 0.512 | 0.859 (0.803, 0.915) | <0.001 | 0.001 (-0.076, 0.079) | 0.971 | 0.020 (-0.027, 0.068) | 0.406 | -0.002 (-0.050, 0.047) | 0.943 |
| | R | 7136 | -0.047 (-0.125, 0.031) | 0.233 | 0.033 (-0.085, 0.152) | 0.582 | 0.850 (0.794, 0.906) | <0.001 | -0.019 (-0.094, 0.056) | 0.626 | -0.020 (-0.067, 0.027) | 0.409 | -0.041 (-0.088, 0.006) | 0.090 |
| Amygdala | L | 7129 | -0.028 (-0.104, 0.048) | 0.470 | 0.034 (-0.085, 0.153) | 0.575 | 0.847 (0.791, 0.902) | <0.001 | 0.012 (-0.061, 0.086) | 0.742 | -0.0002 (-0.047, 0.046) | 0.993 | -0.026 (-0.075, 0.022) | 0.283 |
| | R | 7163 | -0.070 (-0.148, 0.009) | 0.081 | 0.031 (-0.088, 0.150) | 0.609 | 0.855 (0.800, 0.911) | <0.001 | -0.017 (-0.094, 0.060) | 0.659 | 0.008 (-0.039, 0.055) | 0.728 | -0.024 (-0.072, 0.023) | 0.314 |
| Accumbens | L | 7081 | 0.039 (-0.042, 0.120) | 0.347 | 0.046 (-0.073, 0.165) | 0.451 | 0.859 (0.803, 0.915) | <0.001 | -0.061 (-0.138, 0.017) | 0.124 | 0.068 (0.018, 0.119) | 0.008 | -0.036 (-0.087, 0.015) | 0.165 |
| | R | 7072 | 0.122 (0.035, 0.208) | 0.006 | 0.054 (-0.066, 0.174) | 0.381 | 0.853 (0.798, 0.909) | <0.001 | -0.011 (-0.095, 0.073) | 0.796 | 0.077 (0.026, 0.128) | 0.003 | -0.034 (-0.086, 0.017) | 0.195 |
| Ventral DC | L | 7170 | -0.212 (-0.293, -0.132) | <0.001 | 0.008 (-0.111, 0.128) | 0.891 | 0.884 (0.828, 0.939) | <0.001 | 0.032 (-0.045, 0.109) | 0.418 | -0.035 (-0.078, 0.008) | 0.109 | -0.004 (-0.047, 0.038) | 0.844 |
| | R | 7164 | -0.196 (-0.277, -0.116) | <0.001 | 0.009 (-0.111, 0.128) | 0.884 | 0.891 (0.835, 0.947) | <0.001 | 0.049 (-0.029, 0.126) | 0.216 | -0.060 (-0.103, -0.017) | 0.006 | 0.021 (-0.022, 0.064) | 0.348 |

**Note:** Estimated regression coefficients (β), 95% confidence intervals (CI) and *p*-values from the linear mixed-effects models evaluating the moderating effects of O3 in the relationship between brain microstructure and Body Mass Index. Models were controlled for age, sex, education, race, ethnicity, puberty, co-exposure to other air pollutants, head motion during the MRI acquisition and urbanicity. Random intercepts were modelled from MRI scanner device and subject ID. Although all two-way interactions were assessed, this table presents only the ones of primary interest due to space constraints. Abbreviations: ROI: region-of-interest; Ventral DC: ventral diencephalon; RSI: restricted spectrum imaging; RND: restricted normalized directional diffusion; H: hemisphere. *p-values < 0.05 of two-way and three-way interactions that achieved the Benjamini-Hochberg False Discovery Rate threshold correction.

**Supplementary Table 17.** Main effects and interactions for the Oxwt measure across both sexes in RND.

| | | BMI ~ $ROI_{RND}$ * Oxwt * age + sex + caregiver highest education + race + ethnicity + puberty + PM2.5 + head motion + urbanicity address 1 + (1\|MRI device) + (1\| Subject ID) | | | | | | | | | | | | |
|---|---|---|---|---|---|---|---|---|---|---|---|---|---|---|
| | | | $ROI_{RND}$ | | Oxwt | | Age | | $ROI_{RND}$ X Oxwt | | $ROI_{RND}$ x Age | | $ROI_{RND}$ x Oxwt x Age | |
| **ROI** | **H** | **n** | **β (95%CI)** | **p-value** | **β (95%CI)** | **p-value** | **β (95%CI)** | **p-value** | **β (95%CI)** | **p-value** | **β (95%CI)** | **p-value** | **β (95%CI)** | **p-value** |
| **Thalamus** | L | 7217 | -0.142 (-0.217, -0.067) | <0.001 | 0.048 (-0.075, 0.171) | 0.447 | 0.880 (0.824, 0.935) | <0.001 | 0.032 (-0.042, 0.106) | 0.392 | -0.020 (-0.062, 0.023) | 0.362 | -0.025 (-0.068, 0.017) | 0.240 |
| | R | 7198 | -0.084 (-0.160, -0.009) | 0.029 | 0.056 (-0.067, 0.179) | 0.376 | 0.874 (0.818, 0.930) | <0.001 | -0.025 (-0.101, 0.052) | 0.528 | 0.001 (-0.041, 0.044) | 0.949 | -0.032 (-0.075, 0.011) | 0.148 |
| **Caudate** | L | 7097 | 0.157 (0.074, 0.240) | <0.001 | 0.087 (-0.037, 0.211) | 0.169 | 0.854 (0.799, 0.910) | <0.001 | 0.061 (-0.016, 0.139) | 0.120 | 0.100 (0.046, 0.153) | <0.001 | -0.023 (-0.076, 0.030) | 0.390 |
| | R | 7072 | 0.001 (-0.083, 0.084) | 0.984 | 0.056 (-0.069, 0.180) | 0.381 | 0.857 (0.801, 0.912) | <0.001 | -0.044 (-0.123, 0.035) | 0.274 | 0.040 (-0.013, 0.092) | 0.137 | -0.059 (-0.112, -0.006) | 0.029 |
| **Putamen** | L | 7102 | 0.284 (0.189, 0.380) | <0.001 | 0.100 (-0.027, 0.226) | 0.122 | 0.846 (0.791, 0.902) | <0.001 | 0.097 (0.013, 0.181) | 0.024 | 0.128 (0.076, 0.181) | <0.001 | -0.006 (-0.057, 0.045) | 0.821 |
| | R | 7054 | 0.243 (0.153, 0.334) | <0.001 | 0.103 (-0.023, 0.228) | 0.111 | 0.837 (0.781, 0.893) | <0.001 | 0.085 (0.002, 0.167) | 0.044 | 0.117 (0.064, 0.171) | <0.001 | -0.029 (-0.081, 0.024) | 0.283 |
| **Pallidum** | L | 7122 | 0.104 (0.036, 0.172) | 0.003 | 0.070 (-0.053, 0.193) | 0.265 | 0.849 (0.793, 0.905) | <0.001 | 0.039 (-0.025, 0.104) | 0.231 | 0.101 (0.055, 0.147) | <0.001 | -0.009 (-0.055, 0.037) | 0.699 |
| | R | 7081 | 0.211 (0.139, 0.284) | <0.001 | 0.073 (-0.052, 0.198) | 0.253 | 0.851 (0.795, 0.907) | <0.001 | 0.089 (0.023, 0.155) | 0.008 | 0.149 (0.099, 0.200) | <0.001 | -0.012 (-0.063, 0.038) | 0.628 |
| **Hippocampus** | L | 7138 | -0.044 (-0.125, 0.036) | 0.282 | 0.058 (-0.066, 0.181) | 0.359 | 0.855 (0.800, 0.911) | <0.001 | 0.017 (-0.061, 0.095) | 0.675 | 0.019 (-0.028, 0.066) | 0.430 | -0.024 (-0.072, 0.024) | 0.325 |
| | R | 7136 | -0.047 (-0.125, 0.031) | 0.238 | 0.055 (-0.068, 0.178) | 0.379 | 0.851 (0.795, 0.907) | <0.001 | -0.004 (-0.078, 0.070) | 0.917 | -0.017 (-0.063, 0.030) | 0.477 | -0.037 (-0.082, 0.008) | 0.107 |
| **Amygdala** | L | 7129 | -0.026 (-0.102, 0.050) | 0.496 | 0.053 (-0.071, 0.178) | 0.402 | 0.847 (0.791, 0.902) | <0.001 | 0.049 (-0.025, 0.123) | 0.192 | 0.0002 (-0.046, 0.047) | 0.992 | -0.025 (-0.072, 0.022) | 0.303 |
| | R | 7163 | -0.068 (-0.146, 0.010) | 0.087 | 0.052 (-0.071, 0.175) | 0.408 | 0.853 (0.798, 0.909) | <0.001 | -0.013 (-0.089, 0.064) | 0.746 | 0.006 (-0.040, 0.053) | 0.785 | -0.044 (-0.089, 0.001) | 0.057 |
| **Accumbens** | L | 7081 | 0.039 (-0.042, 0.120) | 0.348 | 0.072 (-0.052, 0.195) | 0.257 | 0.858 (0.802, 0.913) | <0.001 | -0.038 (-0.115, 0.040) | 0.340 | 0.066 (0.015, 0.116) | 0.011 | -0.038 (-0.089, 0.013) | 0.143 |
| | R | 7072 | 0.125 (0.038, 0.211) | 0.005 | 0.081 (-0.044, 0.206) | 0.204 | 0.852 (0.797, 0.908) | <0.001 | 0.012 (-0.072, 0.097) | 0.775 | 0.075 (0.025, 0.126) | 0.004 | -0.037 (-0.087, 0.013) | 0.151 |
| **Ventral DC** | L | 7170 | -0.211 (-0.291, -0.131) | <0.001 | 0.035 (-0.089, 0.159) | 0.577 | 0.884 (0.829, 0.940) | <0.001 | 0.037 (-0.043, 0.117) | 0.359 | -0.034 (-0.076, 0.009) | 0.118 | -0.005 (-0.049, 0.039) | 0.830 |
| | R | 7164 | -0.193 (-0.273, -0.113) | <0.001 | 0.040 (-0.084, 0.163) | 0.530 | 0.888 (0.832, 0.944) | <0.001 | 0.054 (-0.026, 0.133) | 0.186 | -0.055 (-0.098, -0.013) | 0.011 | 0.017 (-0.026, 0.060) | 0.436 |

**Note:** Estimated regression coefficients (β), 95% confidence intervals (CI) and *p*-values from the linear mixed-effects models evaluating the moderating effects of Oxwt in the relationship between brain microstructure and Body Mass Index. Models were controlled for age, sex, education, race, ethnicity, puberty, co-exposure to other air pollutants, head motion during the MRI acquisition and urbanicity. Random intercepts were modelled from MRI scanner device and subject ID. Although all two-way interactions were assessed, this table presents only the ones of primary interest due to space constraints. Abbreviations: ROI: region-of-interest; Ventral DC: ventral diencephalon; RSI: restricted spectrum imaging; RND: restricted normalized directional diffusion; H: hemisphere. *p-values < 0.05 of two-way and three-way interactions that achieved the Benjamini-Hochberg False Discovery Rate threshold correction.

**Supplementary Table 18.** Main effects and interactions for the PM2.5 pollutant for males in RND.

| | | BMI ~ $ROI_{RND}$ * PM2.5 * age + caregiver highest education + race + ethnicity + puberty + NO2 + O3 + head motion + urbanicity address 1 + (1\|MRI device) + (1\| Subject ID) | | | | | | | | | | | | |
|---|---|---|---|---|---|---|---|---|---|---|---|---|---|---|
| | | | $ROI_{RND}$ | | PM2.5 | | Age | | $ROI_{RND}$ X PM2.5 | | $ROI_{RND}$ x Age | | $ROI_{RND}$ x PM2.5 x Age | |
| **ROI** | **H** | **n** | **β (95%CI)** | **p-value** | **β (95%CI)** | **p-value** | **β (95%CI)** | **p-value** | **β (95%CI)** | **p-value** | **β (95%CI)** | **p-value** | **β (95%CI)** | **p-value** |
| **Thalamus** | L | 3631 | -0.120 (-0.222, -0.017) | 0.022 | 0.218 (0.037, 0.399) | 0.020 | 0.923 (0.854, 0.991) | <0.001 | 0.060 (-0.040, 0.160) | 0.241 | -0.067 (-0.125, -0.010) | 0.022 | -0.007 (-0.059, 0.045) | 0.795 |
| | R | 3617 | -0.141 (-0.244, -0.038) | 0.007 | 0.212 (0.035, 0.389) | 0.020 | 0.929 (0.860, 0.997) | <0.001 | 0.120 (0.020, 0.220) | 0.019 | -0.059 (-0.117, -0.001) | 0.048 | -0.004 (-0.058, 0.050) | 0.877 |
| **Caudate** | L | 3555 | 0.099 (-0.018, 0.215) | 0.096 | 0.192 (0.002, 0.382) | 0.050 | 0.889 (0.820, 0.957) | <0.001 | 0.089 (-0.042, 0.219) | 0.183 | 0.049 (-0.024, 0.122) | 0.187 | 0.066 (-0.020, 0.153) | 0.134 |
| | R | 3550 | -0.035 (-0.148, 0.079) | 0.549 | 0.208 (0.020, 0.396) | 0.032 | 0.896 (0.828, 0.964) | <0.001 | 0.102 (-0.019, 0.223) | 0.099 | -0.029 (-0.099, 0.041) | 0.419 | 0.101 (0.017, 0.184) | 0.018 |
| **Putamen** | L | 3567 | 0.310 (0.180, 0.440) | <0.001 | 0.140 (-0.052, 0.333) | 0.157 | 0.876 (0.807, 0.944) | <0.001 | 0.078 (-0.061, 0.216) | 0.273 | 0.108 (0.036, 0.179) | 0.003 | 0.126 (0.044, 0.207) | 0.002 |
| | R | 3537 | 0.284 (0.161, 0.407) | <0.001 | 0.150 (-0.043, 0.344) | 0.130 | 0.872 (0.804, 0.941) | <0.001 | 0.162 (0.031, 0.294) | 0.016 | 0.064 (-0.009, 0.137) | 0.085 | 0.177 (0.095, 0.258) | <0.001 |
| **Pallidum** | L | 3582 | 0.092 (-0.002, 0.186) | 0.054 | 0.182 (-0.005, 0.368) | 0.058 | 0.879 (0.809, 0.948) | <0.001 | 0.049 (-0.053, 0.150) | 0.349 | 0.043 (-0.021, 0.108) | 0.188 | 0.103 (0.037, 0.169) | 0.002 |
| | R | 3556 | 0.237 (0.137, 0.337) | <0.001 | 0.191 (0.001, 0.380) | 0.050 | 0.886 (0.816, 0.956) | <0.001 | 0.099 (-0.009, 0.207) | 0.073 | 0.142 (0.073, 0.212) | <0.001 | 0.148 (0.079, 0.218) | <0.001 |
| **Hippocampus** | L | 3586 | -0.060 (-0.172, 0.052) | 0.293 | 0.177 (-0.013, 0.367) | 0.070 | 0.892 (0.824, 0.959) | <0.001 | -0.104 (-0.224, 0.015) | 0.087 | -0.033 (-0.096, 0.029) | 0.291 | -0.015 (-0.078, 0.047) | 0.634 |
| | R | 3585 | -0.032 (-0.139, 0.076) | 0.565 | 0.197 (0.012, 0.382) | 0.039 | 0.896 (0.828, 0.963) | <0.001 | 0.014 (-0.100, 0.129) | 0.806 | -0.054 (-0.115, 0.008) | 0.086 | 0.036 (-0.029, 0.102) | 0.276 |
| **Amygdala** | L | 3589 | 0.022 (-0.085, 0.129) | 0.687 | 0.209 (0.025, 0.392) | 0.027 | 0.890 (0.822, 0.958) | <0.001 | -0.001 (-0.118, 0.116) | 0.985 | -0.039 (-0.103, 0.024) | 0.223 | 0.028 (-0.042, 0.099) | 0.432 |
| | R | 3613 | -0.024 (-0.130, 0.082) | 0.656 | 0.205 (0.022, 0.388) | 0.030 | 0.894 (0.826, 0.961) | <0.001 | 0.033 (-0.082, 0.147) | 0.577 | -0.050 (-0.113, 0.013) | 0.121 | 0.055 (-0.016, 0.126) | 0.129 |
| **Accumbens** | L | 3549 | 0.029 (-0.079, 0.137) | 0.601 | 0.175 (-0.012, 0.362) | 0.069 | 0.890 (0.822, 0.958) | <0.001 | 0.074 (-0.046, 0.194) | 0.225 | 0.022 (-0.046, 0.090) | 0.521 | 0.055 (-0.022, 0.132) | 0.163 |
| | R | 3548 | 0.082 (-0.035, 0.200) | 0.171 | 0.182 (-0.005, 0.369) | 0.058 | 0.887 (0.819, 0.956) | <0.001 | 0.076 (-0.052, 0.204) | 0.242 | 0.019 (-0.048, 0.086) | 0.581 | 0.066 (-0.014, 0.146) | 0.108 |
| **Ventral DC** | L | 3613 | -0.162 (-0.272, -0.051) | 0.004 | 0.190 (0.005, 0.376) | 0.046 | 0.922 (0.854, 0.990) | <0.001 | 0.003 (-0.101, 0.106) | 0.957 | -0.066 (-0.123, -0.009) | 0.023 | 0.006 (-0.041, 0.054) | 0.791 |
| | R | 3618 | -0.148 (-0.258, -0.038) | 0.008 | 0.197 (0.015, 0.380) | 0.035 | 0.923 (0.855, 0.991) | <0.001 | 0.063 (-0.041, 0.166) | 0.238 | -0.086 (-0.144, -0.028) | 0.003 | 0.014 (-0.037, 0.065) | 0.600 |

**Note:** Estimated regression coefficients (β), 95% confidence intervals (CI) and *p*-values from the linear mixed-effects models evaluating the moderating effects of PM2.5 in the relationship between brain microstructure and Body Mass Index. Models were controlled for age, education, race, ethnicity, puberty, co-exposure to other air pollutants, head motion during the MRI acquisition and urbanicity. Random intercepts were modelled from MRI scanner device and subject ID. Although all two-way interactions were assessed, this table presents only the ones of primary interest due to space constraints. Abbreviations: ROI: region-of-interest; Ventral DC: ventral diencephalon; RSI: restricted spectrum imaging; RND: restricted normalized directional diffusion; H: hemisphere. *p-values < 0.05 of two-way and three-way interactions that achieved the Benjamini-Hochberg False Discovery Rate threshold correction.

**Supplementary Table 19.** Main effects and interactions for the NO2 pollutant for males in RND.

| | | BMI ~ $ROI_{RND}$ * NO2 * age + caregiver highest education + race + ethnicity + puberty + PM2.5 + O3 + head motion + urbanicity address 1 + (1\|MRI device) + (1\| Subject ID) | | | | | | | | | | | | |
|---|---|---|---|---|---|---|---|---|---|---|---|---|---|---|
| | | | $ROI_{RND}$ | | NO2 | | Age | | $ROI_{RND}$ X NO2 | | $ROI_{RND}$ x Age | | $ROI_{RND}$ x NO2 x Age | |
| **ROI** | **H** | **n** | **β (95%CI)** | **p-value** | **β (95%CI)** | **p-value** | **β (95%CI)** | **p-value** | **β (95%CI)** | **p-value** | **β (95%CI)** | **p-value** | **β (95%CI)** | **p-value** |
| **Thalamus** | L | 3631 | -0.129 (-0.232, -0.027) | 0.013 | 0.010 (-0.176, 0.196) | 0.919 | 0.916 (0.848, 0.985) | <0.001 | 0.054 (-0.052, 0.160) | 0.317 | -0.063 (-0.120, -0.005) | 0.033 | -0.022 (-0.079, 0.036) | 0.462 |
| | R | 3617 | -0.141 (-0.243, -0.038) | 0.007 | 0.009 (-0.176, 0.194) | 0.927 | 0.918 (0.849, 0.987) | <0.001 | 0.085 (-0.021, 0.192) | 0.117 | -0.050 (-0.109, 0.008) | 0.090 | -0.013 (-0.072, 0.046) | 0.663 |
| **Caudate** | L | 3555 | 0.091 (-0.024, 0.205) | 0.121 | 0.049 (-0.145, 0.243) | 0.621 | 0.885 (0.818, 0.953) | <0.001 | 0.067 (-0.047, 0.180) | 0.249 | 0.054 (-0.017, 0.125) | 0.136 | -0.007 (-0.084, 0.069) | 0.854 |
| | R | 3550 | -0.044 (-0.155, 0.067) | 0.439 | 0.020 (-0.175, 0.214) | 0.844 | 0.889 (0.822, 0.956) | <0.001 | 0.017 (-0.100, 0.134) | 0.777 | -0.019 (-0.088, 0.050) | 0.590 | -0.035 (-0.109, 0.040) | 0.362 |
| **Putamen** | L | 3567 | 0.299 (0.173, 0.426) | <0.001 | 0.078 (-0.116, 0.272) | 0.435 | 0.891 (0.823, 0.958) | <0.001 | 0.120 (-0.004, 0.244) | 0.058 | 0.134 (0.064, 0.204) | <0.001 | 0.042 (-0.029, 0.114) | 0.247 |
| | R | 3537 | 0.270 (0.150, 0.391) | <0.001 | 0.061 (-0.134, 0.256) | 0.542 | 0.883 (0.815, 0.950) | <0.001 | 0.115 (-0.010, 0.241) | 0.071 | 0.100 (0.029, 0.170) | 0.006 | 0.063 (-0.010, 0.137) | 0.091 |
| **Pallidum** | L | 3582 | 0.070 (-0.021, 0.162) | 0.133 | 0.033 (-0.157, 0.223) | 0.736 | 0.884 (0.816, 0.953) | <0.001 | 0.092 (-0.004, 0.187) | 0.059 | 0.053 (-0.010, 0.116) | 0.102 | 0.036 (-0.026, 0.099) | 0.252 |
| | R | 3556 | 0.226 (0.127, 0.324) | <0.001 | 0.031 (-0.160, 0.223) | 0.748 | 0.900 (0.832, 0.969) | <0.001 | 0.078 (-0.022, 0.178) | 0.127 | 0.165 (0.097, 0.233) | <0.001 | 0.062 (-0.009, 0.133) | 0.088 |
| **Hippocampus** | L | 3586 | -0.080 (-0.189, 0.030) | 0.154 | 0.018 (-0.171, 0.206) | 0.854 | 0.891 (0.825, 0.958) | <0.001 | 0.041 (-0.075, 0.157) | 0.486 | -0.027 (-0.089, 0.034) | 0.382 | -0.046 (-0.109, 0.016) | 0.148 |
| | R | 3585 | -0.042 (-0.148, 0.065) | 0.442 | 0.014 (-0.175, 0.203) | 0.888 | 0.890 (0.823, 0.957) | <0.001 | 0.037 (-0.072, 0.146) | 0.502 | -0.047 (-0.108, 0.014) | 0.132 | -0.009 (-0.070, 0.051) | 0.767 |
| **Amygdala** | L | 3589 | 0.010 (-0.096, 0.116) | 0.852 | 0.015 (-0.174, 0.203) | 0.879 | 0.889 (0.822, 0.956) | <0.001 | 0.096 (-0.010, 0.202) | 0.077 | -0.029 (-0.091, 0.034) | 0.370 | 0.010 (-0.052, 0.073) | 0.749 |
| | R | 3613 | -0.034 (-0.140, 0.071) | 0.522 | 0.015 (-0.174, 0.204) | 0.875 | 0.893 (0.826, 0.960) | <0.001 | 0.020 (-0.085, 0.125) | 0.709 | -0.039 (-0.102, 0.023) | 0.213 | -0.033 (-0.093, 0.027) | 0.286 |
| **Accumbens** | L | 3549 | 0.021 (-0.086, 0.128) | 0.702 | 0.040 (-0.152, 0.232) | 0.686 | 0.890 (0.822, 0.957) | <0.001 | 0.011 (-0.096, 0.118) | 0.844 | 0.028 (-0.039, 0.095) | 0.408 | -0.022 (-0.090, 0.045) | 0.517 |
| | R | 3548 | 0.079 (-0.036, 0.194) | 0.177 | 0.031 (-0.160, 0.221) | 0.754 | 0.889 (0.822, 0.956) | <0.001 | 0.042 (-0.078, 0.162) | 0.490 | 0.032 (-0.034, 0.097) | 0.341 | -0.012 (-0.082, 0.057) | 0.733 |
| **Ventral DC** | L | 3613 | -0.170 (-0.281, -0.060) | 0.003 | 0.017 (-0.174, 0.207) | 0.864 | 0.915 (0.847, 0.984) | <0.001 | 0.048 (-0.065, 0.161) | 0.405 | -0.061 (-0.119, -0.002) | 0.044 | 0.011 (-0.045, 0.067) | 0.707 |
| | R | 3618 | -0.150 (-0.261, -0.040) | 0.008 | 0.019 (-0.170, 0.208) | 0.841 | 0.914 (0.845, 0.983) | <0.001 | 0.033 (-0.078, 0.145) | 0.557 | -0.081 (-0.140, -0.021) | 0.008 | 0.012 (-0.043, 0.068) | 0.664 |

**Note:** Estimated regression coefficients (β), 95% confidence intervals (CI) and *p*-values from the linear mixed-effects models evaluating the moderating effects of NO2 in the relationship between brain microstructure and Body Mass Index. Models were controlled for age, education, race, ethnicity, puberty, co-exposure to other air pollutants, head motion during the MRI acquisition and urbanicity. Random intercepts were modelled from MRI scanner device and subject ID. Although all two-way interactions were assessed, this table presents only the ones of primary interest due to space constraints. Abbreviations: ROI: region-of-interest; Ventral DC: ventral diencephalon; RSI: restricted spectrum imaging; RND: restricted normalized directional diffusion; H: hemisphere. *p-values < 0.05 of two-way and three-way interactions that achieved the Benjamini-Hochberg False Discovery Rate threshold correction.

**Supplementary Table 20.** Main effects and interactions for the O3 pollutant for males in RND.

| | | BMI ~ $ROI_{RND}$ * O3 * age + caregiver highest education + race + ethnicity + puberty + PM2.5 + NO2 + head motion + urbanicity address 1 + (1\|MRI device) + (1\| Subject ID) | | | | | | | | | | | | |
|---|---|---|---|---|---|---|---|---|---|---|---|---|---|---|
| | | | $ROI_{RND}$ | | O3 | | Age | | $ROI_{RND}$ X O3 | | $ROI_{RND}$ x Age | | $ROI_{RND}$ x O3 x Age | |
| **ROI** | **H** | **n** | **β (95%CI)** | **p-value** | **β (95%CI)** | **p-value** | **β (95%CI)** | **p-value** | **β (95%CI)** | **p-value** | **β (95%CI)** | **p-value** | **β (95%CI)** | **p-value** |
| **Thalamus** | L | 3631 | -0.122 (-0.225, -0.020) | 0.019 | 0.043 (-0.117, 0.203) | 0.599 | 0.914 (0.846, 0.983) | <0.001 | 0.035 (-0.068, 0.138) | 0.503 | -0.064 (-0.121, -0.007) | 0.028 | 0.005 (-0.053, 0.064) | 0.855 |
| | R | 3617 | -0.143 (-0.246, -0.039) | 0.007 | 0.047 (-0.112, 0.207) | 0.561 | 0.919 (0.850, 0.988) | <0.001 | -0.061 (-0.164, 0.042) | 0.245 | -0.054 (-0.112, 0.004) | 0.070 | -0.024 (-0.084, 0.035) | 0.419 |
| **Caudate** | L | 3555 | 0.073 (-0.043, 0.189) | 0.219 | 0.072 (-0.092, 0.235) | 0.390 | 0.890 (0.822, 0.958) | <0.001 | -0.138 (-0.250, -0.026) | 0.015 | 0.051 (-0.022, 0.124) | 0.168 | -0.044 (-0.118, 0.029) | 0.235 |
| | R | 3550 | -0.067 (-0.180, 0.046) | 0.246 | 0.041 (-0.123, 0.204) | 0.627 | 0.899 (0.831, 0.966) | <0.001 | -0.145 (-0.255, -0.036) | 0.010 | -0.017 (-0.087, 0.054) | 0.647 | -0.033 (-0.104, 0.038) | 0.360 |
| **Putamen** | L | 3567 | 0.300 (0.171, 0.430) | <0.001 | 0.092 (-0.073, 0.257) | 0.274 | 0.885 (0.817, 0.953) | <0.001 | -0.004 (-0.121, 0.114) | 0.953 | 0.119 (0.048, 0.191) | 0.001 | -0.023 (-0.093, 0.047) | 0.522 |
| | R | 3537 | 0.273 (0.152, 0.395) | <0.001 | 0.100 (-0.065, 0.265) | 0.235 | 0.875 (0.806, 0.943) | <0.001 | -0.024 (-0.139, 0.091) | 0.687 | 0.082 (0.010, 0.154) | 0.025 | -0.045 (-0.116, 0.027) | 0.221 |
| **Pallidum** | L | 3582 | 0.062 (-0.031, 0.156) | 0.192 | 0.051 (-0.112, 0.215) | 0.538 | 0.894 (0.825, 0.963) | <0.001 | -0.062 (-0.151, 0.027) | 0.172 | 0.043 (-0.021, 0.107) | 0.186 | -0.032 (-0.095, 0.031) | 0.325 |
| | R | 3556 | 0.232 (0.131, 0.332) | <0.001 | 0.075 (-0.089, 0.238) | 0.371 | 0.904 (0.835, 0.973) | <0.001 | -0.012 (-0.109, 0.086) | 0.815 | 0.156 (0.087, 0.225) | <0.001 | -0.020 (-0.090, 0.050) | 0.576 |
| **Hippocampus** | L | 3586 | -0.090 (-0.200, 0.021) | 0.111 | 0.040 (-0.122, 0.202) | 0.627 | 0.896 (0.828, 0.963) | <0.001 | -0.059 (-0.167, 0.049) | 0.288 | -0.029 (-0.092, 0.033) | 0.360 | 0.013 (-0.052, 0.077) | 0.695 |
| | R | 3585 | -0.059 (-0.166, 0.047) | 0.277 | 0.038 (-0.123, 0.198) | 0.646 | 0.885 (0.817, 0.952) | <0.001 | -0.161 (-0.265, -0.056) | 0.003 | -0.060 (-0.122, 0.002) | 0.060 | -0.071 (-0.134, -0.008) | 0.027 |
| **Amygdala** | L | 3589 | 0.026 (-0.082, 0.133) | 0.640 | 0.067 (-0.095, 0.228) | 0.420 | 0.879 (0.812, 0.947) | <0.001 | 0.015 (-0.091, 0.121) | 0.783 | -0.046 (-0.110, 0.018) | 0.159 | -0.050 (-0.116, 0.017) | 0.143 |
| | R | 3613 | -0.025 (-0.131, 0.082) | 0.650 | 0.062 (-0.100, 0.223) | 0.455 | 0.889 (0.822, 0.957) | <0.001 | 0.004 (-0.104, 0.112) | 0.937 | -0.048 (-0.112, 0.016) | 0.145 | -0.015 (-0.079, 0.049) | 0.646 |
| **Accumbens** | L | 3549 | 0.006 (-0.103, 0.114) | 0.917 | 0.071 (-0.092, 0.235) | 0.394 | 0.900 (0.832, 0.968) | <0.001 | -0.134 (-0.239, -0.030) | 0.012 | 0.035 (-0.033, 0.103) | 0.317 | -0.020 (-0.088, 0.048) | 0.564 |
| | R | 3548 | 0.072 (-0.045, 0.189) | 0.231 | 0.089 (-0.074, 0.251) | 0.285 | 0.891 (0.824, 0.959) | <0.001 | -0.069 (-0.182, 0.045) | 0.236 | 0.026 (-0.041, 0.094) | 0.443 | -0.032 (-0.099, 0.036) | 0.357 |
| **Ventral DC** | L | 3613 | -0.167 (-0.277, -0.056) | 0.003 | 0.017 (-0.144, 0.179) | 0.832 | 0.915 (0.848, 0.983) | <0.001 | 0.040 (-0.069, 0.150) | 0.471 | -0.068 (-0.126, -0.011) | 0.021 | -0.019 (-0.076, 0.038) | 0.506 |
| | R | 3618 | -0.148 (-0.257, -0.038) | 0.008 | 0.027 (-0.133, 0.187) | 0.744 | 0.922 (0.854, 0.989) | <0.001 | 0.109 (-0.001, 0.220) | 0.052 | -0.094 (-0.151, -0.036) | 0.002 | 0.025 (-0.033, 0.084) | 0.396 |

**Note:** Estimated regression coefficients (β), 95% confidence intervals (CI) and *p*-values from the linear mixed-effects models evaluating the moderating effects of O3 in the relationship between brain microstructure and Body Mass Index. Models were controlled for age, education, race, ethnicity, puberty, co-exposure to other air pollutants, head motion during the MRI acquisition and urbanicity. Random intercepts were modelled from MRI scanner device and subject ID. Although all two-way interactions were assessed, this table presents only the ones of primary interest due to space constraints. Abbreviations: ROI: region-of-interest; Ventral DC: ventral diencephalon; RSI: restricted spectrum imaging; RND: restricted normalized directional diffusion; H: hemisphere. *p-values < 0.05 of two-way and three-way interactions that achieved the Benjamini-Hochberg False Discovery Rate threshold correction.

**Supplementary Table 21.** Main effects and interactions for the Oxwt measure in males in RND.

| | | BMI ~ $ROI_{RND}$ * Oxwt * age + caregiver highest education + race + ethnicity + puberty + PM2.5 + head motion + urbanicity address 1 + (1\|MRI device) + (1\| Subject ID) | | | | | | | | | | | | |
|---|---|---|---|---|---|---|---|---|---|---|---|---|---|---|
| | | | $ROI_{RND}$ | | Oxwt | | Age | | $ROI_{RND}$ X Oxwt | | $ROI_{RND}$ x Age | | $ROI_{RND}$ x Oxwt x Age | |
| **ROI** | **H** | **n** | **β (95%CI)** | **p-value** | **β (95%CI)** | **p-value** | **β (95%CI)** | **p-value** | **β (95%CI)** | **p-value** | **β (95%CI)** | **p-value** | **β (95%CI)** | **p-value** |
| **Thalamus** | L | 3631 | -0.123 (-0.225, -0.021) | 0.019 | 0.043 (-0.124, 0.209) | 0.618 | 0.913 (0.845, 0.981) | <0.001 | 0.053 (-0.049, 0.155) | 0.309 | -0.061 (-0.118, -0.004) | 0.036 | -0.008 (-0.065, 0.050) | 0.799 |
| | R | 3617 | -0.139 (-0.242, -0.035) | 0.009 | 0.050 (-0.116, 0.216) | 0.558 | 0.918 (0.850, 0.987) | <0.001 | -0.007 (-0.109, 0.096) | 0.899 | -0.052 (-0.110, 0.006) | 0.079 | -0.026 (-0.086, 0.033) | 0.388 |
| **Caudate** | L | 3555 | 0.089 (-0.027, 0.204) | 0.133 | 0.084 (-0.087, 0.256) | 0.336 | 0.887 (0.819, 0.955) | <0.001 | -0.083 (-0.193, 0.028) | 0.142 | 0.046 (-0.026, 0.119) | 0.212 | -0.039 (-0.112, 0.033) | 0.291 |
| | R | 3550 | -0.057 (-0.170, 0.056) | 0.321 | 0.043 (-0.129, 0.215) | 0.624 | 0.894 (0.827, 0.962) | <0.001 | -0.119 (-0.228, -0.010) | 0.032 | -0.025 (-0.096, 0.045) | 0.485 | -0.048 (-0.120, 0.023) | 0.186 |
| **Putamen** | L | 3567 | 0.302 (0.173, 0.431) | <0.001 | 0.116 (-0.055, 0.288) | 0.184 | 0.885 (0.817, 0.953) | <0.001 | 0.055 (-0.060, 0.169) | 0.348 | 0.121 (0.049, 0.193) | 0.001 | 0.008 (-0.062, 0.078) | 0.823 |
| | R | 3537 | 0.270 (0.149, 0.391) | <0.001 | 0.117 (-0.056, 0.289) | 0.186 | 0.876 (0.808, 0.944) | <0.001 | 0.037 (-0.077, 0.152) | 0.526 | 0.087 (0.015, 0.159) | 0.018 | -0.001 (-0.072, 0.070) | 0.982 |
| **Pallidum** | L | 3582 | 0.072 (-0.022, 0.165) | 0.132 | 0.063 (-0.107, 0.234) | 0.467 | 0.891 (0.822, 0.960) | <0.001 | -0.006 (-0.094, 0.082) | 0.887 | 0.045 (-0.019, 0.109) | 0.171 | 0.001 (-0.062, 0.064) | 0.982 |
| | R | 3556 | 0.237 (0.137, 0.337) | <0.001 | 0.082 (-0.089, 0.253) | 0.349 | 0.904 (0.835, 0.972) | <0.001 | 0.027 (-0.068, 0.122) | 0.579 | 0.161 (0.092, 0.229) | <0.001 | 0.020 (-0.050, 0.090) | 0.578 |
| **Hippocampus** | L | 3586 | -0.084 (-0.195, 0.027) | 0.140 | 0.047 (-0.122, 0.216) | 0.584 | 0.889 (0.822, 0.956) | <0.001 | -0.036 (-0.146, 0.074) | 0.523 | -0.032 (-0.094, 0.031) | 0.317 | -0.014 (-0.079, 0.051) | 0.667 |
| | R | 3585 | -0.055 (-0.162, 0.052) | 0.318 | 0.046 (-0.122, 0.214) | 0.592 | 0.885 (0.818, 0.952) | <0.001 | -0.118 (-0.222, -0.014) | 0.026 | -0.056 (-0.118, 0.006) | 0.078 | -0.060 (-0.122, 0.001) | 0.055 |
| **Amygdala** | L | 3589 | 0.023 (-0.084, 0.130) | 0.669 | 0.062 (-0.107, 0.231) | 0.474 | 0.882 (0.814, 0.949) | <0.001 | 0.057 (-0.048, 0.161) | 0.286 | -0.043 (-0.106, 0.021) | 0.190 | -0.031 (-0.097, 0.035) | 0.357 |
| | R | 3613 | -0.027 (-0.134, 0.079) | 0.615 | 0.061 (-0.108, 0.230) | 0.480 | 0.888 (0.821, 0.955) | <0.001 | 0.006 (-0.099, 0.111) | 0.908 | -0.049 (-0.112, 0.015) | 0.132 | -0.031 (-0.093, 0.031) | 0.328 |
| **Accumbens** | L | 3549 | 0.009 (-0.099, 0.118) | 0.867 | 0.083 (-0.088, 0.254) | 0.343 | 0.897 (0.829, 0.965) | <0.001 | -0.113 (-0.217, -0.008) | 0.035 | 0.024 (-0.044, 0.092) | 0.493 | -0.035 (-0.105, 0.034) | 0.319 |
| | R | 3548 | 0.082 (-0.036, 0.199) | 0.173 | 0.091 (-0.080, 0.261) | 0.297 | 0.889 (0.822, 0.957) | <0.001 | -0.041 (-0.155, 0.072) | 0.474 | 0.025 (-0.043, 0.092) | 0.474 | -0.033 (-0.098, 0.033) | 0.328 |
| **Ventral DC** | L | 3613 | -0.166 (-0.276, -0.056) | 0.003 | 0.019 (-0.149, 0.187) | 0.826 | 0.918 (0.850, 0.985) | <0.001 | 0.064 (-0.050, 0.177) | 0.271 | -0.063 (-0.120, -0.006) | 0.031 | -0.006 (-0.066, 0.053) | 0.837 |
| | R | 3618 | -0.146 (-0.255, -0.037) | 0.009 | 0.022 (-0.145, 0.188) | 0.798 | 0.917 (0.849, 0.985) | <0.001 | 0.121 (0.007, 0.234) | 0.037 | -0.081 (-0.138, -0.024) | 0.006 | 0.034 (-0.026, 0.094) | 0.263 |

**Note:** Estimated regression coefficients (β), 95% confidence intervals (CI) and *p*-values from the linear mixed-effects models evaluating the moderating effects of Oxwt in the relationship between brain microstructure and Body Mass Index. Models were controlled for age, education, race, ethnicity, puberty, co-exposure to other air pollutants, head motion during the MRI acquisition and urbanicity. Random intercepts were modelled from MRI scanner device and subject ID. Although all two-way interactions were assessed, this table presents only the ones of primary interest due to space constraints. Abbreviations: ROI: region-of-interest; Ventral DC: ventral diencephalon; RSI: restricted spectrum imaging; RND: restricted normalized directional diffusion; H: hemisphere. *p-values < 0.05 of two-way and three-way interactions that achieved the Benjamini-Hochberg False Discovery Rate threshold correction.

**Supplementary Table 22.** Main effects and interactions for the PM2.5 pollutant for females in RND.

| | | BMI ~ $ROI_{RND}$ * PM2.5 * age + caregiver highest education + race + ethnicity + puberty + NO2 + O3 + head motion + urbanicity address 1 + (1\|MRI device) + (1\| Subject ID) | | | | | | | | | | | | |
|---|---|---|---|---|---|---|---|---|---|---|---|---|---|---|
| | | | $ROI_{RND}$ | | PM2.5 | | Age | | $ROI_{RND}$ X PM2.5 | | $ROI_{RND}$ x Age | | $ROI_{RND}$ x PM2.5 x Age | |
| **ROI** | **H** | **n** | **β (95%CI)** | **p-value** | **β (95%CI)** | **p-value** | **β (95%CI)** | **p-value** | **β (95%CI)** | **p-value** | **β (95%CI)** | **p-value** | **β (95%CI)** | **p-value** |
| **Thalamus** | L | 3586 | -0.138 (-0.246, -0.030) | 0.012 | 0.098 (-0.073, 0.270) | 0.264 | 0.793 (0.698, 0.889) | <0.001 | -0.090 (-0.195, 0.016) | 0.095 | 0.024 (-0.039, 0.087) | 0.452 | -0.018 (-0.077, 0.042) | 0.564 |
| | R | 3581 | -0.021 (-0.129, 0.088) | 0.707 | 0.096 (-0.073, 0.266) | 0.267 | 0.773 (0.677, 0.868) | <0.001 | -0.065 (-0.171, 0.041) | 0.229 | 0.064 (0.001, 0.126) | 0.046 | 0.035 (-0.024, 0.094) | 0.243 |
| **Caudate** | L | 3542 | 0.197 (0.083, 0.312) | 0.001 | 0.070 (-0.103, 0.242) | 0.429 | 0.792 (0.697, 0.888) | <0.001 | -0.075 (-0.202, 0.053) | 0.252 | 0.144 (0.065, 0.224) | <0.001 | -0.032 (-0.122, 0.057) | 0.480 |
| | R | 3522 | 0.075 (-0.044, 0.193) | 0.217 | 0.087 (-0.085, 0.259) | 0.321 | 0.774 (0.678, 0.871) | <0.001 | -0.079 (-0.208, 0.050) | 0.229 | 0.094 (0.013, 0.174) | 0.023 | 0.037 (-0.049, 0.123) | 0.403 |
| **Putamen** | L | 3535 | 0.207 (0.075, 0.339) | 0.002 | 0.080 (-0.094, 0.253) | 0.370 | 0.763 (0.667, 0.858) | <0.001 | -0.009 (-0.153, 0.134) | 0.899 | 0.123 (0.044, 0.201) | 0.002 | 0.041 (-0.050, 0.132) | 0.378 |
| | R | 3517 | 0.192 (0.063, 0.321) | 0.004 | 0.076 (-0.097, 0.249) | 0.392 | 0.754 (0.657, 0.850) | <0.001 | -0.050 (-0.196, 0.095) | 0.497 | 0.148 (0.067, 0.230) | <0.001 | 0.017 (-0.078, 0.112) | 0.731 |
| **Pallidum** | L | 3540 | 0.133 (0.035, 0.230) | 0.008 | 0.104 (-0.062, 0.270) | 0.223 | 0.749 (0.654, 0.844) | <0.001 | 0.090 (-0.011, 0.192) | 0.081 | 0.159 (0.090, 0.227) | <0.001 | 0.063 (-0.0004, 0.126) | 0.052 |
| | R | 3525 | 0.189 (0.087, 0.292) | <0.001 | 0.077 (-0.095, 0.249) | 0.384 | 0.751 (0.655, 0.847) | <0.001 | -0.073 (-0.181, 0.035) | 0.185 | 0.148 (0.070, 0.225) | <0.001 | -0.019 (-0.093, 0.056) | 0.627 |
| **Hippocampus** | L | 3552 | 0.023 (-0.094, 0.140) | 0.704 | 0.081 (-0.094, 0.256) | 0.367 | 0.773 (0.677, 0.868) | <0.001 | -0.103 (-0.225, 0.019) | 0.097 | 0.059 (-0.013, 0.131) | 0.106 | 0.022 (-0.051, 0.095) | 0.558 |
| | R | 3551 | -0.030 (-0.144, 0.083) | 0.601 | 0.083 (-0.090, 0.256) | 0.349 | 0.766 (0.671, 0.862) | <0.001 | -0.124 (-0.248, -0.0004) | 0.050 | 0.007 (-0.064, 0.078) | 0.845 | -0.027 (-0.100, 0.047) | 0.472 |
| **Amygdala** | L | 3540 | -0.042 (-0.147, 0.063) | 0.433 | 0.095 (-0.077, 0.267) | 0.283 | 0.766 (0.671, 0.861) | <0.001 | -0.146 (-0.259, -0.034) | 0.011 | 0.032 (-0.036, 0.101) | 0.352 | -0.055 (-0.130, 0.019) | 0.144 |
| | R | 3550 | -0.080 (-0.191, 0.032) | 0.161 | 0.122 (-0.047, 0.291) | 0.160 | 0.773 (0.679, 0.868) | <0.001 | -0.044 (-0.162, 0.075) | 0.471 | 0.049 (-0.020, 0.119) | 0.163 | 0.025 (-0.051, 0.101) | 0.518 |
| **Accumbens** | L | 3532 | 0.058 (-0.058, 0.174) | 0.326 | 0.106 (-0.063, 0.275) | 0.221 | 0.783 (0.688, 0.879) | <0.001 | 0.023 (-0.101, 0.147) | 0.720 | 0.115 (0.038, 0.192) | 0.004 | -0.010 (-0.092, 0.073) | 0.821 |
| | R | 3524 | 0.162 (0.041, 0.282) | 0.009 | 0.074 (-0.097, 0.246) | 0.396 | 0.785 (0.689, 0.882) | <0.001 | -0.083 (-0.212, 0.046) | 0.209 | 0.141 (0.063, 0.218) | <0.001 | -0.057 (-0.140, 0.026) | 0.178 |
| **Ventral DC** | L | 3557 | -0.220 (-0.332, -0.108) | <0.001 | 0.068 (-0.110, 0.247) | 0.453 | 0.799 (0.704, 0.894) | <0.001 | -0.101 (-0.202, -0.0003) | 0.050 | -0.001 (-0.064, 0.062) | 0.968 | -0.024 (-0.077, 0.029) | 0.380 |
| | R | 3546 | -0.222 (-0.335, -0.109) | <0.001 | 0.077 (-0.099, 0.253) | 0.395 | 0.810 (0.714, 0.905) | <0.001 | -0.070 (-0.172, 0.032) | 0.180 | -0.022 (-0.085, 0.041) | 0.495 | 0.002 (-0.051, 0.055) | 0.936 |

**Note:** Estimated regression coefficients (β), 95% confidence intervals (CI) and *p*-values from the linear mixed-effects models evaluating the moderating effects of PM2.5 in the relationship between brain microstructure and Body Mass Index. Models were controlled for age, education, race, ethnicity, puberty, co-exposure to other air pollutants, head motion during the MRI acquisition and urbanicity. Random intercepts were modelled from MRI scanner device and subject ID. Although all two-way interactions were assessed, this table presents only the ones of primary interest due to space constraints. Abbreviations: ROI: region-of-interest; Ventral DC: ventral diencephalon; RSI: restricted spectrum imaging; RND: restricted normalized directional diffusion; H: hemisphere. *p-values < 0.05 of two-way and three-way interactions that achieved the Benjamini-Hochberg False Discovery Rate threshold correction.

**Supplementary Table 23.** Main effects and interactions for the NO2 pollutant for females in RND.

| | | BMI ~ $ROI_{RND}$ * NO2 * age + caregiver highest education + race + ethnicity + puberty + PM2.5 + O3 + head motion + urbanicity address 1 + (1\|MRI device) + (1\| Subject ID) | | | | | | | | | | | | |
|---|---|---|---|---|---|---|---|---|---|---|---|---|---|---|
| | | | $ROI_{RND}$ | | NO2 | | Age | | $ROI_{RND}$ X NO2 | | $ROI_{RND}$ x Age | | $ROI_{RND}$ x NO2 x Age | |
| **ROI** | **H** | **n** | **β (95%CI)** | **p-value** | **β (95%CI)** | **p-value** | **β (95%CI)** | **p-value** | **β (95%CI)** | **p-value** | **β (95%CI)** | **p-value** | **β (95%CI)** | **p-value** |
| **Thalamus** | L | 3586 | -0.149 (-0.256, -0.042) | 0.006 | 0.007 (-0.169, 0.183) | 0.940 | 0.805 (0.710, 0.899) | <0.001 | -0.036 (-0.147, 0.076) | 0.532 | 0.022 (-0.041, 0.085) | 0.488 | -0.043 (-0.107, 0.021) | 0.192 |
| | R | 3581 | -0.029 (-0.138, 0.079) | 0.594 | 0.008 (-0.167, 0.183) | 0.926 | 0.789 (0.693, 0.884) | <0.001 | -0.049 (-0.164, 0.066) | 0.402 | 0.059 (-0.004, 0.122) | 0.067 | -0.024 (-0.087, 0.039) | 0.458 |
| **Caudate** | L | 3542 | 0.179 (0.069, 0.290) | 0.002 | 0.033 (-0.145, 0.211) | 0.717 | 0.795 (0.701, 0.889) | <0.001 | 0.053 (-0.060, 0.166) | 0.355 | 0.153 (0.076, 0.230) | <0.001 | -0.047 (-0.130, 0.035) | 0.262 |
| | R | 3522 | 0.042 (-0.074, 0.157) | 0.479 | 0.005 (-0.174, 0.183) | 0.958 | 0.793 (0.698, 0.888) | <0.001 | -0.022 (-0.142, 0.099) | 0.726 | 0.114 (0.037, 0.191) | 0.004 | -0.072 (-0.155, 0.011) | 0.091 |
| **Putamen** | L | 3535 | 0.200 (0.074, 0.326) | 0.002 | 0.031 (-0.148, 0.211) | 0.732 | 0.774 (0.680, 0.868) | <0.001 | 0.093 (-0.046, 0.233) | 0.188 | 0.135 (0.059, 0.211) | 0.001 | -0.024 (-0.108, 0.060) | 0.577 |
| | R | 3517 | 0.179 (0.055, 0.302) | 0.005 | 0.027 (-0.152, 0.207) | 0.766 | 0.766 (0.672, 0.861) | <0.001 | 0.042 (-0.088, 0.173) | 0.525 | 0.158 (0.081, 0.236) | <0.001 | -0.092 (-0.178, -0.006) | 0.037 |
| **Pallidum** | L | 3540 | 0.132 (0.036, 0.229) | 0.007 | -0.002 (-0.175, 0.172) | 0.984 | 0.754 (0.659, 0.849) | <0.001 | 0.089 (-0.013, 0.190) | 0.088 | 0.158 (0.090, 0.226) | <0.001 | -0.005 (-0.071, 0.062) | 0.891 |
| | R | 3525 | 0.164 (0.066, 0.263) | 0.001 | -0.002 (-0.178, 0.175) | 0.984 | 0.759 (0.664, 0.854) | <0.001 | 0.071 (-0.032, 0.174) | 0.179 | 0.151 (0.076, 0.227) | <0.001 | -0.012 (-0.089, 0.066) | 0.769 |
| **Hippocampus** | L | 3552 | -0.020 (-0.131, 0.091) | 0.724 | -0.002 (-0.179, 0.174) | 0.979 | 0.787 (0.693, 0.881) | <0.001 | 0.027 (-0.095, 0.150) | 0.665 | 0.065 (-0.005, 0.136) | 0.069 | -0.038 (-0.113, 0.036) | 0.311 |
| | R | 3551 | -0.067 (-0.176, 0.042) | 0.228 | 0.012 (-0.164, 0.188) | 0.894 | 0.779 (0.685, 0.873) | <0.001 | 0.001 (-0.113, 0.115) | 0.988 | 0.012 (-0.057, 0.081) | 0.737 | -0.011 (-0.081, 0.059) | 0.763 |
| **Amygdala** | L | 3540 | -0.059 (-0.162, 0.044) | 0.263 | 0.001 (-0.177, 0.179) | 0.992 | 0.775 (0.681, 0.870) | <0.001 | 0.047 (-0.060, 0.154) | 0.389 | 0.033 (-0.034, 0.099) | 0.333 | -0.031 (-0.100, 0.038) | 0.379 |
| | R | 3550 | -0.087 (-0.197, 0.022) | 0.120 | 0.005 (-0.171, 0.181) | 0.953 | 0.782 (0.688, 0.876) | <0.001 | -0.010 (-0.125, 0.105) | 0.859 | 0.060 (-0.007, 0.128) | 0.079 | -0.067 (-0.137, 0.003) | 0.060 |
| **Accumbens** | L | 3532 | 0.059 (-0.052, 0.171) | 0.299 | 0.015 (-0.161, 0.191) | 0.865 | 0.783 (0.689, 0.877) | <0.001 | 0.050 (-0.076, 0.175) | 0.439 | 0.113 (0.040, 0.186) | 0.003 | -0.015 (-0.097, 0.067) | 0.717 |
| | R | 3524 | 0.152 (0.035, 0.269) | 0.011 | 0.027 (-0.150, 0.205) | 0.764 | 0.782 (0.688, 0.876) | <0.001 | 0.053 (-0.075, 0.180) | 0.416 | 0.137 (0.063, 0.211) | <0.001 | -0.031 (-0.112, 0.050) | 0.452 |
| **Ventral DC** | L | 3557 | -0.233 (-0.344, -0.123) | <0.001 | 0.021 (-0.157, 0.200) | 0.814 | 0.808 (0.713, 0.903) | <0.001 | -0.019 (-0.131, 0.093) | 0.738 | -0.010 (-0.074, 0.054) | 0.762 | -0.015 (-0.078, 0.048) | 0.639 |
| | R | 3546 | -0.222 (-0.335, -0.109) | <0.001 | 0.034 (-0.144, 0.213) | 0.709 | 0.820 (0.725, 0.915) | <0.001 | -0.009 (-0.121, 0.104) | 0.878 | -0.033 (-0.097, 0.031) | 0.318 | -0.015 (-0.077, 0.047) | 0.626 |

**Note:** Estimated regression coefficients (β), 95% confidence intervals (CI) and *p*-values from the linear mixed-effects models evaluating the moderating effects of NO2 in the relationship between brain microstructure and Body Mass Index. Models were controlled for age, education, race, ethnicity, puberty, co-exposure to other air pollutants, head motion during the MRI acquisition and urbanicity. Random intercepts were modelled from MRI scanner device and subject ID. Although all two-way interactions were assessed, this table presents only the ones of primary interest due to space constraints. Abbreviations: ROI: region-of-interest; Ventral DC: ventral diencephalon; RSI: restricted spectrum imaging; RND: restricted normalized directional diffusion; H: hemisphere. *p-values < 0.05 of two-way and three-way interactions that achieved the Benjamini-Hochberg False Discovery Rate threshold correction.

**Supplementary Table 24.** Main effects and interactions for the O3 pollutant for females in RND.

| | | BMI ~ $ROI_{RND}$ * O3 * age + caregiver highest education + race + ethnicity + puberty + PM2.5 + NO2 + head motion + urbanicity address 1 + (1\|MRI device) + (1\| Subject ID) | | | | | | | | | | | | |
|---|---|---|---|---|---|---|---|---|---|---|---|---|---|---|
| | | | $ROI_{RND}$ | | O3 | | Age | | $ROI_{RND}$ X O3 | | $ROI_{RND}$ x Age | | $ROI_{RND}$ x O3 x Age | |
| **ROI** | **H** | **n** | **β (95%CI)** | **p-value** | **β (95%CI)** | **p-value** | **β (95%CI)** | **p-value** | **β (95%CI)** | **p-value** | **β (95%CI)** | **p-value** | **β (95%CI)** | **p-value** |
| **Thalamus** | L | 3586 | -0.159 (-0.266, -0.052) | 0.004 | 0.061 (-0.095, 0.217) | 0.445 | 0.797 (0.702, 0.892) | <0.001 | 0.023 (-0.084, 0.131) | 0.671 | 0.033 (-0.030, 0.096) | 0.310 | -0.021 (-0.084, 0.041) | 0.504 |
| | R | 3581 | -0.026 (-0.135, 0.082) | 0.632 | 0.077 (-0.079, 0.232) | 0.334 | 0.778 (0.683, 0.874) | <0.001 | -0.042 (-0.154, 0.069) | 0.455 | 0.070 (0.008, 0.133) | 0.027 | -0.028 (-0.090, 0.035) | 0.388 |
| **Caudate** | L | 3542 | 0.223 (0.108, 0.337) | <0.001 | 0.111 (-0.048, 0.271) | 0.173 | 0.789 (0.696, 0.883) | <0.001 | 0.198 (0.084, 0.311) | 0.001 | 0.152 (0.073, 0.230) | <0.001 | 0.026 (-0.055, 0.106) | 0.531 |
| | R | 3522 | 0.056 (-0.061, 0.174) | 0.348 | 0.086 (-0.072, 0.244) | 0.287 | 0.784 (0.688, 0.879) | <0.001 | 0.041 (-0.073, 0.154) | 0.485 | 0.116 (0.037, 0.194) | 0.004 | -0.026 (-0.105, 0.054) | 0.530 |
| **Putamen** | L | 3535 | 0.208 (0.079, 0.338) | 0.002 | 0.105 (-0.053, 0.264) | 0.194 | 0.765 (0.670, 0.859) | <0.001 | 0.076 (-0.043, 0.195) | 0.211 | 0.139 (0.063, 0.216) | <0.001 | -0.011 (-0.088, 0.066) | 0.779 |
| | R | 3517 | 0.192 (0.065, 0.319) | 0.003 | 0.102 (-0.057, 0.262) | 0.209 | 0.751 (0.656, 0.846) | <0.001 | 0.087 (-0.030, 0.204) | 0.146 | 0.158 (0.080, 0.237) | <0.001 | -0.029 (-0.108, 0.050) | 0.474 |
| **Pallidum** | L | 3540 | 0.133 (0.036, 0.230) | 0.008 | 0.104 (-0.051, 0.259) | 0.189 | 0.750 (0.655, 0.845) | <0.001 | 0.021 (-0.073, 0.115) | 0.663 | 0.167 (0.099, 0.235) | <0.001 | -0.034 (-0.102, 0.035) | 0.337 |
| | R | 3525 | 0.193 (0.092, 0.293) | <0.001 | 0.085 (-0.074, 0.243) | 0.295 | 0.740 (0.645, 0.835) | <0.001 | 0.106 (0.013, 0.199) | 0.025 | 0.151 (0.076, 0.227) | <0.001 | -0.061 (-0.137, 0.014) | 0.110 |
| **Hippocampus** | L | 3552 | -0.015 (-0.127, 0.097) | 0.794 | 0.090 (-0.068, 0.247) | 0.264 | 0.780 (0.685, 0.876) | <0.001 | 0.036 (-0.073, 0.145) | 0.516 | 0.075 (0.003, 0.147) | 0.041 | -0.015 (-0.089, 0.058) | 0.685 |
| | R | 3551 | -0.047 (-0.157, 0.063) | 0.405 | 0.087 (-0.071, 0.245) | 0.279 | 0.770 (0.674, 0.865) | <0.001 | 0.104 (-0.002, 0.210) | 0.056 | 0.013 (-0.058, 0.083) | 0.719 | -0.015 (-0.085, 0.056) | 0.682 |
| **Amygdala** | L | 3540 | -0.075 (-0.179, 0.029) | 0.157 | 0.057 (-0.099, 0.214) | 0.472 | 0.772 (0.677, 0.868) | <0.001 | -0.017 (-0.118, 0.084) | 0.743 | 0.042 (-0.026, 0.110) | 0.223 | 0.006 (-0.065, 0.076) | 0.878 |
| | R | 3550 | -0.097 (-0.207, 0.013) | 0.086 | 0.061 (-0.095, 0.216) | 0.444 | 0.776 (0.682, 0.871) | <0.001 | -0.058 (-0.166, 0.050) | 0.291 | 0.064 (-0.005, 0.134) | 0.069 | -0.033 (-0.104, 0.037) | 0.354 |
| **Accumbens** | L | 3532 | 0.068 (-0.047, 0.183) | 0.245 | 0.083 (-0.073, 0.240) | 0.298 | 0.773 (0.678, 0.868) | <0.001 | 0.020 (-0.093, 0.134) | 0.724 | 0.114 (0.039, 0.190) | 0.003 | -0.039 (-0.116, 0.038) | 0.320 |
| | R | 3524 | 0.161 (0.041, 0.281) | 0.009 | 0.095 (-0.063, 0.253) | 0.240 | 0.771 (0.676, 0.867) | <0.001 | 0.039 (-0.083, 0.162) | 0.531 | 0.142 (0.066, 0.218) | <0.001 | -0.028 (-0.108, 0.053) | 0.500 |
| **Ventral DC** | L | 3557 | -0.240 (-0.351, -0.129) | <0.001 | 0.054 (-0.104, 0.213) | 0.500 | 0.804 (0.709, 0.898) | <0.001 | 0.038 (-0.070, 0.146) | 0.493 | -0.001 (-0.064, 0.063) | 0.985 | 0.006 (-0.058, 0.071) | 0.844 |
| | R | 3546 | -0.228 (-0.341, -0.116) | <0.001 | 0.051 (-0.107, 0.209) | 0.529 | 0.817 (0.721, 0.912) | <0.001 | 0.005 (-0.103, 0.112) | 0.933 | -0.022 (-0.085, 0.042) | 0.506 | 0.012 (-0.052, 0.076) | 0.714 |

**Note:** Estimated regression coefficients (β), 95% confidence intervals (CI) and *p*-values from the linear mixed-effects models evaluating the moderating effects of O3 in the relationship between brain microstructure and Body Mass Index. Models were controlled for age, education, race, ethnicity, puberty, co-exposure to other air pollutants, head motion during the MRI acquisition and urbanicity. Random intercepts were modelled from MRI scanner device and subject ID. Although all two-way interactions were assessed, this table presents only the ones of primary interest due to space constraints. Abbreviations: ROI: region-of-interest; Ventral DC: ventral diencephalon; RSI: restricted spectrum imaging; RND: restricted normalized directional diffusion; H: hemisphere. *p-values < 0.05 of two-way and three-way interactions that achieved the Benjamini-Hochberg False Discovery Rate threshold correction.

**Supplementary Table 25.** Main effects and interactions for the Oxwt measure in females in RND.

| | | **BMI ~ ROI$_{RND}$ * Oxwt * age + caregiver highest education + race + ethnicity + puberty + PM2.5 + head motion + urbanicity address 1 + (1\|MRI device) + (1\| Subject ID)** | | | | | | | | | | | | |
|---|---|---|---|---|---|---|---|---|---|---|---|---|---|---|
| | | | **ROI$_{RND}$** | | **Oxwt** | | **Age** | | **ROI$_{RND}$ X Oxwt** | | **ROI$_{RND}$ x Age** | | **ROI$_{RND}$ x Oxwt x Age** | |
| **ROI** | **H** | **n** | **β (95%CI)** | **p-value** | **β (95%CI)** | **p-value** | **β (95%CI)** | **p-value** | **β (95%CI)** | **p-value** | **β (95%CI)** | **p-value** | **β (95%CI)** | **p-value** |
| **Thalamus** | L | 3586 | -0.159 (-0.266, -0.053) | 0.003 | 0.059 (-0.102, 0.220) | 0.474 | 0.797 (0.703, 0.892) | <0.001 | 0.004 (-0.103, 0.110) | 0.947 | 0.031 (-0.031, 0.094) | 0.326 | -0.043 (-0.105, 0.019) | 0.175 |
| | R | 3581 | -0.028 (-0.137, 0.080) | 0.608 | 0.074 (-0.088, 0.235) | 0.372 | 0.781 (0.685, 0.876) | <0.001 | -0.063 (-0.177, 0.051) | 0.275 | 0.068 (0.006, 0.130) | 0.033 | -0.038 (-0.102, 0.025) | 0.238 |
| **Caudate** | L | 3542 | 0.217 (0.103, 0.332) | <0.001 | 0.108 (-0.058, 0.273) | 0.202 | 0.787 (0.693, 0.881) | <0.001 | 0.179 (0.071, 0.286) | 0.001 | 0.151 (0.073, 0.229) | <0.001 | -0.002 (-0.078, 0.074) | 0.950 |
| | R | 3522 | 0.053 (-0.065, 0.171) | 0.380 | 0.079 (-0.085, 0.244) | 0.346 | 0.782 (0.687, 0.877) | <0.001 | 0.020 (-0.093, 0.133) | 0.731 | 0.114 (0.036, 0.192) | 0.004 | -0.057 (-0.136, 0.022) | 0.155 |
| **Putamen** | L | 3535 | 0.219 (0.088, 0.350) | 0.001 | 0.107 (-0.059, 0.272) | 0.207 | 0.763 (0.669, 0.858) | <0.001 | 0.108 (-0.013, 0.229) | 0.079 | 0.139 (0.062, 0.215) | <0.001 | -0.015 (-0.090, 0.060) | 0.689 |
| | R | 3517 | 0.203 (0.074, 0.331) | 0.002 | 0.102 (-0.064, 0.268) | 0.229 | 0.749 (0.654, 0.844) | <0.001 | 0.092 (-0.025, 0.209) | 0.123 | 0.156 (0.078, 0.235) | <0.001 | -0.059 (-0.136, 0.019) | 0.137 |
| **Pallidum** | L | 3540 | 0.134 (0.036, 0.232) | 0.007 | 0.086 (-0.074, 0.247) | 0.293 | 0.750 (0.656, 0.845) | <0.001 | 0.060 (-0.034, 0.153) | 0.211 | 0.162 (0.094, 0.230) | <0.001 | -0.026 (-0.094, 0.042) | 0.460 |
| | R | 3525 | 0.186 (0.085, 0.287) | <0.001 | 0.072 (-0.093, 0.236) | 0.394 | 0.743 (0.648, 0.838) | <0.001 | 0.118 (0.026, 0.210) | 0.012 | 0.150 (0.074, 0.225) | <0.001 | -0.046 (-0.119, 0.026) | 0.210 |
| **Hippocampus** | L | 3552 | -0.018 (-0.130, 0.095) | 0.757 | 0.077 (-0.086, 0.240) | 0.356 | 0.778 (0.683, 0.873) | <0.001 | 0.042 (-0.068, 0.152) | 0.450 | 0.073 (0.002, 0.144) | 0.045 | -0.034 (-0.105, 0.037) | 0.347 |
| | R | 3551 | -0.053 (-0.164, 0.057) | 0.342 | 0.080 (-0.083, 0.244) | 0.335 | 0.771 (0.677, 0.866) | <0.001 | 0.088 (-0.016, 0.193) | 0.098 | 0.016 (-0.054, 0.086) | 0.654 | -0.018 (-0.084, 0.049) | 0.603 |
| **Amygdala** | L | 3540 | -0.074 (-0.178, 0.030) | 0.164 | 0.050 (-0.114, 0.213) | 0.551 | 0.770 (0.675, 0.864) | <0.001 | 0.014 (-0.089, 0.116) | 0.792 | 0.040 (-0.028, 0.107) | 0.248 | -0.016 (-0.084, 0.052) | 0.645 |
| | R | 3550 | -0.094 (-0.204, 0.016) | 0.095 | 0.057 (-0.105, 0.218) | 0.491 | 0.773 (0.679, 0.867) | <0.001 | -0.048 (-0.155, 0.059) | 0.377 | 0.063 (-0.006, 0.131) | 0.073 | -0.061 (-0.127, 0.005) | 0.072 |
| **Accumbens** | L | 3532 | 0.067 (-0.049, 0.183) | 0.256 | 0.078 (-0.085, 0.241) | 0.349 | 0.774 (0.679, 0.869) | <0.001 | 0.042 (-0.072, 0.156) | 0.470 | 0.114 (0.039, 0.188) | 0.003 | -0.037 (-0.112, 0.038) | 0.339 |
| | R | 3524 | 0.166 (0.046, 0.286) | 0.007 | 0.094 (-0.071, 0.259) | 0.264 | 0.771 (0.676, 0.866) | <0.001 | 0.062 (-0.062, 0.185) | 0.328 | 0.139 (0.063, 0.215) | <0.001 | -0.039 (-0.117, 0.039) | 0.331 |
| **Ventral DC** | L | 3557 | -0.239 (-0.349, -0.129) | <0.001 | 0.058 (-0.104, 0.221) | 0.483 | 0.805 (0.710, 0.899) | <0.001 | 0.026 (-0.086, 0.139) | 0.646 | -0.001 (-0.064, 0.062) | 0.979 | -0.004 (-0.069, 0.061) | 0.907 |
| | R | 3546 | -0.227 (-0.339, -0.114) | <0.001 | 0.063 (-0.100, 0.225) | 0.450 | 0.817 (0.722, 0.912) | <0.001 | -0.0006 (-0.111, 0.111) | 0.999 | -0.022 (-0.085, 0.041) | 0.497 | 0.00004 (-0.063, 0.063) | 0.999 |

**Note:** Estimated regression coefficients (β), 95% confidence intervals (CI) and *p*-values from the linear mixed-effects models evaluating the moderating effects of Oxwt in the relationship between brain microstructure and Body Mass Index. Models were controlled for age, education, race, ethnicity, puberty, co-exposure to other air pollutants, head motion during the MRI acquisition and urbanicity. Random intercepts were modelled from MRI scanner device and subject ID. Although all two-way interactions were assessed, this table presents only the ones of primary interest due to space constraints. Abbreviations: ROI: region-of-interest; Ventral DC: ventral diencephalon; RSI: restricted spectrum imaging; RND: restricted normalized directional diffusion; H: hemisphere. *p-values < 0.05 of two-way and three-way interactions that achieved the Benjamini-Hochberg False Discovery Rate threshold correction.
